# Multi-ancestry analysis of gene-sleep interactions in 126,926 individuals identifies multiple novel blood lipid loci that contribute to our understanding of sleep-associated adverse blood lipid profile

**DOI:** 10.1101/559393

**Authors:** Raymond Noordam, Maxime M Bos, Heming Wang, Thomas W Winkler, Amy R Bentley, Tuomas O. Kilpeläinen, Paul S de Vries, Yun Ju Sung, Karen Schwander, Brian E Cade, Alisa Manning, Hugues Aschard, Michael R Brown, Han Chen, Nora Franceschini, Solomon K Musani, Melissa Richard, Dina Vojinovic, Stella Aslibekyan, Traci M Bartz, Lisa de las Fuentes, Mary Feitosa, Andrea R Horimoto, Marjan Ilkov, Minjung Kho, Aldi Kraja, Changwei Li, Elise Lim, Yongmei Liu, Dennis O Mook-Kanamori, Tuomo Rankinen, Salman M Tajuddin, Ashley van der Spek, Zhe Wang, Jonathan Marten, Vincent Laville, Maris Alver, Evangelos Evangelou, Maria E Graff, Meian He, Brigitte Kühnel, Leo-Pekka Lyytikäinen, Pedro Marques-Vidal, Ilja M Nolte, Nicholette D Palmer, Rainer Rauramaa, Xiao-Ou Shu, Harold Snieder, Stefan Weiss, Wanqing Wen, Lisa R Yanek, Correa Adolfo, Christie Ballantyne, Larry Bielak, Nienke R Biermasz, Eric Boerwinkle, Niki Dimou, Gudny Eiriksdottir, Chuan Gao, Sina A Gharib, Daniel J Gottlieb, José Haba-Rubio, Tamara B Harris, Sami Heikkinen, Raphaël Heinzer, James E Hixson, Georg Homuth, M Arfan Ikram, Pirjo Komulainen, Jose E Krieger, Jiwon Lee, Jingmin Liu, Kurt K Lohman, Annemarie I Luik, Reedik Mägi, Lisa W Martin, Thomas Meitinger, Andres Metspalu, Yuri Milaneschi, Mike A Nalls, Jeff O’Connell, Annette Peters, Patricia Peyser, Olli T Raitakari, Alex P Reiner, Patrick CN Rensen, Treva K Rice, Stephen S Rich, Till Roenneberg, Jerome I Rotter, Pamela J Schreiner, James Shikany, Stephen S Sidney, Mario Sims, Colleen M Sitlani, Tamar Sofer, Konstantin Strauch, Morris A Swertz, Kent D Taylor, André G Uitterlinden, Cornelia M van Duijn, Henry Völzke, Melanie Waldenberger, Robert B Wallance, Ko Willems van Dijk, Caizheng Yu, Alan B Zonderman, Diane M Becker, Paul Elliott, Tõnu Esko, Christian Gieger, Hans J Grabe, Timo A Lakka, Terho Lehtimäki, Lifelines Cohort Study, Kari E North, Brenda WJH Penninx, Peter Vollenweider, Lynne E Wagenknecht, Tangchun Wu, Yong-Bing Xiang, Wei Zheng, Donna K Arnett, Claude Bouchard, Michele K Evans, Vilmundur Gudnason, Sharon Kardia, Tanika N Kelly, Stephen B Kritchevsky, Ruth JF Loos, Alexandre C Pereira, Mike Province, Bruce M Psaty, Charles Rotimi, Xiaofeng Zhu, Najaf Amin, L Adrienne Cupples, Myriam Fornage, Ervin F Fox, Xiuqing Guo, W James Gauderman, Kenneth Rice, Charles Kooperberg, Patricia B Munroe, Ching-Ti Liu, Alanna C Morrison, Dabeeru C Rao, Diana van Heemst, Susan Redline

## Abstract

Both short and long sleep are associated with an adverse lipid profile, likely through different biological pathways. To provide new insights in the biology of sleep-associated adverse lipid profile, we conducted multi-ancestry genome-wide sleep-SNP interaction analyses on three lipid traits (HDL-c, LDL-c and triglycerides). In the total study sample (discovery + replication) of 126,926 individuals from 5 different ancestry groups, when considering either long or short total sleep time interactions in joint analyses, we identified 49 novel lipid loci, and 10 additional novel lipid loci in a restricted sample of European-ancestry cohorts. In addition, we identified new gene-sleep interactions for known lipid loci such as *LPL* and *PCSK9*. The novel gene-sleep interactions had a modest explained variance in lipid levels: most notable, gene-short-sleep interactions explained 4.25% of the variance in triglyceride concentration. Collectively, these findings contribute to our understanding of the biological mechanisms involved in sleep-associated adverse lipid profiles.

## Introduction

Dyslipidemia is defined as abnormalities in one or more types of lipids, such as high blood LDL-cholesterol (LDL-c) and triglyceride (TG) concentrations and a low HDL-cholesterol (HDL-c) concentration. High LDL-c and TG are well-established modifiable causal risk factors for cardiovascular disease ^1-3^, and therefore are a primary focus for preventive and therapeutic interventions. Over 300 genetic loci have been identified to be associated with blood lipid concentrations ^4-10^. Recent studies showed that only 12.3% of the total variance in lipid concentration is explained by common single nucleotide polymorphisms (SNPs), suggesting additional lipid loci could be uncovered ^10^. Some of the unexplained heritability may be due to the presence of gene-environment and gene-gene interactions. Recently, high levels of physical activity were shown to modify the effects of four genetic loci on lipid levels ^11^, an additional 18 novel lipid loci were identified when considering interactions with high alcohol consumption ^12^, and 13 novel lipid loci were identified when considering interaction with smoking status ^13^, suggesting that behavioral factors may interact with genetic loci to influence lipid levels.

Sleep is increasingly recognized as a fundamental behavior that influences a wide range of physiological processes ^14^. A large volume of epidemiological research implicates disturbed sleep in the pathogenesis of atherosclerosis ^15^, and specifically, both a long and short sleep duration are associated with an adverse blood lipid profile ^16-26^. However, it is unknown whether sleep duration modifies genetic risk factors for adverse blood lipid profiles. We hypothesize that short and long habitual sleep duration may modify genetic associations with blood lipid levels. The identification of SNPs involved in such interactions will facilitate our understanding of the biological background of sleep-associated adverse lipid profiles.

We investigated gene-sleep duration interaction effects on blood lipid levels as part of the Gene-Lifestyle Interactions Working Group within the Cohorts for Heart and Aging Research in Genomic Epidemiology (CHARGE) Consortium ^27,28^. To permit the detection of both such sleep-duration-SNP interactions and lipid-SNP associations accounting for total sleep duration, a 2 degree of freedom (2df) test that jointly tests the SNP main effect and the interaction effect was applied ^29^. Given that there are differences among ancestry groups in sleep behaviours and lipid levels, analysis of data from cohorts of varying ancestries may further the discovery of robust interactions between genetic loci and sleep traits. We focused on short total sleep time (STST; defined as the lower 20% of age- and sex-adjusted sleep duration residuals) and long total sleep time (LTST; defined as the upper 20% of age- and sex-adjusted sleep duration residuals) as exposures compared to the remaining individuals in the study population, given that each extreme sleep trait has been associated with multiple metabolic and health outcomes ^15-26,30-34^. Within the present study, we report multi-ancestry sleep-by-SNP interaction analyses for blood lipid levels that successfully identified several novel loci for blood lipid traits.

## Results

### Study population

Discovery analyses were performed in up to 62,457 individuals (40,041 European-ancestry, 14,908 African-ancestry, 4,460 Hispanic-ancestry, 2,379 Asian-ancestry, and 669 Brazilian/mixed-ancestry individuals) from 21 studies spanning 5 different ancestry groups (**Supplementary Tables 1-3**). Of the total discovery analysis, 13,046 (20.9%) individuals were classified as short sleepers and 12,317 (19.7%) individuals as long sleepers. Replication analyses were performed in up to 64,469 individuals (47,612 European-ancestry, 12,578 Hispanic-ancestry, 3,133 Asian-ancestry, and 1,146 African-ancestry individuals) from 19 studies spanning 4 different ancestry groups (**Supplementary Tables 4-6**). Of the total replication analysis, 12,952 (20.1%) individuals were classified as short sleepers and 12,834 (19.9%) individuals as long sleepers.

### Identification of novel loci for lipid traits when considering potential interaction with long or short total sleep time

An overview of the multi-ancestry analyses process for both STST and LTST is presented in **Figure 1**. In the combined discovery and replication meta-analyses comprising all contributing ancestry groups, we found that many SNPs replicated for the lipid traits (P_joint_ in replication < 0.05 with similar directions of effect as in the discovery analyses and P_joint_ in combined discovery and replication analysis < 5×10^-8^). Notably, we replicated 2,395 and 2,576 SNPs for HDL-c, 2,012 and 2,074 SNPs for LDL-c, and 2,643 and 2,734 SNPs for TG in the joint model with LTST and STST respectively.

**Figure 1:**
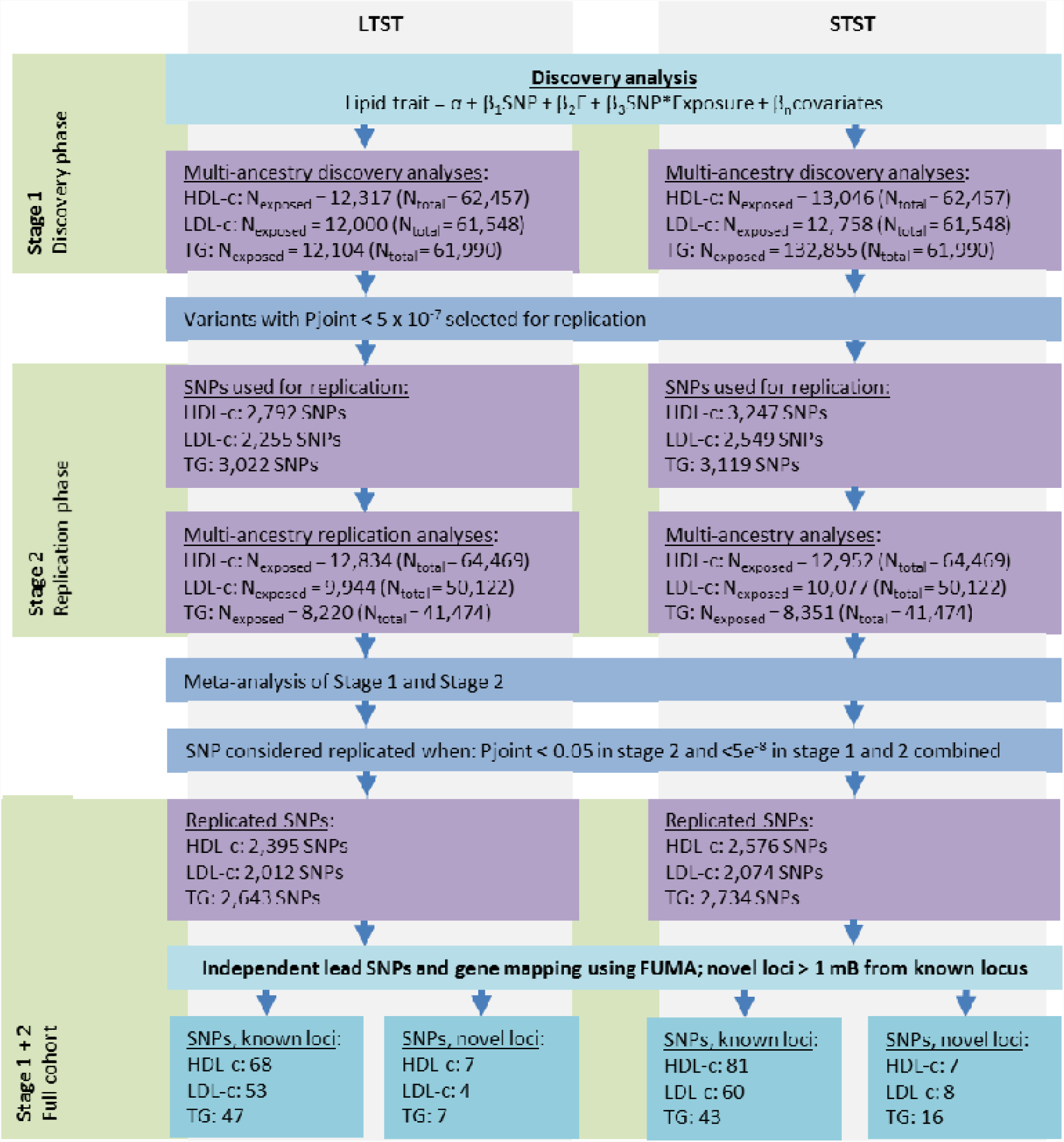
Project overview and SNP selection in the multi-ancestry analyses. Project overview of the multi-ancestry analyses of how the new lipid loci were identified in the present project. Replicated variants had to have a P_joint_ < 5×10^-7^ in stage 1, P_joint_ < 0.05 with similar direction of effect in stage 2, and P_joint_ <5×10^-8^ in the combined stage 1 + 2.

Most of the replicated SNPs were mapped to known loci (**Supplementary Tables 7 and 8**). In addition, we identified lead SNPs mapping to novel regions when considering the joint model with potential interaction for either STST or LTST (>1 Mb distance from known locus). Ultimately, in the multi-ancestry analysis, we identified 14 novel loci for HDL-c, 12 novel loci for LDL-c, and 23 novel loci for TG (R^2^ < 0.1; **Figure 2**). Of these, 7 loci for HDL-c, 4 loci for LDL-c and 7 loci for TG were identified after considering an interaction with LTST (**Supplementary Table 9**). Furthermore, 7 loci for HDL-c, 8 loci for LDL-c and 16 loci for TG were identified when considering an interaction with STST (**Supplementary Table 10**). Importantly, none of the novel loci for the three lipid traits identified through LTST were identified in the analyses with STST, and *vice versa*. Furthermore, the novel lipid loci were specific to a single lipid trait. Regional plots of the newly identified loci from the multi-ancestry analyses are presented in **Supplementary Figures 1-6**. Using the R-based VarExp package^35^, we calculated the explained variance based on the summary statistics of the combined discovery and replication analysis. Collectively, novel lead SNPs identified with LTST explained 0.97% of the total HDL-c variation, 0.13% of the total LDL-c variation, and 1.51% of the total TG variation. In addition, novel lead SNPs identified with STST explained 1.00% of the total HDL-c variation, 0.38% of the total LDL-c variation, and 4.25% of the total TG variation.

**Figure 2:**
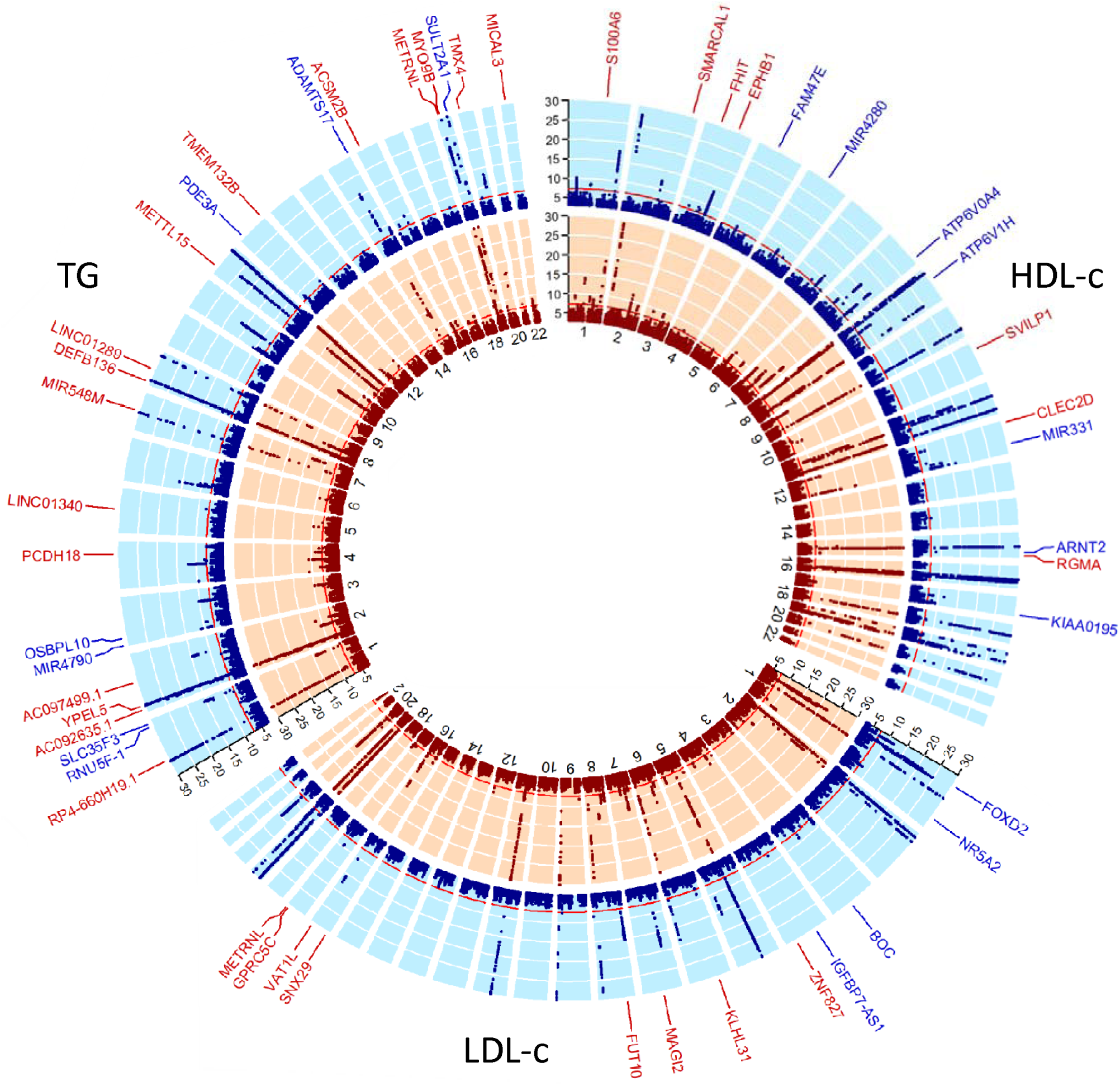
Circular –log(p-value) plots of the multi-ancestry sleep-SNP interactions analyses for the three lipid traits. Plot visualizes the –log(p_joints_) for HDL-c, LDL-c and TG per chromosome. In red (inner circle) are the –log(p-value) plots for the analyses taking into account potential interaction with short total sleep time. In blue (outer circle) are the –log(p-value plots for the analyses taking into account potential interaction with long total sleep time. Loci defined as novel and replicated are labeled. Replicated variants had to have a P_joint_ < 5×10^-7^ in stage 1, P_joint_ < 0.05 in stage 2, and P_joint_ <5×10^-8^ in the combined stage 1 + 2. Labeled gene names in red were identified in the STST analysis; Labeled gene names in blue were identified in the LTST analysis. All –log(p_joints_) > 30 were truncated to 30 for visualization purposes only. The unlabeled regions with P_joint_ < 5×10^-8^ were in known loci. Figure prepared using the R package circlize^100^.

In the analyses restricted to European-ancestry individuals (overview **Supplementary Figure 7)**, we identified 10 additional novel loci (7 novel loci with LTST and 3 novel loci with STST; **Supplementary Figure 8**), which were not identified in the multi-ancestry analyses. Of these, we identified 4 loci for HDL-c, 2 loci for LDL-c, and 1 locus for TG with LTST (**Supplementary Table 11**). In addition, we identified 1 locus for HDL-c and 2 for TG with STST (**Supplementary Table 12**). Again, we observed no overlapping findings between the two sleep exposures and the three lipid traits. Regional plots of the identified novel loci were presented in **Supplementary Figures 9-13**.

### Sleep × SNP interactions in identified novel and known lipid loci in the combined sample of discovery and replication studies

Based on a total of 402 novel and known lead SNPs for both exposures and the three lipid traits that were identified using the joint test in the combined sample of discovery and replication studies, we subsequently explored the extent to which the effects were driven by interaction with the sleep exposure trait being tested (using to a 1df interaction test ^29^). We corrected the 1df interaction p-value for multiple testing using the false-discovery rate ^36^ considering all 402 lead SNPs for the present investigation, which was equivalent in our study to a 1df interaction p-value <5×10^-4^. Overall, in the multi-ancestry meta-analyses, the novel lipid loci show clearly stronger interaction with either LTST or STST than the loci defined as known (**Figure 3**). The majority of the newly identified lead variants were generally common, with minor allele frequencies (MAF) mostly > 0.2, and SNP × sleep interaction effects were not specifically identified in lower frequency SNPs (e.g., MAF<0.05).

**Figure 3:**
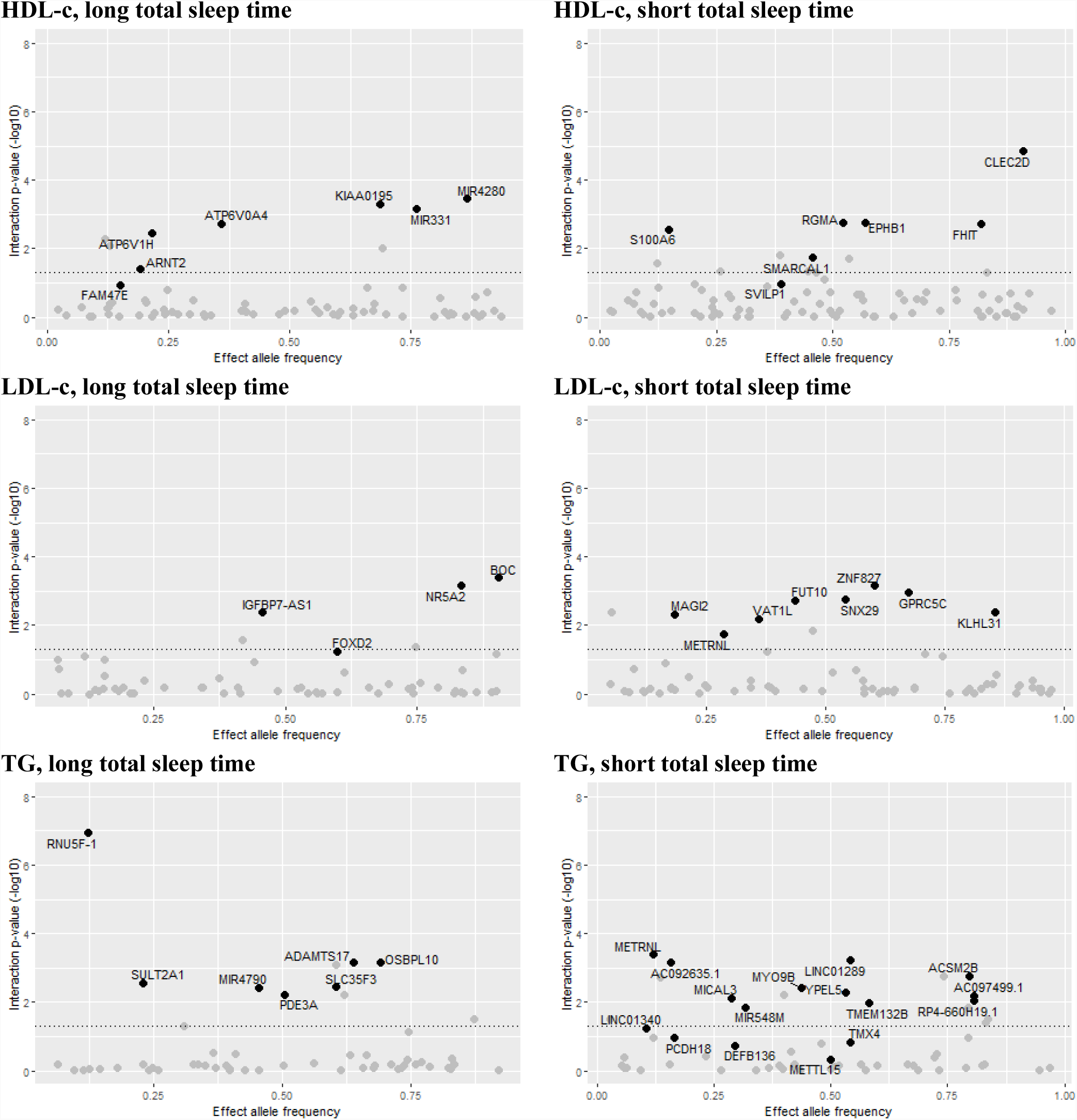
Sleep-interactions on lipid traits in novel and known loci in the multi-ancestry meta-analyses. Plot displaying the –log(p-value) of the 1df interaction between the SNP and either LTST or STST on the lipid trait after correction for multiple testing using false-discovery rate against the allele frequency of the effect allele. Dotted horizontal line resembles the cut-off for the 1df interaction p-value_FDR_ <0.05 after correction for multiple testing using false-discovery rate. In black are the novel loci for lipid traits; in grey are the identified lead SNPs mapped within a 1 Mb physical distance from a known lipid locus. Visualization of the plots was performed using the R package ggplot2 ^101^.

Out of the 7 novel HDL-c loci identified in the joint model with LTST, 6 had a 1df interaction p-value_FDR_ < 0.05, notably lead SNPs mapped to *ATP6V1H, ARTN2, ATP6V0A4, KIAA0195, MIR331*, and *MIR4280.* Based on exposure-stratified analyses in the discovery cohorts, we further explored the effect sizes per exposure group. The lead SNPs that showed significant sleep × SNP interaction also showed effect estimates that modestly differed between LTST exposure groups (**Supplementary Table 13**). Interestingly, two lead SNPs near known HDL-c loci showed a 1df interaction p-value_FDR_ < 0.05, including SNPs near *CETP* and *LIPC* (**Supplementary Table 7**). Out of the 7 novel HDL-c loci identified in the joint model with STST, we found 6 loci with a 1df interaction p-value_FDR_ < 0.05, notably lead SNPs mapped to *S1000A6, SMARCAL1, RGMA, EPHB1, FHIT* and *CLEC2D.* Again, their effect estimates differed between the exposure groups in the discovery multi-ancestry meta-analysis (**Supplementary Table 14; Figure 4**). Some lead SNPs near known HDL-c loci showed evidence of a 1df interaction with STST (e.g., *MADD* and *LPL;* p-value_FDR_ < 0.05).

**Figure 4:**
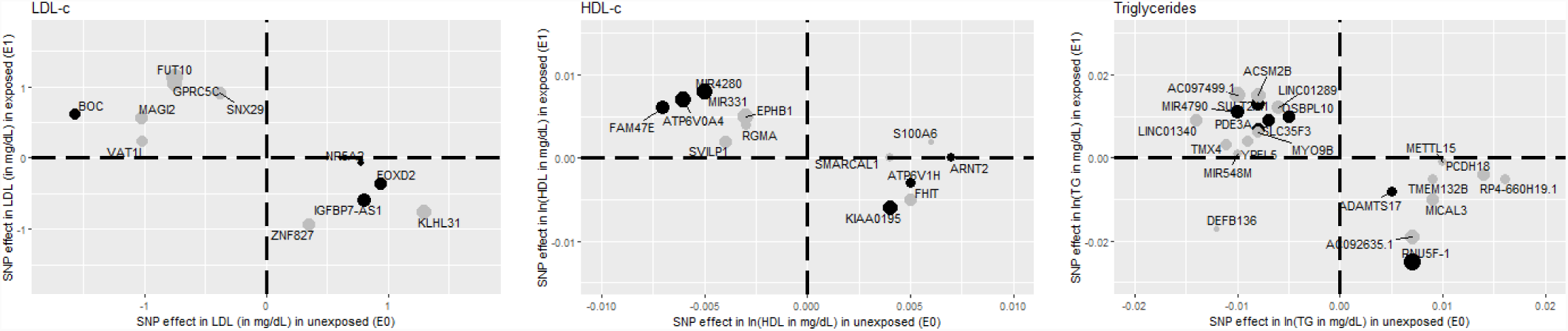
Comparison of SNP-main effects stratified by exposure in the multi-ancestry discovery meta-analyses. X-axis displays the effect sizes of the novel lead SNPs as observed in the meta-analyses of the unexposed individuals (LTST = “0”, STST = “0”). Y-axis displays the effect sizes of the novel lead SNPs as observed in the meta-analyses of the exposed individuals (LTST = “1”, STST = “1”). In black are the novel lead SNPs identified with LTST; in grey are the novel lead SNPs identified with STST. Sizes of the dots were weighted to the difference observed between exposed and unexposed. Visualization of the plots was performed using the R package ggplot2 ^101^.

For all four novel lead SNPs associated with LDL-c when considering LTST, we observed a 1df interaction p-value_FDR_ < 0.05; notably, lead SNPs mapped to *IGFBP7-AS1, FOXD2, NR5A2* and *BOC*. One locus that mapped within a 1 Mb physical distance from known LDL-c locus (*PCSK9*) showed 1df interaction with LTST (**Supplementary Table 7**). Similarly, all 8 independent novel lead SNPs associated with LDL-c when considering STST, had a 1df interaction p-value_FDR_ <0.05; notably, lead SNPs mapped to *MAGI2, METRNL, VAT1L, FUT10, SNX29, ZNF827, GPRC5C* and *KLHL31*. In addition, of the known LDL-c loci, lead SNPs mapped within a physical distance of 1 Mb of *APOB* and *SLC22A1* showed a 1df interaction p-value_FDR_ < 0.05 (**Supplementary Table 8**). For both analyses, we observed that effect estimates differed between the LTST and STST exposure groups in the multi-ancestry discovery analysis (**Supplementary Table 13 and 14; Figure 4**).

All 7 independent novel lead SNPs associated with TG when considering LTST, had a 1df interaction p-value_FDR_ <0.05; notably, lead SNPs mapped to *RNU5F*-1, *SULT2A1, MIR4790, PDE3A, SLC35F3, ADAMTS17* and *OSBPL10*. In addition, we found some evidence for long sleep-SNP interaction in lead SNPs near known TG loci, including lead SNPs near *AKR1C4* and *NAT2* (**Supplementary Table 7**). Of the 16 novel lead SNPs associated with TG when considering STST, we observed 12 lead SNPs with a 1df interaction p-value <5×10^-4^ (p-value_FDR_ < 0.05), including lead SNPs mapped to *LINC0140, METRNL, AC092635.1, MICAL3, MIR548M, MYO9B, YPEL5, LINC01289, TMEM132B, ACSM2B, AC097499.1* and *RP4-660H19.1.* In addition, we observed some lead SNPs within 1 Mb physical distance from known TG loci, such as *MMP3* and *NECTIN2* (**Supplementary Table 8**). For both LTST and STST analyses, we again observed differing effects dependent on the exposure group in the discovery meta-analyses (**Supplementary Table 13 and 14; Figure 4**).

### Look-ups and bioinformatics analyses

Based on the lead SNPs mapped to novel loci, we conducted a look-up in GWAS summary statistics data on different questionnaire-based sleep phenotypes from up to 337,074 European-ancestry individuals of the UK Biobank (**Supplementary Table 15**). We only observed the TG-identified rs7924896 (*METTL15*) to be associated with snoring (p-value = 1e^-5^) after correction for a total of 343 explored SNP-sleep associations (7 sleep phenotypes × 49 genes; 10 SNPs were unavailable; threshold for significance = 1.46e^-4^). In general, we did not find substantial evidence that the identified novel lead SNPs were associated with coronary artery disease in the CARGIoGRAMplusC4D consortium (**Supplementary Table 16**).

Newly identified lipid loci were further explored in the GWAS catalogue (**Supplementary Table 17**). Several of the mapped genes of our novel lead SNPs have previously been identified with multiple other traits, such as body mass index (*FHIT, KLH31, ADAMTS17, MAGI2*), mental health (*FHIT* [autism/schizophrenia, depression], *SNX13* [cognition]), gamma-glutamyltransferase (*ZNF827, MICAL3*), and inflammatory processes (*ZNF827, NR5A2*).

We additionally investigated differential expression of the novel lead SNPs using data from multiple tissues from the GTEx consortium ^37,38^ (**Supplementary Table 18**). Lead SNPs were frequently associated with mRNA expression levels of the mapped gene and with trans-eQTLs. For example, rs429921 (mapped to *VAT1L*) was associated with differential mRNA expression levels of *CLEC3A* and *WWOX,* which are located more upstream on chromosome 16 (**Supplementary Figure 4**). rs3826692 (mapped to *MYO9B*) was specifically associated with differential expression of the nearby *USE1* gene. Identified SNPs were frequently associated with differential expression in the arteries. For example, rs6501801 (*KIAA0195*) was associated with differential expression in arteries at different locations. Several of the other identified SNPs showed differential expression in multiple tissues, including the gastrointestinal tract, (subcutaneous/visceral) adipose tissue, brain, heart, muscle, lung, liver, nervous system, skin, spleen, testis, thyroid and whole blood.

## Discussion

We investigated SNP-sleep interactions in a large, multi-ancestry, meta-analysis of blood lipid levels. Given the growing evidence that sleep influences metabolism ^39-44^, at least in part through effects on gene expression, we hypothesized that short/long habitual sleep duration may modify the effects of genetic loci on lipid levels. In a total study population of 126,926 individuals from 5 different ancestry groups, we identified 49 novel lipid loci when considering either long or short total sleep time in the analyses. An additional 10 novel lipid loci were identified in analyses in Europeans only. Of the newly identified loci, most loci at least in part driven by differing effects in short/long sleepers compared to the rest of the study population, which was reflected by significant FDR-corrected 1df interactions. Multiple of the novel genes identified by our efforts have been previously identified in relation to adiposity, hepatic function, inflammation or psychosocial traits, collectively contributing to potential biological mechanisms involved in sleep-associated adverse lipid profile.

In addition to the over 300 genetic loci that already have been identified in relation to blood lipid concentrations in different efforts ^4-10^, we identified 49 additional loci associated with either HDL-c, LDL-c or TG in our multi-ancestry analysis. Considering the novel TG loci identified by considering interactions with total sleep duration explain an additional 4.25% and 1.51% of the total variation in TG concentrations, for STST and LTST, respectively. While the additionally explained variance for LDL-c (0.38% and 0.13%) and HDL-c (1.00% and 0.97%) was low/modest, the novel lead SNPs identified include genes known to be associated with adiposity, inflammatory disorders, cognition, and liver function, thus identifying pathways by which sleep disturbances may influence lipid biology.

Across multiple populations, both short and long sleep duration have been associated with cardiovascular disease and diabetes ^45^. There are numerous likely mechanisms for these associations. Experimental sleep loss results in inflammation, cellular stress in brain and peripheral tissues, and altered expression of genes associated with oxidative stress ^46,47^. The impact of long sleep on metabolism is less well understood than the effect of short sleep, and multiple of the associations seem to overlap with short sleep as well. Long sleep duration is associated with decreased energy expenditure, increased sedentary time, depressed mood, and obesity-related factors associated with inflammation and a pro-thrombotic state ^48^, as well as with higher C-reactive protein and interleukin-6 concentrations ^49^. However, studies that adjusted for multiple confounders, including obesity, depression, and physical activity, showed that long sleep remained a significant predictor of adverse cardiovascular outcomes ^45,50^. Therefore, the adverse effects of long sleep also may partly reflect altered sleep-wake rhythms and chronodisruption resulting from misalignment between the internal biological clock with timing of sleep and other behaviours that track with sleep, such as timing of food intake, activity, and light exposure ^51^. Altered sleep-wake and circadian rhythms influence glucocorticoid signalling and autonomic nervous system excitation patterns across the day ^41^, which can influence the phase of gene expression. These inputs appear to be particularly relevant for genes controlling lipid biosynthesis, absorption and degradation, many of which are rhythmically regulated and under circadian control ^52^. Moreover, the molecular circadian clock acts as a rate limiting step in cholesterol and bile synthesis, supporting the potential importance of circadian disruption in lipid biology ^53^. Collectively, these data suggest different biological mechanisms involved in short and long sleep-associated adverse lipid profiles.

Consistent with different hypothesized physiological effects of short and long sleep, we observed no overlap in the novel loci that were identified by modelling interactions with short or long sleep duration. Novel lipid loci that were identified after considering STST include *FHIT, MAGI2* and *KLH3,* which have been previously associated with body mass index (BMI) ^54-60^. Interestingly, although not genome-wide significant, variation in *MAGI2* has been associated with sleep duration ^61^, however, we did not find evidence for an association with rs10244093 in *MAGI2* with any sleep phenotype in the UK Biobank sample. Variants in *MICAL3* and *ZNF827,* that were also identified after considering STST, have been associated with serum liver enzymes gamma-glutamyl transferase measurement and/or aspartate aminotransferase levels ^62,63^, which have been implicated in cardiometabolic disturbances ^64-67^ and associated with prolonged work hours (which often results in short or irregular sleep) ^68^. Other loci identified through interactions with STST were in genes previously associated with neurocognitive and neuropsychiatric conditions, possibly reflecting associations mediated by heightened levels of cortisol and sympathetic activity that frequently accompany short sleep.

In relation to LTST, the novel genes identified have been previously related to inflammation-driven diseases of the intestine, blood pressure and blood count measurements, including traits influenced by circadian rhythms ^69,70^. However, no novel identified lipid loci with LTST directly interacted with genes involved in the central circadian clock (e.g., *PER2, CRY2* and *CLOCK*) in the KEGG pathways database ^71^. The novel loci *NR5A2* and *SLC35F3* have been associated with inflammation-driven diseases of the intestine ^72,73^. Ulcerative colitis, an inflammatory bowel disease, has been associated with both longer sleep duration ^74^ and circadian disruption ^69^. *ARNT2*, also identified via a LTST interaction, heterodimerizes with transcriptional factors implicated in homeostasis and environmental stress responses ^75,76^. A linkage association study has reported nominal association of this gene with lipids in a Caribbean Hispanic population ^77^.

We identified a number of additional genetic lead SNPs in the meta-analyses performed in European-Americans only. For example, we identified rs3938236 mapped to *SPRED1* to be associated with HDL-c after accounting for potential interaction with LTST. Interestingly, this gene has been previously associated with hypersomnia in Caucasian and Japanese populations ^*78*^, but was not identified in our larger multi-ancestry analysis, possibly due to cultural differences in sleep behaviours ^79^.

We additionally found evidence, amongst others, in the known lipid loci *APOB, PCSK9* and *LPL* for interaction with either short or long sleep. Associations have been observed previously between short sleep and ApoB concentrations, have been observed previously ^80^. LPL expression has been shown to follows a diurnal rhythm in several metabolic organs ^43,81^, and disturbing sleeping pattern by altered light exposure can lower LPL activity, at least in brown adipose tissue ^43^. Similar effects of sleep on hepatic secretion of ApoB and PCSK9 may be expected. Indeed, in humans PCSK9 has a diurnal rhythm synchronous with hepatic cholesterol synthesis ^82^. Although the interaction effects we observed were rather weak, the supporting evidence from the literature suggests that sleep potentially modifies the effect of some of the well-known lipid regulators that are also targets for therapeutic interventions.

Some of the novel lipid loci have been previously associated with traits related to sleep. For example, *MAGI2* and *MYO9B* ^61^ have been suggestively associated with sleep duration and quality, respectively. Genetic variation in *TMEM132B* has been associated with excessive daytime sleepiness ^83^, and *EPHB1* has been associated with self-reported chronotype ^84^. These findings suggest some shared genetic component of lipid regulation and sleep biology. However, with the exception of the *METTL15*-mapped rs7924896 variant in relation to snoring, none of the lead SNPs mapped to the new lipid loci were associated with any of the investigated sleep phenotypes in the UK Biobank population, suggesting no or minimal shared component in sleep and lipid biology but rather that sleep duration specifically modifies the effect of the variant on the lipid traits.

The present study was predominantly comprised of individuals of European ancestry, despite our efforts to include as many studies of diverse ancestries as possible. For this reason, additional efforts are required to specifically study gene-sleep interactions in those of African, Asian and Hispanic ancestry once more data becomes available. In line, we identified several loci that were identified only in the European-ancestry analysis, and not in the multi-ancestry analysis, suggestion that there might be ancestry-specific effects. The multi-ancestry analysis highlighted the genetic regions that are more likely to play a role in sleep-associated adverse lipid profiles across ancestries. In addition, our study used questionnaire-based data on sleep duration. Although the use of questionnaires likely increased measurement error and decreased statistical power, questionnaire-based assessments of sleep duration have provided important epidemiological data, including the identification of genetic variants for sleep traits in genome-wide association studies^83^. Identified variants for sleep traits have been recently successfully validated using accelerometer data ^85^. At this time, we did not have sufficient data to evaluate other measures of sleep duration such as polysomnography or accelerometery; however, a more comprehensive characterization and circadian traits likely will refine our understanding of the interaction of these fundamental phenotypes and lipid biology.

In summary, the gene-sleep interaction efforts described in the present multi-ancestry study identified many novel genetic loci associated with either HDL-c, LDL-c or triglycerides levels. Multiple of the novel genetic loci were driven by interactions with either short or long sleep duration, and were mapped to genes also associated with adiposity, inflammatory or neuropsychiatric traits. Collectively, the results highlight the interactions between extreme sleep-wake exposures and lipid biology.

## Online methods

Details regarding motivation and methodology of this and other projects of the CHARGE Gene-Lifestyle Interactions Working Group are available elsewhere^86^.

### Participants

Analyses were performed locally by the different participating studies. Discovery and replication analyses comprised men and women between the age of 18 and 80 years, and were conducted separately for the different contributing ancestry groups, including: European, African, Asian, Hispanic, and Brazilian (discovery analysis only). Descriptions of the different participating studies are described in detail in the **Supplementary Materials**, and study-specific characteristics (sizes, trait distribution and data preparation) are presented in **Supplementary Tables 1-6**. Every effort was made to include as many studies as possible. The present work was approved by the Institutional Review Board of Washington University in St. Louis and complies with all relevant ethical regulations. Each participating study obtained written informed consent from all participants and received approval from the appropriate local institutional review boards.

### Lipid traits

We conducted all analyses on the following lipid traits: HDL-c, LDL-c, and TG. TG and LDL-c concentrations were measured in samples from individuals who had fasted for at least 8 hours. LDL-c could be either directly assayed or derived using the Friedewald equation^87^ (the latter being restricted to those with TG ≤ 400 mg/dL). We furthermore corrected LDL-c for the use of lipid-lowering drugs, defined as any use of a statin drug or any unspecified lipid-lowering drug after the year 1994 (when statin use became common in general practice). If LDL-c was directly assayed, the concentration of LDL-c was corrected by dividing the LDL-c concentration by 0.7. If LDL-c was derived using the Friedewald equation, we first divided the concentration of total cholesterol by 0.8 before LDL-c was calculated by the Friedewald equation. Due to the skewed distribution of HDL-c and TG, we ln-transformed the concentration prior to the analyses; no transformation for LDL-c was required. When an individual cohort measured the lipid traits during multiple visits, the visit with the largest available sample and concurrent availability of the sleep questions was selected.

### Nocturnal total sleep time

Contributing cohorts collected information on the habitual sleep duration using either a single question such as “on an average night, how long do you sleep?” or as part of a standardized sleep questionnaire (e.g., the Pittsburgh Sleep Quality Index questionnaire^88^). For the present project, we defined both STST and LTST. To harmonize the sleep duration data across cohorts and to minimize misclassification of total sleep duration caused by any differences in reporting by age or sex, we calculated residuals of total sleep time adjusted for age and sex. Exposure to STST was defined as the lowest 20% of the sex- and age-adjusted sleep-time residuals (coded as “1”). Exposure to LTST was defined as the highest 20% of the sex- and age-adjusted sleep-time residuals (coded as “1”). For both sleep-time definitions, we considered the remaining 80% of the population as being unexposed to either STST or LTST (coded as “0”). In some cases, we used a cohort-specific definition in case data were not suitable (e.g., multiple-choice question for total sleep time with limited number of alternatives) for the calculation of sex- and age-adjusted residuals. In these studies, no residuals were calculated, but instead, we defined STST or LTST based on the alternatives available in the multiple-choice question after consultation of a sleep expert.

### Genotype Data

Genotyping was performed by each participating study locally using genotyping arrays from either Illumina (San Diego, CA, USA) or Affymetrix (Santa Clara, CA, USA). Each study conducted imputation using various software programs. The cosmopolitan reference panel from the 1000 Genomes Project Phase I Integrated Release Version 3 Haplotypes (2010-11 data freeze, 2012-03-14 haplotypes) was specified for imputation. Only SNPs on the autosomal chromosomes with a minor allele frequency of at least 0.01 were considered in the analyses. Specific details of each participating study’s genotyping platform and imputation software are described (**Supplementary Tables 3** and **6)**.

### Stage 1 Analysis (discovery phase)

The discovery phase of the present project included 21 cohorts contributing data from 28 study/ancestry groups, and included up to 62,457 participants of EUR, AFR, ASN, HISP and BR ancestry (**Supplementary Tables 1-3**). All cohorts ran statistical models according to a standardized analysis protocol. The main model for this project examined the SNP-main effect and the multiplicative interaction term between the SNP and either LTST or STST:

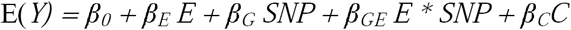

In which E is the sleep exposure variable (LTST/STST) and C are the (study-specific) covariates, which was similar to what we have done in previous studies ^4,11,12^. In addition, we examined the SNP-main effect (without incorporating LTST/STST) and the SNP-main effect stratified by the exposure:

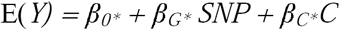

All models were performed for each lipid trait and separately for the different ancestry groups. Consequently, per ancestry group, we requested a total of 7 GWA analyses per lipid trait. All models were adjusted for age, sex, field center (if required), and the first principal components to correct for population stratification. The number of principal components included in the model was chosen according to cohort-specific preferences (ranging from 0 to 10). All studies were asked to provide the effect estimates (SNP-main and -interaction effect) with accompanying robust estimates of the standard error for all requested models. A robust estimate of the covariance between the main and interaction effects was also provided. To obtain robust estimates of covariance matrices and standard errors, studies with unrelated participants used R packages such as either sandwich^89^ or ProbABEL^90^. Studies including related individuals used either generalized estimating equations (R package geepack^91^) or linear mixed models (GenABEL^92^, MMAP, or R) were used. Sample code provided to studies to generate these data has been previously published (see **Supplementary Materials** ^86^).

Upon completion of the analyses by local institution, all summary data were stored centrally for further processing and meta-analyses. We performed estimative quality control (QC) using the R-based package EasyQC^93^ (www.genepi-regensburg.de/easyqc) at the study level (examining the results of each study individually), and subsequently at the ancestry level (after combining all ancestry-specific cohorts using meta-analyses). Study-level QC consisted of excluding all SNPs with MAF < 0.01, harmonization of alleles, comparison of allele frequencies with ancestry-appropriate 1000 Genomes reference data, and harmonization of all SNPids to a standardized nomenclature according to chromosome and position. Ancestry-level QC included the compilation of summary statistics on all effect estimates, standard errors and p-values across studies to identify potential outliers, and production of SE-N and QQ plots to identify analytical problems (such as improper trait transformations)^94^.

Prior to the ancestry-specific meta-analyses, we excluded the following SNPs from the cohort-level data files: all SNPs with an imputation quality < 0.5, and all SNPs with a minor allele count in the exposed group (LTST or STST equals “1”) x imputation quality of less than 20. SNPs in the European-ancestry and multi-ancestry analyses had to be present in at least 3 cohorts and 5000 participants. Due to the limited sample size of the non-European ancestries (either discovery or replication), we did not take into account this filter in those ancestry-level meta-analyses.

Meta-analyses were conducted for all models using the inverse variance-weighted fixed effects method as implemented in METAL^95^ (http://genome.sph.umich.edu/wiki/METAL). We evaluated both a 1 degree (1df) of freedom test of interaction effect and a 2 degree of freedom (2df) joint test of main and interaction effects, following previously published methods^29^. A 1df Wald test was used to evaluate the 1df interaction, as well as the main effect in models without an interaction term. A 2df Chi-squared test was used to jointly test the effects of both the variant and the variant x LTST/STST interaction^96^. Meta-analyses were conducted within each ancestry separately. Multi-ancestry meta-analyses were conducted on all ancestry-specific meta-analyses. Genomic control correction was applied on all cohorts incorporated in the ancestry-level meta-analyses as well as on the final meta-analyses for the publication. From this effort, we selected all SNPs associated with any of the lipid traits with p ≤ 5×10^-7^ for replication in the Stage 2 analysis.

### Stage 2 Analysis (replication phase)

All SNPs selected in Stage 1 for replication were evaluated in the interaction model in up to 18 cohorts contributing data from 20 study groups totalling up to 64,469 individuals (**Supplementary Tables 4-6**). As we had a limited number of individuals from non-European ancestry in the replication analyses, we did not consider an the non-European ancestries separately and only focussed on a European-ancestry and multi-ancestry analysis.

Study- and ancestry-level QC was carried out as in stage 1. In contrast to stage 1, no additional filters were included for the number of studies or individuals contributing data to stage 2 meta-analyses, as these filters were implemented to reduce the probability of false positives, and were less relevant in stage 2. Stage 2 SNPs were evaluated in all ancestry groups and for all traits, no matter what specific meta-analysis met the p-value threshold in the stage 1 analysis. We did not apply genomic control to any of the Stage 2 analyses given the expectation of association.

An additional meta-analysis was performed combining the Stage 1 and 2 meta-analyses. SNPs (irrespective of being known or novel) were considered to be replicated when Stage 1 p-values <5×10^-7^, Stage 2 p-value <0.05 with a similar direction of effect as in the discovery meta-analysis, and Stage 1+2 p-value <5×10^-8^. Replicated SNPs were subsequently used in different bioinformatics tools for further processing. In addition, 1 df p-values (SNP-sleep interaction effect only) of the lead SNPs of both the replicated known and novel genetic loci were calculated to explore whether genetic variant were specifically driven by SNP-main or SNP-interaction effects. Based on the total number of lead SNPs across all analyses, we performed correction using the false-discovery rate to quantify statistical significance ^36^.

### Bioinformatics

Replicated SNPs were first processed using the online tool FUMA^97^ to identify independent lead SNPs and to perform gene mapping. From the SNP that has a P_joint_ < 5×10^-8^, we determined lead SNPs that were independent from each other at R^2^ < 0.1 using the 1000G Phase 3 EUR as a reference panel population. Independent lead SNPs with a physical distance >1 mB from a known locus were considered as novel. Regional plots of the novel loci from the European- and multi-ancestry meta-analyses were made using the online LocusZoom tool ^98^. The explained variance of the newly identified genetic independent variants was calculated based on the summary statistics of the combined analysis of Stage 1 and 2 using the R-based VarExp package, which has been previously validated to provide similar results to individual participant data ^35^. Differential expression analyses of the lead SNPs in the newly identified genetic loci was performed using GTEx (https://gtexportal.org/home/) ^37,38^.

### Look-ups of novel loci in publicly available databases

Newly identified genetic loci for the three lipid traits were further explored in the GWAS catalogue (https://www.ebi.ac.uk/gwas/) to investigate the role of the newly identified mapped genes in other traits. Furthermore, we extracted the lead SNPs from the novel genetic loci from publically available GWAS data from the UK Biobank (http://www.nealelab.is/uk-biobank/) for different questionnaire-based sleep phenotypes, notably “daytime snoozing/sleeping (narcolepsy)”, “getting up in the morning”, “morning/evening person (chronotype)”, “nap during the day”, “sleep duration”, “sleeplessness/insomnia”, and “snoring”. GWAS in the UK Biobank were done in European-ancestry individuals only (N up to 337,074). In addition, we extracted the newly identified lead SNPs from publically available summary-statistics data on coronary artery disease of the CARDIoGRAMplusC4D consortium, which included 60,801 cases of coronary artery disease and 123,504 controls ^99^.

## Acknowledgments

This project was supported by a grant from the US National Heart, Lung, and Blood Institute (NHLBI) of the National Institutes of Health (R01HL118305). This research was supported in part by the Intramural Research Program of the National Human Genome Research Institute, National Institutes of Health. Tuomas O. Kilpeläinen was supported by the Danish Council for Independent Research (DFF – 6110-00183) and the Novo Nordisk Foundation (NNF18CC0034900, NNF17OC0026848 and NNF15CC0018486). Diana van Heemst was supported by the European Commission funded project HUMAN (Health-2013-INNOVATION-1-602757). Susan Redline was supported in part by NIH R35HL135818 and HL11338. Study-specific acknowledgments can be found in the **Supplementary Materials**. Data on coronary artery disease have been contributed by the Myocardial Infarction Genetics and CARDIoGRAM investigators and have been downloaded from www.CARDIOGRAMPLUSC4D.ORG.

## Author contribution

RN and MMB conducted the centralized data analysis, which included quality control checks, meta-analyses and bioinformatics. RN, MMB and SR drafted the initial version of the manuscript. RN, MMB, HW, TWW, ARB, TOK, PBM, CTL, ACM, DCR, DvH and SR were part of the writing group and were mainly responsible for the study design, interpretation of the data, and critical commenting on the initial draft versions of the manuscript. All other co-authors were responsible for cohort-level data collection, cohort-level data analysis, and critical reviews of the draft manuscript. All authors approved the final version of the manuscript that was submitted to the journal.

