## Supplementary Methods and Figures for "Multi-ancestry analysis of gene-sleep interactions in 126,926 individuals identifies multiple novel blood lipid loci that contribute to our understanding of sleep-associated adverse blood lipid profile"

**Supplementary Methods and Figures, R.Noordam et al**

| **Stage 1 Study Descriptions** | 2 |
| --- | --- |
| **Stage 1 Study Acknowledgements** | 9 |
| **Stage 2 Study Descriptions** | 14 |
| **Stage 2 Study Acknowledgements** | 20 |
| **Supplementary Figure 1**: Regional plots of the replicated novel hits of the joint effect of long total sleep time on HDL cholesterol in the multi-ancestry meta-analyses | 24 |
| **Supplementary Figure 2**: Regional plots of the replicated novel hits of the joint effect of short total sleep time on HDL cholesterol in the multi-ancestry meta-analyses | 28 |
| **Supplementary Figure 3**: Regional plots of the replicated novel hits of the joint effect of long total sleep time on LDL cholesterol in the multi-ancestry meta-analyses | 32 |
| **Supplementary Figure 4**: Regional plots of the replicated novel hits of the joint effect of short total sleep time on LDL cholesterol in the multi-ancestry meta-analyses | 34 |
| **Supplementary Figure 5:** Regional plots of the replicated novel hits of the joint effect of long total sleep time on Triglycerides in the multi-ancestry meta-analyses | 38 |
| **Supplementary Figure 6:** Regional plots of the replicated novel hits of the joint effect of short total sleep time on Triglycerides cholesterol in the multi-ancestry meta-analyses | 42 |
| **Supplementary Figure 7:** Project overview and SNP selection in the European-ancestry analyses | 49 |
| **Supplementary Figure 8:** Circular –log(p-value) plots of the European-ancestry sleep-SNP interactions analyses for the three lipid traits | 50 |
| **Supplementary Figure 9:** Regional plots of the replicated additional novel hits of the joint effect of long total sleep time on HDL cholesterol in the European meta-analyses | 52 |
| **Supplementary Figure 10:** Regional plots of the replicated additional novel hits of the joint effect of long short sleep time on HDL cholesterol in the European meta-analyses | 54 |
| **Supplementary Figure 11:** Regional plots of the replicated additional novel hits of the joint effect of long total sleep time on LDL cholesterol in the European meta-analyses | 55 |
| **Supplementary Figure 12:** Regional plots of the replicated additional novel hits of the joint effect of long total sleep time on Triglycerides in the European meta-analyses | 56 |
| **Supplementary Figure 13:** Regional plots of the replicated additional novel hits of the joint effect of short total sleep time on Triglycerides in the European meta-analyses | 57 |

**STAGE 1 STUDY DESCRIPTIONS:**

Brief descriptions are provided below for each of the discovery studies, some of which are based outside the United States:

**AGES (Age Gene/Environment Susceptibility Reykjavik Study):** The AGES Reykjavik study originally comprised a random sample of 30,795 men and women born in 1907-1935 and living in Reykjavik, Iceland in 1967. A total of 19,381 people attended, resulting in a 71% recruitment rate. The study sample was divided into six groups by birth year and birth date within month. One group was designated for longitudinal follow up and was examined in all stages; another was designated as a control group and was not included in examinations until 1991. Other groups were invited to participate in specific stages of the study. Between 2002 and 2006, the AGES Reykjavik study re-examined 5,764 survivors of the original cohort who had participated before in the Reykjavik Study. Blood samples were drawn from all participants after overnight fasting. Serum total cholesterol and HDL were analyzed on a Hitachi 912 using reagents from Roche Diagnostics (Mannheim, Germany) and following the manufacturer’s instructions (CV% was 2.5% and 3.5% respectively). Serum LDL cholesterol was calculated using the Friedewald equation when triglycerides < 4.5 mmol/L. Information on sleep duration was collected using the single question for both summer and winter: “During a typical 24-hour period in the summer time/winter time, how many hours do you spend sleeping at night?” Genotyping was performed using the Illumina 370CNV BeadChip array on 3,664 participants. Sample exclusion criteria included sample failure, genotype mismatch with reference panel, and sex mismatch, resulting in clean genotype data on 3,219 individuals. Standard protocols for working with Illumina data were followed with clustering score greater than 0.4. From a total of 353,202 SNPS, 325,094 were used for imputation after exclusion of SNPs with call rate < 97%, HWE deviation < 1 x 10^-6^, mishap (PLINK haplotype-based test for non-random missing genotype data[2]) p < 1 x 10^-9^, and mismatched positions between Illumina, dbSNP and/or HapMap. Imputation was done using MACH1.0.16 against all the HapMap CEPH haplotypes (release 22/NCBI build 36) resulting in 2,533,153 total SNPs for analysis. All participants signed an informed consent form. The AGES-Reykjavik Study GWAS was approved by the National Bioethics Committee (VSN: 00-063) and the Data Protection Authority. A detailed description of the AGES Reykjavik Study has been described previously ([Harris TB *et al.*](http://www.ncbi.nlm.nih.gov/pubmed/17351290?ordinalpos=5&itool=EntrezSystem2.PEntrez.Pubmed.Pubmed_ResultsPanel.Pubmed_RVDocSum) Age, Gene/Environment Susceptibility-Reykjavik Study: multidisciplinary applied phenomics.Am J Epidemiol. 2007 May 1;165(9):1076-87. Epub 2007 Mar 10. PMID:17351290)

**ARIC (Atherosclerosis Risk in Communities):** The ARIC study is a population-based prospective cohort study of cardiovascular disease sponsored by the National Heart, Lung, and Blood Institute (NHLBI). ARIC included 15,792 individuals, predominantly European American and African American, aged 45-64 years at baseline (1987-89), chosen by probability sampling from four US communities. Cohort members completed three additional triennial follow-up examinations, a fifth exam in 2011-2013, and a sixth exam in 2016-2017. The ARIC study has been described in detail previously (The ARIC Investigators. The Atherosclerosis Risk in Communities (ARIC) study: Design and objectives. Am J Epidemiol. 1989;129:687-702).

**Baependi Heart Study (Brazil):** The Baependi Heart Study, is an ongoing family-based cohort conducted in a rural town of the state of Minas Gerais. The study has enrolled approximate 2,200 individuals (over 10% of the town’s adult population) and 10-year follow up period of longitudinal data. Briefly, probands were selected at random across 11 out of the 12 census districts in Baependi. After enrolment, the proband's first-degree (parents, siblings, and offspring), second-degree (half-siblings, grandparents/grandchildren, uncles/aunts, nephews/nieces, and double cousins), and third-degree (first cousins, great uncles/aunts, and great nephews/nieces) relatives, and his/her respective spouse's relatives resident both within Baependi (municipal and rural area) and surrounding towns were invited to participate. Only individuals age 18 and older were eligible to participate in the study. The study is conducted from a clinic/office in an easily accessible sector of the town, where the questionnaires were completed. A broad range of phenotypes ranging from cardiovascular, neurocognitive, psychiatric, imaging, physiologic and several layers of endophenotypes like metabolomics and lipidomics have been collected throughout the years Details about follow-up visits and available data can be found in the cohort profile paper (PMID: 18430212). DNA samples were genotyped using the Affymetrix 6.0 genechip. After quality control, the data were prephased using SHAPEIT and imputed using IMPUTE2 based on 1000 Genomes haplotypes.

**CARDIA (Coronary Artery Risk Development in Young Adults):** CARDIA is a prospective multicenter study with 5,115 adults Caucasian and African American participants of the age group 18-30 years, recruited from four centers at the baseline examination in 1985-1986. The recruitment was done from the total community in Birmingham, AL, from selected census tracts in Chicago, IL and Minneapolis, MN; and from the Kaiser Permanente health plan membership in Oakland, CA. The details of the study design for the CARDIA study have been previously published^1^. Nine examinations have been completed since initiation of the study, respectively in the years 0, 2, 5, 7, 10, 15, 20, 25 and 30. Written informed consent was obtained from participants at each examination and all study protocols were approved by the institutional review boards of the participating institutions. Age and race were self‐reported using standardized questionnaires, as were use of cholesterol‐lowering medication. participants were asked to fast for 12 hours before each clinic visit.

1.	Friedman GD, Cutter GR, Donahue RP, Hughes GH, Hully SB, Jacobs DR Jr., Liu K, Savage PJ. CARDIA: study design, recruitment, and some characteristics of the examined subjects. J Clin Epidemiol. 1988;41:1105–1116.

**CHS (Cardiovascular Health Study):** CHS is a population-based cohort study of risk factors for cardiovascular disease in adults 65 years of age or older conducted across four field centers (1). The original predominantly European ancestry cohort of 5,201 persons was recruited in 1989-1990 from random samples of the Medicare eligibility lists and an additional predominately African-American cohort of 687 persons was enrolled in 1992-93 for a total sample of 5,888. Blood samples were drawn from all participants at their baseline examination and DNA was subsequently extracted from available samples. European ancestry participants were excluded from the GWAS study sample due to prevalent coronary heart disease, congestive heart failure, peripheral vascular disease, valvular heart disease, stroke, or transient ischemic attack at baseline. After QC, genotyping was successful for 3271 European ancestry and 823 African-American participants. CHS was approved by institutional review committees at each site and individuals in the present analysis gave informed consent including consent to use of genetic information for the study of cardiovascular disease. Information on habitual sleep duration was collected 4 years after the cholesterol measurement.

1.	Fried LP, Borhani NO, Enright P, Furberg CD, Gardin JM, Kronmal RA, et al. The Cardiovascular Health Study: design and rationale. Ann Epidemiol 1991; 1:263-76.

**ERF (Erasmus Rucphen Family study):** Erasmus Rucphen Family is a family based study that includes inhabitants of a genetically isolated community in the South-West of the Netherlands, studied as part of the Genetic Research in Isolated Population (GRIP) program (Padro LM, et al; Ann Hum Genet. 2015l 69(Pt 3):288-95). The goal of the study is to identify the risk factors in the development of complex disorders. Study population includes approximately 3,000 individuals who are living descendants of 22 couples who lived in the isolate between 1850 and 1900 and had at least six children baptized in the community church (Aulchenko Y, et al; Eur J Hum Genet. 2015; 12 527-534). All data were collected between 2002 and 2005. All participants gave informed consent, and the Medical Ethics Committee of the Erasmus University Medical Centre approved the study.

**FHS (Framingham Heart Study):** FHS began in 1948 with the recruitment of an original cohort of 5,209 men and women (mean age 44 years; 55 percent women). In 1971 a second generation of study participants was enrolled; this cohort (mean age 37 years; 52% women) consisted of 5,124 children and spouses of children of the original cohort. A third generation cohort of 4,095 children of offspring cohort participants (mean age 40 years; 53 percent women) was enrolled in 2002-2005 and are seen every 4 to 8 years. Details of study designs for the three cohorts are summarized elsewhere. At each clinic visit, a medical history was obtained with a focus on cardiovascular content, and participants underwent a physical examination including measurement of height and weight from which BMI was calculated.

**GenSalt (Genetic Epidemiology Network of Salt Sensitivity):** GenSalt is a multi-center, family based study designed to identify, through dietary sodium and potassium intervention, salt-sensitivity susceptibility genes which may underlie essential hypertension in rural Han Chinese families. Approximately 629 families with at least one ‘proband’ with high blood pressure were recruited and tested for a wide variety of physiological, metabolic and biochemical measures at baseline and at multiple times during the 3-week intervention. The intervention consisted of one week on a low sodium diet, followed by one week on a high sodium diet, and finally one week on a high sodium diet with a potassium supplement.

**HANDLS (Healthy Aging in Neighborhoods of Diversity across the Life Span):** HANDLS is a community-based, longitudinal epidemiologic study examining the influences of race and socioeconomic status (SES) on the development of age-related health disparities among a sample of socioeconomically diverse African Americans and whites. This unique study will assess over a 20-year period physical parameters and also evaluate genetic, biologic, demographic, and psychosocial, parameters of African American and white participants in higher and lower SES to understand the driving factors behind persistent black-white health disparities in overall longevity, cardiovascular disease, and cognitive decline. The study recruited 3,722 participants from Baltimore, MD with a mean age of 47.7 years, 2,200 African Americans and 1,522 whites, with 41% reporting household incomes below the 125% poverty delimiter. Genotyping was done on a subset of self-reporting African American participants by the Laboratory of Neurogenetics, National Institute on Aging, National Institutes of Health (NIH). A larger genotyping effort included a small subset of self-reporting European ancestry samples. This research was supported by the Intramural Research Program of the NIH, NIA and the National Center on Minority Health and Health Disparities.

**Health ABC (Health, Aging, and Body Composition):** Cohort description: The Health ABC study is a prospective cohort study investigating the associations between body composition, weight-related health conditions, and incident functional limitation in older adults. Health ABC enrolled well-functioning, community-dwelling black (n=1281) and white (n=1794) men and women aged 70-79 years between April 1997 and June 1998. Participants were recruited from a random sample of white and all black Medicare eligible residents in the Pittsburgh, PA, and Memphis, TN, metropolitan areas. Participants have undergone annual exams and semi-annual phone interviews. The current study sample consists of 1559 white participants who attended the second exam in 1998-1999 with available genotyping data.

Genotyping: Genotyping was performed by the Center for Inherited Disease Research (CIDR) using the Illumina Human1M-Duo BeadChip system. Samples were excluded from the dataset for the reasons of sample failure, genotypic sex mismatch, and first-degree relative of an included individual based on genotype data. Genotyping was successful in 1663 Caucasians. Analysis was restricted to SNPs with minor allele frequency ≥ 1%, call rate ≥97% and HWE p≥10-6. Genotypes were available on 914,263 high quality SNPs for imputation based on the HapMap CEU (release 22, build 36) using the MACH software (version 1.0.16). A total of 2,543,888 imputed SNPs were analyzed for association with vitamin D levels.

**HERITAGE (Health, Risk Factors, Exercise Training and Genetics):** The HERITAGE is the only known family-based study of exercise intervention to evaluate the role of genes and sequence variants involved in the response to a physically active lifestyle. The current study is based on the data collected at baseline of the study from 99 White families (244 males, 255 females). All subjects were required to be sedentary and free of chronic diseases at baseline. There are over 18 trait domains (e.g. dietary, lipids and lipoproteins, glucose and insulin metabolism [fasting and IVGTT], steroids, body composition and body fat distribution, cardiorespiratory fitness), for a grand total of over one thousand variables. Moreover, most of the outcome traits were measured twice on two separate days both at baseline and after exercise training was completed. Marker data include a genome-wide linkage scan and GWAS, in addition to a large number of candidate genes.

**JHS (Jackson Heart Study):** The Jackson Heart Study is a longitudinal, community-based observational cohort study investigating the role of environmental and genetic factors in the development of cardiovascular disease in African Americans. Between 2000 and 2004, a total of 5,306 participants were recruited from a tri-county area (Hinds, Madison, and Rankin Counties) that encompasses Jackson, MS. Details of the design and recruitment for the Jackson Heart Study cohort has been previously published.1-3 Briefly, approximately 30% of participants were former members of the Atherosclerosis Risk in Communities (ARIC) study. The remainder were recruited by either 1) random selection from the Accudata list, 2) commercial listing, 3) a constrained volunteer sample, in which recruitment was distributed among defined demographic cells in proportions designed to mirror those in the overall population, or through the Jackson Heart Study Family Study.

1.	Wyatt SB, Diekelmann N, Henderson F, Andrew ME, Billingsley G, Felder SH et al. A community-driven model of research participation: the Jackson Heart Study Participant Recruitment and Retention Study. Ethn Dis 2003; 13(4):438-455.

2.	Taylor HA, Jr., Wilson JG, Jones DW, et al. Toward resolution of cardiovascular health disparities in African Americans: design and methods of the Jackson Heart Study. Ethn Dis 2005; 15:S6-17.

3.	Fuqua SR, Wyatt SB, Andrew ME, et al. Recruiting African-American research participation in the Jackson Heart Study: methods, response rates, and sample description. Ethn Dis 2005; 15:S6-29.

**MESA (Multi-Ethnic Study of Atherosclerosis):** The Multi-Ethnic Study of Atherosclerosis (MESA) is a study of the characteristics of subclinical cardiovascular disease and the risk factors that predict progression to clinically overt cardiovascular disease or progression of the subclinical disease. MESA consisted of a diverse, population-based sample of an initial 6,814 asymptomatic men and women aged 45-84. 38 percent of the recruited participants were white, 28 percent African American, 22 percent Hispanic, and 12 percent Asian, predominantly of Chinese descent. Participants were recruited from six field centers across the United States: Wake Forest University, Columbia University, Johns Hopkins University, University of Minnesota, Northwestern University and University of California - Los Angeles. Participants are being followed for identification and characterization of cardiovascular disease events, including acute myocardial infarction and other forms of coronary heart disease (CHD), stroke, and congestive heart failure; for cardiovascular disease interventions; and for mortality. The first examination took place over two years, from July 2000 - July 2002. It was followed by four examination periods that were 17-20 months in length. Participants have been contacted every 9 to 12 months throughout the study to assess clinical morbidity and mortality.

Bild DE, Bluemke DA, Burke GL, Detrano R, Diez Roux AV, Folsom AR, Greenland P, Jacob DR Jr, Kronmal R, Liu K, Nelson JC, O'Leary D, Saad MF, Shea S, Szklo M, Tracy RP. Multi-ethnic study of atherosclerosis: objectives and design. Am J Epidemiol. 2002 Nov 1;156(9):871-81. PubMed PMID: 12397006.

**NEO (The Netherlands Epidemiology of Obesity study):** The NEO was designed for extensive phenotyping to investigate pathways that lead to obesity-related diseases. The NEO study is a population-based, prospective cohort study that includes 6,671 individuals aged 45–65 years, with an oversampling of individuals with overweight or obesity. At baseline, information on demography, lifestyle, and medical history have been collected by questionnaires. In addition, samples of 24-h urine, fasting and postprandial blood plasma and serum, and DNA were collected. Genotyping was performed using the Illumina HumanCoreExome chip, which was subsequently imputed to the 1000 genome reference panel. Participants underwent an extensive physical examination, including anthropometry, electrocardiography, spirometry, and measurement of the carotid artery intima-media thickness by ultrasonography. The collection of data started in September 2008 and completed at the end of September 2012.

**RS (Rotterdam Study):** The Rotterdam Study is a prospective, population-based cohort study among individuals living in the well-defined Ommoord district in the city of Rotterdam in The Netherlands (Ikram MA, et al; Eur J Epidemiol. 2017; 32(9): 807-850). The aim of the study is to determine the occurrence of cardiovascular, neurological, ophthalmic, endocrine, hepatic, respiratory, and psychiatric diseases in elderly people. The cohort was initially defined in 1990 with 7,983 persons, aged 55 years and older, who underwent a home interview and extensive physical examination at the baseline and during follow-up rounds every 3-4 years (RS-I). Cohort was extended in 2000/2001 (RS-II, 3,011 individuals aged 55 years and older) and 2006/2008 (RS-III, 3,932 subjects, aged 45 and older). The Rotterdam Study has been approved by the Medical Ethics Committee of the Erasmus MC (registration number MEC 02.1015) and by the Dutch Ministry of Health, Welfare and Sport (Population Screening Act WBO, license number 1071272-159521-PG). All participants provided written informed consent to participate in the study and to have their medical information obtained from treating physicians.

**WHI (Women’s Health Initiative):** WHI is a long-term national health study that focuses on strategies for preventing common diseases such as heart disease, cancer and fracture in postmenopausal women. A total of 161,838 women aged 50–79 years old were recruited from 40 clinical centers in the US between 1993 and 1998. WHI consists of an observational study, two clinical trials of postmenopausal hormone therapy (HT, estrogen alone or estrogen plus progestin), a calcium and vitamin D supplement trial, and a dietary modification trial ^1^. Study recruitment and exclusion criteria have been described previously ^1^. Recruitment was done through mass mailing to age-eligible women obtained from voter registration, driver’s license and Health Care Financing Administration or other insurance list, with emphasis on recruitment of minorities and older women ^2^. Exclusions included participation in other randomized trials, predicted survival < 3 years, alcoholism, drug dependency, mental illness and dementia. Study protocols and consent forms were approved by the IRB at all participating institutions. Socio-demographic characteristics, lifestyle, medical history and self-reported medications were collected using standardized questionnaires at the screening visit. The genome wide association study (GWAS) non-overlapping samples are composed of a case-control study (WHI Genomics and Randomized Trials Network – GARNET, which included all coronary heart disease, stroke, venous thromboembolic events and selected diabetes cases that happened during the active intervention phase in the WHI HT clinical trials and aged matched controls), women selected to be "representative" of the HT trial (mostly younger white HT subjects that were also enrolled in the WHI memory study - WHIMS) and the WHI SNP Health Association Resource (WHI SHARe), a randomly selected sample of 8,515 African American and 3,642 Hispanic women from WHI. GWAS was performed using Affymetrix 6.0 (WHI-SHARe), HumanOmniExpressExome-8v1_B (WHIMS), Illumina HumanOmni1-Quad v1-0 B (GARNET). Extensive quality control (QC) of the GWAS data included alignment (“flipping”) to the same reference panel, imputation to the 1000G data (using the recent reference panel - v3.20101123), identification of genetically related individuals, and computations of principal components (PCs) using methods developed by Price et al. (using EIGENSOFT software 53), and finally the comparison with self-reported ethnicity. After QC and exclusions from analysis protocol, the number of women included in analysis is 4,423 whites for GARNET, 5,202 white for WHIMS, 7,919 for SHARe African American and 3,377 for SHARe Hispanics.

1.	Hays J, Hunt JR, Hubbell FA, Anderson GL, Limacher M, Allen C, Rossouw JE. The women's health initiative recruitment methods and results. Ann Epidemiol. 2003;13:S18-77

2.	Design of the women's health initiative clinical trial and observational study. The women's health initiative study group. Control Clin Trials. 1998;19:61-109

**STAGE 1 STUDY ACKNOWLEDGMENTS:**

Infrastructure for the CHARGE Consortium is supported in part by the National Heart, Lung, and Blood Institute grant R01HL105756. Infrastructure for the Gene-Lifestyle Working Group is supported by the National Heart, Lung, and Blood Institute grant R01HL118305.

**AGES (Age Gene/Environment Susceptibility Reykjavik Study):** This study has been funded by NIH contract N01-AG012100, HSSN271201200022C, the NIA Intramural Research Program, an Intramural Research Program Award (ZIAEY000401) from the National Eye Institute, an award from the National Institute on Deafness and Other Communication Disorders (NIDCD) Division of Scientific Programs (IAA Y2-DC_1004-02), Hjartavernd (the Icelandic Heart Association), and the Althingi (the Icelandic Parliament). The study is approved by the Icelandic National Bioethics Committee, VSN: 00-063. The researchers are indebted to the participants for their willingness to participate in the study.

**ARIC (Atherosclerosis Risk in Communities) study:** The ARIC study has been funded in whole or in part with Federal funds from the National Heart, Lung, and Blood Institute, National Institutes of Health, Department of Health and Human Services (contract numbers HHSN268201700001I, HHSN268201700002I, HHSN268201700003I, HHSN268201700004I and HHSN268201700005I), R01HL087641, R01HL059367 and R01HL086694; National Human Genome Research Institute contract U01HG004402; and National Institutes of Health contract HHSN268200625226C. The authors thank the staff and participants of the ARIC study for their important contributions. Infrastructure was partly supported by Grant Number UL1RR025005, a component of the National Institutes of Health and NIH Roadmap for Medical Research.

**Baependi Heart Study (Brazil):** The Baependi Heart Study was supported by Fundação de Amparo a Pesquisa do Estado de São Paulo (FAPESP) (Grant 2013/17368-0), Coordenação de Aperfeiçoamento de Pessoal de Nível Superior (CAPES) and Hospital Samaritano Society (Grant 25000.180.664/2011-35), through Ministry of Health to Support Program Institutional Development of the Unified Health System (SUS-PROADI).

**CARDIA (Coronary Artery Risk Development in Young Adults):** The CARDIA Study is conducted and supported by the National Heart, Lung, and Blood Institute in collaboration with the University of Alabama at Birmingham (HHSN268201300025C & HHSN268201300026C), Northwestern University (HHSN268201300027C), University of Minnesota (HHSN268201300028C), Kaiser Foundation Research Institute (HHSN268201300029C), and Johns Hopkins University School of Medicine (HHSN268200900041C). CARDIA is also partially supported by the Intramural Research Program of the National Institute on Aging. Genotyping was funded as part of the NHLBI Candidate-gene Association Resource (N01-HC-65226) and the NHGRI Gene Environment Association Studies (GENEVA) (U01-HG004729, U01-HG04424, and U01-HG004446). This manuscript has been reviewed and approved by CARDIA for scientific content.

**CHS (Cardiovascular Health Study):** This Cardiovascular Health Study (CHS) research was supported by NHLBI contracts HHSN268201200036C, HHSN268200800007C, HHSN268200960009C, HHSN268201800001C N01HC55222, N01HC85079, N01HC85080, N01HC85081, N01HC85082, N01HC85083, N01HC85086; and NHLBI grants U01HL080295, U01HL130114, R01HL087652, R01HL105756, R01HL103612, R01HL085251, and R01HL120393 with additional contribution from the National Institute of Neurological Disorders and Stroke (NINDS). Additional support was provided through R01AG023629 from the National Institute on Aging (NIA). A full list of principal CHS investigators and institutions can be found at CHS-NHLBI.org. The provision of genotyping data was supported in part by the National Center for Advancing Translational Sciences, CTSI grant UL1TR001881, and the National Institute of Diabetes and Digestive and Kidney Disease Diabetes Research Center (DRC) grant DK063491 to the Southern California Diabetes Endocrinology Research Center. The content is solely the responsibility of the authors and does not necessarily represent the official views of the National Institutes of Health.

**ERF (Erasmus Rucphen Family study):** The ERF study as a part of EUROSPAN (European Special Populations Research Network) was supported by European Commission FP6 STRP grant number 018947 (LSHG-CT-2006-01947) and also received funding from the European Community's Seventh Framework Programme (FP7/2007-2013)/grant agreement HEALTH-F4-2007-201413 by the European Commission under the programme "Quality of Life and Management of the Living Resources" of 5th Framework Programme (no. QLG2-CT-2002-01254). The ERF study was further supported by ENGAGE consortium and CMSB. High-throughput analysis of the ERF data was supported by joint grant from Netherlands Organisation for Scientific Research and the Russian Foundation for Basic Research (NWO-RFBR 047.017.043). This study was further supported by the European Union's Horizon 2020 research and innovation programme as part of the Common mechanisms and pathways in Stroke and Alzheimer's disease (CoSTREAM) project (www.costream.eu, grant agreement No 667375); the Netherlands Organisation for Health Research and Development (ZonMW) as part of the Joint Programming for Neurological Disease (JPND) project PERADES (Defining Genetic, Polygenic and Environmental Risk for Alzheimer’s Disease using multiple powerful cohorts, focused Epigenetics and Stem cell metabolomics - grant number 733051021); the European Union’s Horizon 2020 research and innovation programme Marie Skłodowska-Curie Research and Innovation Staff Exchange (RISE) under the grant agreement No 645740 as part of the Personalized pREvention of Chronic DIseases (PRECeDI) project and the CardioVasculair Onderzoek Nederland (CVON 2012-03). We are grateful to all study participants and their relatives, general practitioners and neurologists for their contributions and to P. Veraart for her help in genealogy, J. Vergeer for the supervision of the laboratory work, P. Snijders for his help in data collection and E.M. van Leeuwen for genetic imputation.

**FHS (Framingham Heart Study):** This research was conducted in part using data and resources from the Framingham Heart Study of the National Heart Lung and Blood Institute of the National Institutes of Health and Boston University School of Medicine. The analyses reflect intellectual input and resource development from the Framingham Heart Study investigators participating in the SNP Health Association Resource (SHARe) project. This work was partially supported by the National Heart, Lung and Blood Institute's Framingham Heart Study (Contract Nos. N01-HC-25195 and HHSN268201500001I) and its contract with Affymetrix, Inc for genotyping services (Contract No. N02-HL-6-4278). A portion of this research utilized the Linux Cluster for Genetic Analysis (LinGA-II) funded by the Robert Dawson Evans Endowment of the Department of Medicine at Boston University School of Medicine and Boston Medical Center. This research was partially supported by grant R01-DK089256 from the National Institute of Diabetes and Digestive and Kidney Diseases (MPIs: Michael Province, L. Ching-Ti Liu, Kari North) and training grant T32GM074905-14 from NIH/National Institute of General Medical Sciences.

**GenSalt (Genetic Epidemiology Network of Salt Sensitivity):** The Genetic Epidemiology Network of Salt Sensitivity is supported by research grants (U01HL072507, R01HL087263, and R01HL090682) from the National Heart, Lung, and Blood Institute, National Institutes of Health, Bethesda, MD.

**HANDLS (Healthy Aging in Neighborhoods of Diversity across the Life Span):** The Healthy Aging in Neighborhoods of Diversity across the Life Span (HANDLS) study was supported by the Intramural Research Program of the NIH, National Institute on Aging and the National Center on Minority Health and Health Disparities (project # Z01-AG000513 and human subjects protocol number 09-AG-N248). Data analyses for the HANDLS study utilized the high-performance computational resources of the Biowulf Linux cluster at the National Institutes of Health, Bethesda, MD. (http://biowulf.nih.gov; http://hpc.nih.gov)).

**Health ABC (Health, Aging, and Body Composition):** Health ABC was funded by the National Institutes of Aging. This research was supported by NIA contracts N01AG62101, N01AG62103, and N01AG62106. The GWAS was funded by NIA grant 1R01AG032098-01A1 to Wake Forest University Health Sciences and genotyping services were provided by the Center for Inherited Disease Research (CIDR). CIDR is fully funded through a federal contract from the National Institutes of Health to The Johns Hopkins University, contract number HHSN268200782096C. This research was supported in part by the Intramural Research Program of the NIH, National Institute on Aging.

**HERITAGE (Health, Risk Factors, Exercise Training and Genetics):** The HERITAGE Family Study was supported by National Heart, Lung, and Blood Institute grant HL-45670.

**JHS (Jackson Heart Study):** The Jackson Heart Study is supported by contracts HSN268201300046C, HHSN268201300047C, HHSN268201300048C, HHSN268201300049C, HHSN268201300050C from the National Heart, Lung, and Blood Institute on Minority Health and Health Disparities. The authors acknowledge the Jackson Heart Study team institutions (University of Mississippi Medical Center, Jackson State University and Tougaloo College) and participants for their long-term commitment that continues to improve our understanding of the genetic epidemiology of cardiovascular and other chronic diseases among African Americans.

**MESA (Multi-Ethnic Study of Atherosclerosis):** This research was supported by the Multi-Ethnic Study of Atherosclerosis (MESA) contracts HHSN268201500003I, N01-HC-95159, N01-HC-95160, N01-HC-95161, N01-HC-95162, N01-HC-95163, N01-HC-95164, N01-HC-95165, N01-HC-95166, N01-HC-95167, N01-HC-95168, N01-HC-95169, UL1-TR-000040, UL1-TR-001079, and UL1-TR-001420.  Funding for MESA Share genotyping was provided by NHLBI Contract N02-HL-6-4278. This publication was partially developed under a STAR research assistance agreement, No. RD831697 (MESA Air), awarded by the U.S Environmental Protection Agency. It has not been formally reviewed by the EPA. The views expressed in this document are solely those of the authors and the EPA does not endorse any products or commercial services mentioned in this publication. The provision of genotyping data was supported in part by the National Center for Advancing Translational Sciences, CTSI grant UL1TR001881, and the National Institute of Diabetes and Digestive and Kidney Disease Diabetes Research Center (DRC) grant DK063491 to the Southern California Diabetes Endocrinology Research Center.  The authors thank the participants of the MESA study, the Coordinating Center, MESA investigators, and study staff for their valuable contributions.  A full list of participating MESA investigators and institutions can be found at <http://www.mesa-nhlbi.org>.

**NEO (The Netherlands Epidemiology of Obesity study):** The authors of the NEO study thank all individuals who participated in the Netherlands Epidemiology in Obesity study, all participating general practitioners for inviting eligible participants and all research nurses for collection of the data. We thank the NEO study group, Petra Noordijk, Pat van Beelen and Ingeborg de Jonge for the coordination, lab and data management of the NEO study. The genotyping in the NEO study was supported by the Centre National de Génotypage (Paris, France), headed by Jean-Francois Deleuze. The NEO study is supported by the participating Departments, the Division and the Board of Directors of the Leiden University Medical Center, and by the Leiden University, Research Profile Area Vascular and Regenerative Medicine. Dennis Mook-Kanamori is supported by Dutch Science Organization (ZonMW-VENI Grant 916.14.023).

**RS (Rotterdam Study):** The Rotterdam Study is funded by Erasmus Medical Center and Erasmus University, Rotterdam, Netherlands Organization for the Health Research and Development (ZonMw), the Research Institute for Diseases in the Elderly (RIDE), the Ministry of Education, Culture and Science, the Ministry for Health, Welfare and Sports, the European Commission (DG XII), and the Municipality of Rotterdam. The authors are grateful to the study participants, the staff from the Rotterdam Study and the participating general practitioners and pharmacists.

The generation and management of GWAS genotype data for the Rotterdam Study was executed by the Human Genotyping Facility of the Genetic Laboratory of the Department of Internal Medicine, Erasmus MC, Rotterdam, The Netherlands. The GWAS datasets are supported by the Netherlands Organisation of Scientific Research NWO Investments (nr. 175.010.2005.011, 911-03-012), the Genetic Laboratory of the Department of Internal Medicine, Erasmus MC, the Research Institute for Diseases in the Elderly (014-93-015; RIDE2), the Netherlands Genomics Initiative (NGI)/Netherlands Organisation for Scientific Research (NWO) Netherlands Consortium for Healthy Aging (NCHA), project nr. 050-060-810. We thank Pascal Arp, Mila Jhamai, Marijn Verkerk, Lizbeth Herrera, Marjolein Peters and Carolina Medina-Gomez for their help in creating the GWAS database, and Karol Estrada, Yurii Aulchenko and Carolina Medina-Gomez for the creation and analysis of imputed data. This study was further supported by the European Union's Horizon 2020 research and innovation programme as part of the Common mechanisms and pathways in Stroke and Alzheimer's disease (CoSTREAM) project (www.costream.eu, grant agreement No 667375); the Netherlands Organisation for Health Research and Development (ZonMW) as part of the Joint Programming for Neurological Disease (JPND) project PERADES (Defining Genetic, Polygenic and Environmental Risk for Alzheimer’s Disease using multiple powerful cohorts, focused Epigenetics and Stem cell metabolomics - grant number 733051021); the European Union’s Horizon 2020 research and innovation programme Marie Skłodowska-Curie Research and Innovation Staff Exchange (RISE) under the grant agreement No 645740 as part of the Personalized pREvention of Chronic DIseases (PRECeDI) project and the CardioVasculair Onderzoek Nederland (CVON 2012-03).

**WHI (Women’s Health Initiative):** The WHI program is funded by the National Heart, Lung, and Blood Institute, National Institutes of Health, U.S. Department of Health and Human Services through contracts HHSN268201100046C, HHSN268201100001C, HHSN268201100002C, HHSN268201100003C, HHSN268201100004C, and HHSN271201100004C. The authors thank the WHI investigators and staff for their dedication, and the study participants for making the program possible. A full listing of WHI investigators can be found at:

http://www.whi.org/researchers/Documents%20%20Write%20a%20Paper/WHI%20Investigator%20Short%20List.pdf

**STAGE 2 STUDY DESCRIPTIONS:**

Brief descriptions are provided below for each of the replication studies/cohorts:

**Airwave (The Airwave Health Monitoring Study):** The Airwave Health Monitoring Study (22) was established to evaluate possible health risks associated with use of TETRA, a digital communication system used by police forces and other emergency services in Great Britain since 2001. The study has been broadened to investigate more generally the health of the work force. From 2004, participants from each force who agreed to participate were enrolled either with an enrolment questionnaire or a comprehensive health screening performed locally. This includes questionnaire, 7-day food diaries, anthropometry, measurements of cardiovascular and cognitive function, blood chemistry, coagulation and hematology. By March 2015, the study had recruited 53,606 participants, of whom 45,433 had attended the health screening, and 14,002 have genotype data (1000G imputed).

Ref: Elliott, P. et al. The Airwave Health Monitoring Study of police officers and staff in Great Britain: rationale, design and methods. Environ Res 134, 280-5 (2014).

**CFS (Cleveland Family Study):** The Cleveland Family Study (CFS) is a family-based, longitudinal study designed to characterize the genetic and non-genetic risk factors for sleep apnea. In total, 2534 individuals (46% African American) from 352 families were studied on up to 4 occasions over a period of 16 years (1990-2006). The initial aim of the study was to quantify the familial aggregation of sleep apnea. 632 African Americans were genotyped on the Affymetrix array 6.0 platform through the CARe Consortium with suitable genotying quality control. A further 122 African-Americans had genotyping based on the Illumina OmniExpress + Exome platform. Genomes were imputed separately for each chip based on a 1000 Genomes Project Phase 3 Version 5 cosmopolitan template using SHAPEIT and IMPUTE2.

**Colaus (Cohorte Lausannoise):** The cohort is a random sample of the population aged 35-75 years of the city of Lausanne, Switzerland. Recruitment began in June 2003 and ended in May 2006, and the first follow-up was conducted between April 2009 and September 2012. The CoLaus study was approved by the Institutional Ethics Committee of the University of Lausanne and informed consent was appropriately obtained by all participants. Both at baseline and follow-up, all participants attended the outpatient clinic of the University Hospital of Lausanne in the morning after an overnight fast. Data were collected by trained field interviewers in a single visit lasting about 60 min.

**DFTJ (Dongfeng-Tongji Cohort Study):** The DFTJ-cohort study includes 27,009 retired employees from a state-owned automobile enterprise in China. This study was launched in 2008 and will be followed up every 5 years. In 2013 we conducted the first follow-up. By using semi-structural questionnaire and health examination, those having cancer or severe diseases were excluded. Fasting blood samples and detailed epidemiology data were collected. The main goal of the cohort was to identify the environmental and genetic risk factors and the gene-environment interactions on chronic diseases, and to find novel biomarkers for chronic disease and mortality prediction. Finally, 1,461 included in the present study with GWAS data. All of the participants wrote informed consent and the ethical committees in the Tongji Medical College approved this research project. Detailed information has been described in elsewhere(1).

QC criteria and imputation methods:

We did the GWAS scan on the DFTJ-cohort with Affymetrix Genome-Wide Human SNP Array 6.0 chips. In total, we genotyped 906,703 SNPs among 1,461 subjects. After stringent QC filtering, SNPs with MAF < 0.01, Hardy-Weinberg Equilibrium (HWE) < 0.0001, and SNP call rate < 95% were excluded. Individuals with call rates < 95% were also not included for further analysis. In total, we retained 1,452 subjects with 658,288 autosomal SNPs for statistical analyses, with an overall call rate of 99.68%. We used MACH 1.0 software to impute untyped SNPs using the LD information from the HapMap phase II database (CHB+JPT as a reference set (2007-08_rel22, released 2007-03-02). Imputed SNPs with high genotype information content (Rsq > 0.3 for MACH) were kept for the further association analysis.

**DR’s EXTRA (Dose-Responses to Exercise Training):** The Dose-Responses to Exercise Training (DR’s EXTRA) Study is a 4-year RCT on the effects of regular physical exercise and healthy diet on endothelial function, atherosclerosis and cognition in a randomly selected population sample (n=3000) of Eastern Finnish men and women, identified from the national population register, aged 55-74 years. Of the eligible sample, 1410 individuals were randomized into one of the 6 groups: aerobic exercise, resistance exercise, diet, combined aerobic exercise and diet, combined resistance exercise and diet, or reference group following baseline assessments. During the four year intervention the drop-out rate was 15%.

**EGCUT (Estonian Genome Center - University of Tartu (Estonian Biobank)):** The Estonian Biobank is a population-based biobank of the Estonian Genome Center at the University of Tartu ([www.biobank.ee](http://www.biobank.ee); EGCUT). The entire project is conducted according to the Estonian Gene Research Act and all of the participants have signed a broad informed consent. The cohort  comprises 51535 individuals, aged 18+ and closely reflects the age, sex and geographical distribution of the Estonian population. All of the subjects are recruited randomly by general practitioners and physicians at hospitals. A Computer Assisted Personal interview was filled within 1-2 hours at a doctor’s office, which included personal, genealogical, educational, occupational history and lifestyle data. Anthropometric measurements, blood pressure and resting heart rate were measured and venous blood taken during the visit. Medical history and current health status are recorded according to ICD-10 codes.

**GeneSTAR (Genetic Studies of Atherosclerosis Risk):** GeneSTAR is a family-based prospective study of more than 4000 participants begun in 1983 to determine phenotypic and genetic causes of premature cardiovascular disease. Families were identified from 1983-2006 from probands with a premature coronary disease event prior to 60 years of age who were identified at the time of hospitalization in any of 10 hospitals in the Baltimore, Maryland area. Their apparently healthy 30-59 year old siblings without known coronary disease were recruited and screened between 1983 and 2006. From 2003-2006, adult offspring over 21 years of age of all participating siblings and probands, as well as the coparents of the offspring were recruited and screened. Genotyping was performed in 3,232 participants on the Illumina 1Mv1_c platform.

**HCHS/SOL (Hispanic Community Health Study/ Study of Latinos):** The HCHS/SOL is a community-based cohort study of 16,415 self-identified Hispanic/Latino persons aged 18–74 years and selected from households in predefined census-block groups across four US field centers (in Chicago, Miami, the Bronx, and San Diego). The census-block groups were chosen to provide diversity among cohort participants with regard to socioeconomic status and national origin or background. The HCHS/SOL cohort includes participants who self-identified as having a Hispanic/Latino background; the largest groups are Central American (n = 1,730), Cuban (n = 2,348), Dominican (n = 1,460), Mexican (n = 6,471), Puerto Rican (n = 2,728), and South American (n = 1,068). The HCHS/SOL baseline clinical examination occurred between 2008 and 2011 and included comprehensive biological, behavioral, and sociodemographic assessments. Consenting HCHS/SOL subjects were genotyped at Illumina on the HCHS/SOL custom 15041502 B3 array. The custom array comprised the Illumina Omni 2.5M array (HumanOmni2.5-8v.1-1) ancestry-informative markers, known GWAS hits and drug absorption, distribution, metabolism, and excretion (ADME) markers, and additional custom content including ~150,000 SNPs selected from the CLM (Colombian in Medellin, Colombia), MXL (Mexican Ancestry in Los Angeles, California), and PUR (Puerto Rican in Puerto Rico) samples in the 1000Genomes phase 1 data to capture a greater amount of Amerindian genetic variation. QA/QC procedures yielded a total of 12,803 unique study participants for imputation and downstream association analyses.

**IRAS Family Study (Insulin Resistance Atherosclerosis Study):** The IRASFS was a family study designed to examine the genetic and epidemiologic basis of glucose homeostasis traits and abdominal adiposity. Briefly, self-reported Mexican American pedigrees were recruited in San Antonio, TX and San Luis Valley, CO. Probands with large families were recruited from the initial non-family-based IRAS, which was modestly enriched for impaired glucose tolerance and T2D. Inclusion of IRASFS data is limited to 1040 normoglycemic individuals in 88 pedigrees with genotype data from the Illumina OmniExpress and Omni 1S arrays and imputation to the 1000 Genome Integrated Reference Panel (phase I).

**KORA (Cooperative Health Research in the Augsburg Region):** The KORA study is a series of independent population-based epidemiological surveys of participants living in the region of Augsburg, Southern Germany. All survey participants are residents of German nationality identified through the registration office and were examined in 1994/95 (KORA S3) and 1999/2001 (KORA S4). In the KORA S3 and S4 studies 4,856 and 4,261 subjects have been examined implying response rates of 75% and 67%, respectively. 3,006 subjects participated in a 10-year follow-up examination of S3 in 2004/05 (KORA F3), and 3080 of S4 in 2006/2008 (KORA F4). The age range of the participants was 25 to 74 years at recruitment. Informed consent has been given by all participants. The study has been approved by the local ethics committee. Individuals for genotyping in KORA F3 and KORA F4 were randomly selected and these genotypes are taken for the analysis of the phenotypes in KORA S3 and KORA S4.

**Lifelines (Netherlands Biobank):** Lifelines (https://lifelines.nl/) is a multi-disciplinary prospective population-based cohort study using a unique three-generation design to examine the health and health-related behaviors of 165,000 persons living in the North East region of The Netherlands. It employs a broad range of investigative procedures in assessing the biomedical, socio-demographic, behavioral, physical and psychological factors which contribute to the health and disease of the general population, with a special focus on multimorbidity. In addition, the Lifelines project comprises a number of cross-sectional sub-studies which investigate specific age-related conditions. These include investigations into metabolic and hormonal diseases, including obesity, cardiovascular and renal diseases, pulmonary diseases and allergy, cognitive function and depression, and musculoskeletal conditions. All survey participants are between 18 and 90 years old at the time of enrollment. Recruitment has been going on since the end of 2006, and over 130,000 participants had been included by April 2013. At the baseline examination, the participants in the study were asked to fill in a questionnaire (on paper or online) before the first visit. During the first and second visit, the first or second part of the questionnaire, respectively, are checked for completeness, a number of investigations are conducted, and blood and urine samples are taken. Lifelines is a facility that is open for all researchers. Information on application and data access procedure is summarized on [www.lifelines.nl](http://www.lifelines.nl).

Scholtens S, Smidt N, Swertz MA, Bakker SJ, Dotinga A, Vonk JM, et al. Cohort Profile: LifeLines, a three-generation cohort study and biobank. Int J Epidemiol. 2014 Dec 14.

**NESDA (Netherlands Study of Depression and Anxiety):** NESDA is a multi-center study designed to examine the long-term course and consequences of depressive and anxiety disorders (http://www.nesda.nl)^1^. NESDA included both individuals with depressive and/or anxiety disorders and controls without psychiatric conditions. Inclusion criteria were age 18-65 years and self-reported western European ancestry while exclusion criteria were not being fluent in Dutch and having a primary diagnosis of another psychiatric condition (psychotic disorder, obsessive compulsive disorder, bipolar disorder, or severe substance use disorder).

Reference: Penninx, B.W. et al. The Netherlands Study of Depression and Anxiety (NESDA): rationale, objectives and methods. Int J Methods Psychiatr Res 17, 121-40 (2008).

**SHIP (Study of Health in Pomerania):** The Study of Health In Pomerania (SHIP) is a prospective longitudinal population-based cohort study in Mecklenburg-West Pomerania assessing the prevalence and incidence of common diseases and their risk factors (PMID: 20167617). SHIP encompasses the two independent cohorts SHIP and SHIP-TREND. Participants aged 20 to 79 with German citizenship and principal residency in the study area were recruited from a random sample of residents living in the three local cities, 12 towns as well as 17 randomly selected smaller towns. Individuals were randomly selected stratified by age and sex in proportion to population size of the city, town or small towns, respectively. A total of 4,308 participants were recruited between 1997 and 2001 in the SHIP cohort. Between 2008 and 2012 a total of 4,420 participants were recruited in the SHIP-TREND cohort. Individuals were invited to the SHIP study centre for a computer-assisted personal interviews and extensive physical examinations. The study protocol was approved by the medical ethics committee of the University of Greifswald. Oral and written informed consents was obtained from each of the study participants

Genome-wide SNP-typing was performed using the Affymetrix Genome-Wide Human SNP Array 6.0 or the Illumina Human Omni 2.5 array (SHIP-TREND samples). Array processing was carried out in accordance with the manufacturer’s standard recommendations. Genotypes were determined using GenomeStudio Genotyping Module v1.0 (GenCall) for SHIP-TREND and the Birdseed2 clustering algorithm for SHIP. Imputation of genotypes in SHIP and SHIP-TREND was performed with the software IMPUTE v2.2.2 based on 1000 Genomes release March 2012.

**SWHS/SMHS (Shanghai Women's Health Study/** **Shanghai Men's Health Study):** The Shanghai Women's Health Study (SWHS) is an ongoing population-based cohort study of approximately 75,000 women who were aged 40-70 years at study enrollment and resided in in urban Shanghai, China; 56,832 (75.8%) provided a blood samples. Recruitment for the SWHS was initiated in 1997 and completed in 2000. The self-administered questionnaire includes information on demographic characteristics, disease and surgery histories, personal habits (such as cigarette smoking, alcohol consumption, tea drinking, and ginseng use), menstrual history, residential history, occupational history, and family history of cancer.

The Shanghai Men’s Health Study (SMHS) is an ongoing population-based cohort study of 61,480 Chinese men who were aged between 40 and 74 years, were free of cancer at enrollment, and lived in urban Shanghai, China; 45,766 (74.4%) provided a blood samples. Recruitment for the SMHS was initiated in 2002 and completed in 2006. The self-administered questionnaire includes information on demographic characteristics, disease and surgery histories, personal habits (such as cigarette smoking, alcohol consumption, tea drinking, and ginseng use), residential history, occupational history, and family history of cancer.

Genotyping and imputation: Genomic DNA was extracted from buffy coats by using a Qiagen DNA purification kit (Valencia, CA) or Puregene DNA purification kit (Minneapolis, MN) according to the manufacturers’ instructions and then used for genotyping assays. The GWAS genotyping was performed using the Affymetrix Genome-Wide Human SNP Array 6.0 (Affy6.0) platform or Illumina 660, following manufacturers’ protocols. After sample quality control, we exclude SNPs with 1) MAF <0.01; 2) call rate <95%; 2) bad genotyping cluster; and 3) concordance rate <95% among duplicated QC samples. Genotypes were imputed using the program MACH (http://www.sph.umich.edu/csg/abecasis/MACH/download/), which determines the probable distribution of missing genotypes conditional on a set of known haplotypes, while simultaneously estimating the fine-scale recombination map. Phased autosome SNP data from HapMap Phase II Asians (release 22) were used as the reference. To test for associations between the imputed SNP data with BMI, linear regression (additive model) was used, in which SNPs were represented by the expected allele count, an approach that takes into account the degree of uncertainty of genotype imputation (http://www.sph.umich.edu/csg/abecasis/MACH/download/).

**YFS (The Cardiovascular Risk in Young Finns Study):** The YFS is a population-based follow up-study started in 1980. The main aim of the YFS is to determine the contribution made by childhood lifestyle, biological and psychological measures to the risk of cardiovascular diseases in adulthood. In 1980, over 3,500 children and adolescents all around Finland participated in the baseline study. The follow-up studies have been conducted mainly with 3-year intervals. The latest 30-year follow-up study was conducted in 2010-11 (ages 33-49 years) with 2,063 participants. The study was approved by the local ethics committees (University Hospitals of Helsinki, Turku, Tampere, Kuopio and Oulu) and was conducted following the guidelines of the Declaration of Helsinki. All participants gave their written informed consent.

**STAGE 2 STUDY ACKNOWLEDGMENTS:**

Infrastructure for the CHARGE Consortium is supported in part by the National Heart, Lung, and Blood Institute grant R01HL105756. Infrastructure for the Gene-Lifestyle Working Group is supported by the National Heart, Lung, and Blood Institute grant R01HL118305.

**Airwave (The Airwave Health Monitoring Study):** We thank all participants in the Airwave Health Monitoring Study. The study is funded by the Home Office (Grant number 780-TETRA) with additional support from the National Institute for Health Research (NIHR), Imperial College Healthcare NHS Trust (ICHNT) and Imperial College Biomedical Research Centre (BRC). The study has ethical approval from the National Health Service Multi-site Research Ethics Committee (MREC/13/NW/0588). This work used computing resources provided by the MRC- funded UK MEDical Bioinformatics partnership programme (UK MED-BIO) (MR/L01632X/1). P.E. would like to acknowledge support from the Medical Research Council and Public Health England for the MRC-PHE Centre for Environment and Health (MR/L01341X/1) and from the NIHR NIHR Health Protection Research Unit in Health Impact of Environmental Hazards (HPRU-2012-10141). P.E. is supported by the UK-DRI Dementia Research Institute at Imperial College London which receives funding from UK DRI Ltd funded by the UK Medical Research Council, Alzheimer’s Society and Alzheimer’s Research UK. P.E. is associate director of Health Data Research UK-London which receives funding from a consortium led by the UK Medical Research Council.

**CFS (Cleveland Family Study):** The CFS was supported by the National Institutes of Health, the National Heart, Lung, Blood Institute grant HL113338, R01HL098433, HL46380.

**CoLaus (Cohorte Lausannoise):** The CoLaus study was and is supported by research grants from GlaxoSmithKline, the Faculty of Biology and Medicine of Lausanne, and the Swiss National Science Foundation (grants 33CSCO-122661, 33CS30-139468 and 33CS30-148401).

**DFTJ (Dongfeng-Tongji Cohort Study):** This work was supported by grants from the Foundation of National Key Program of Research and Development of China (2016YFC0900800), the Programme of Introducing Talents of Discipline, the grants from the National Natural Science Foundation (grant NSFC-81473051, 81522040 and 81230069), and the Program for the New Century Excellent Talents in University (NCET-11-0169).

**DR’s EXTRA (Dose-Responses to Exercise Training):** The study was supported by grants from Ministry of Education and Culture of Finland (722 and 627; 2004-2010); Academy of Finland (102318, 104943, 123885, 211119); European Commission FP6 Integrated Project (EXGENESIS), LSHM-CT-2004-005272; City of Kuopio; Juho Vainio Foundation; Finnish Diabetes Association; Finnish Foundation for Cardiovascular Research; Kuopio University Hospital; Päivikki and Sakari Sohlberg Foundation; Social Insurance Institution of Finland 4/26/2010.

**EGCUT (Estonian Genome Center - University of Tartu (Estonian Biobank)):** This study was supported by EU H2020 grants 692145, 676550, 654248, Estonian Research Council Grant IUT20-60 and PUT1660, NIASC, EIT – Health and NIH-BMI Grant No: 2R01DK075787-06A1 and EU through the European Regional Development Fund (Project No. 2014-2020.4.01.15-0012 GENTRANSMED.

**GeneSTAR (Genetic Studies of Atherosclerosis Risk):** GeneSTAR was supported by National Institutes of Health grants from the National Heart, Lung, and Blood Institute (HL49762, HL59684, HL58625, HL071025, U01 HL72518, HL087698, HL092165, HL099747, and K23HL105897), National Institute of Nursing Research (NR0224103), National Institute of Neurological Disorders and Stroke (NS062059), and by grants from the National Center for Research Resources to the Johns Hopkins General Clinical Research Center (M01-RR000052) and the Johns Hopkins Institute for Clinical & Translational Research (UL1 RR 025005).

**HCHS/SOL (Hispanic Community Health Study/ Study of Latinos):** The baseline examination of HCHS/SOL was supported by contracts from the National Heart, Lung, and Blood Institute (NHLBI) to the University of North Carolina (N01-HC65233), University of Miami (N01-HC65234), Albert Einstein College of Medicine (N01-HC65235), Northwestern University (N01-HC65236), and San Diego State University (N01-HC65237). The National Institute on Minority Health and Health Disparities, National Institute on Deafness and Other Communication Disorders, National Institute of Dental and Craniofacial Research (NIDCR), National Institute of Diabetes and Digestive and Kidney Diseases (NIDDK), National Institute of Neurological Disorders and Stroke, and NIH Office of Dietary Supplements additionally contributed funding to HCHS/SOL. The Genetic Analysis Center at the University of Washington was supported by NHLBI and NIDCR contracts (HHSN268201300005C AM03 and MOD03). Additional analysis support was provided by 1R01DK101855-01 and 13GRNT16490017. Genotyping was also supported by National Center for Advancing Translational Sciences UL1TR000124 and NIDDK DK063491 to the Southern California Diabetes Endocrinology Research Center. This research was also supported in part by the Intramural Research Program of the NIDDK, contract no. HHSB268201200054C, and Illumina.

**IRAS Family Study (Insulin Resistance Atherosclerosis Study):** The IRASFS is supported by the National Heart Lung and Blood Institute (HL060944, HL061019, and HL060919). Genotyping for this study was supported by the GUARDIAN Consortium with grant support from the National Institute of Diabetes and Digestive and Kidney Diseases (NIDDK; DK085175) and in part by UL1TR000124 (CTSI) and DK063491 (DRC). The authors thank study investigators, staff, and participants for their valuable contributions.

**KORA (Cooperative Health Research in the Augsburg Region):** The KORA study was initiated and financed by the Helmholtz Zentrum München – German Research Center for Environmental Health, which is funded by the German Federal Ministry of Education and Research (BMBF) and by the State of Bavaria. Furthermore, KORA research was supported within the Munich Center of Health Sciences (MC-Health), Ludwig-Maximilians-Universität, as part of LMUinnovativ. This work was further supported by a grant (WA 4081/1-1) from the German Research Foundation.

**LifeLines (Netherlands Biobank):** The Lifelines Cohort Study, and generation and management of GWAS genotype data for the Lifelines Cohort Study is supported by the Netherlands Organization of Scientific Research NWO (grant 175.010.2007.006), the Economic Structure Enhancing Fund (FES) of the Dutch government, the Ministry of Economic Affairs, the Ministry of Education, Culture and Science, the Ministry for Health, Welfare and Sports, the Northern Netherlands Collaboration of Provinces (SNN), the Province of Groningen, University Medical Center Groningen, the University of Groningen, Dutch Kidney Foundation and Dutch Diabetes Research Foundation.

The authors wish to acknowledge the services of the Lifelines Cohort Study, the contributing research centers delivering data to Lifelines, and all the study participants.

**NESDA (Netherlands Study of Depression and Anxiety):** Funding was obtained from the Netherlands Organization for Scientific Research (Geestkracht program grant 10-000-1002); the Center for Medical Systems Biology (CSMB, NWO Genomics), Biobanking and Biomolecular Resources Research Infrastructure (BBMRI-NL), VU University’s Institutes for Health and Care Research (EMGO+) and Neuroscience Campus Amsterdam, University Medical Center Groningen, Leiden University Medical Center, National Institutes of Health (NIH, R01D0042157-01A, MH081802, Grand Opportunity grants 1RC2 MH089951 and 1RC2 MH089995). Part of the genotyping and analyses were funded by the Genetic Association Information Network (GAIN) of the Foundation for the National Institutes of Health.Computing was supported by BiG Grid, the Dutch e-Science Grid, which is financially supported by NWO.

**SHIP (Study of Health in Pomerania):** SHIP is part of the Community Medicine Research net of the University of Greifswald, Germany, which is funded by the Federal Ministry of Education and Research (grants no. 01ZZ9603, 01ZZ0103, and 01ZZ0403), the Ministry of Cultural Affairs as well as the Social Ministry of the Federal State of Mecklenburg-West Pomerania, and the network ‘Greifswald Approach to Individualized Medicine (GANI_MED)’ funded by the Federal Ministry of Education and Research (grant 03IS2061A). Genome-wide data were supported by the Federal Ministry of Education and Research (grant no. 03ZIK012) and a joint grant from Siemens Healthcare, Erlangen, Germany and the Federal State of Mecklenburg- West Pomerania. The University of Greifswald is a member of the ‘Center of Knowledge Interchange’ program of the Siemens AG.

**SWHS/SMHS (Shanghai Women's Health Study/** **Shanghai Men's Health Study):** We thank all the individuals who took part in these studies and all the researchers who have enabled this work to be carried out. The Shanghai Women’s Health Study and the Shanghai Men’s Health Study are supported by research grants UM1CA182910 and UM1CA173640 from the U.S. National Cancer Institute, respectively.

**YFS (The Cardiovascular Risk in Young Finns Study):** The Young Finns Study has been financially supported by the Academy of Finland: grants 286284, 134309 (Eye), 126925, 121584, 124282, 129378 (Salve), 117787 (Gendi), and 41071 (Skidi); the Social Insurance Institution of Finland; Kuopio, Tampere and Turku University Hospital Medical Funds (grant X51001); Juho Vainio Foundation; Paavo Nurmi Foundation; Finnish Foundation for Cardiovascular Research ; Finnish Cultural Foundation; Tampere Tuberculosis Foundation; Emil Aaltonen Foundation; Yrjö Jahnsson Foundation; Signe and Ane Gyllenberg Foundation; and Diabetes Research Foundation of Finnish Diabetes Association.

The expert technical assistance in the statistical analyses by Leo-Pekka Lyytikäinen and Irina Lisinen is gratefully acknowledged.

**Supplementary Figure 1: Regional plots of the replicated novel hits of the joint effect of long total sleep time on HDL cholesterol in the multi-ancestry meta-analyses**

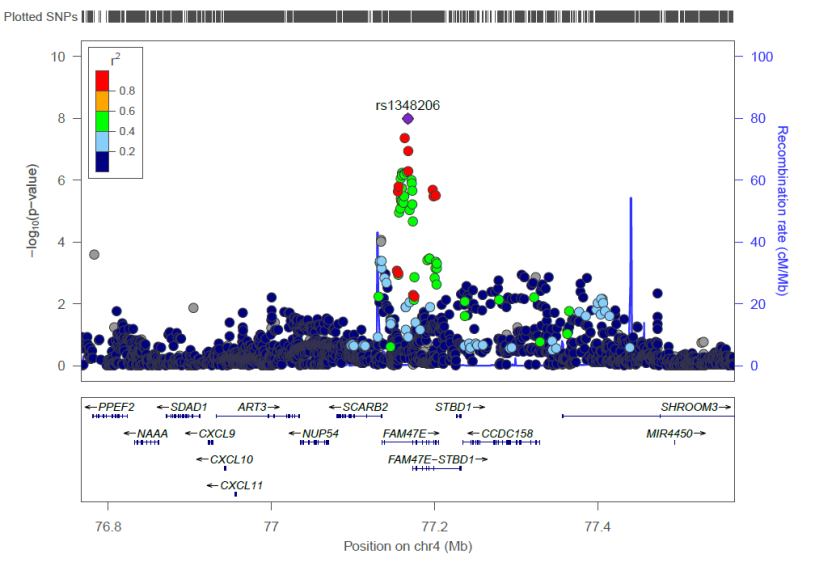

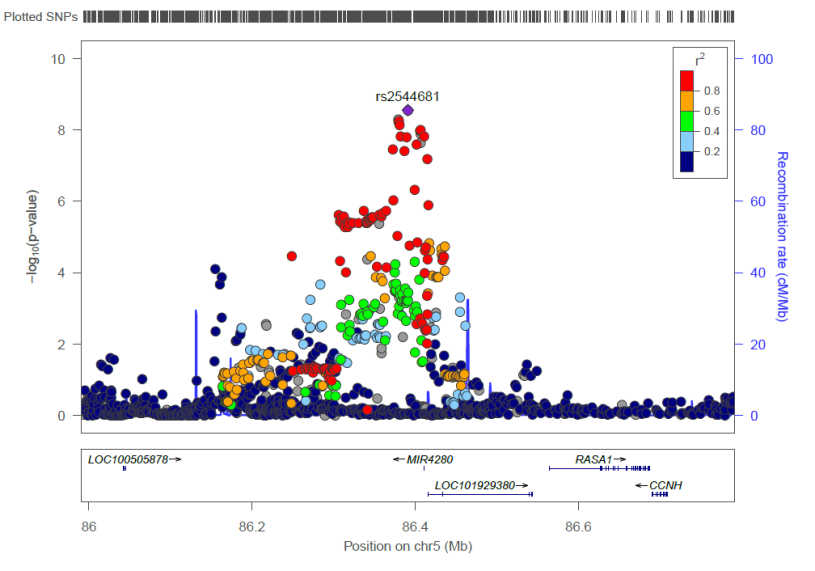

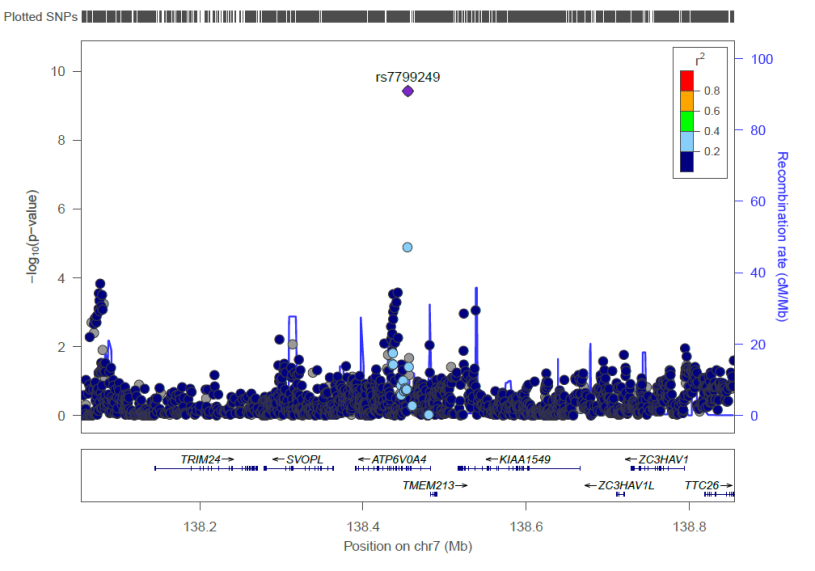

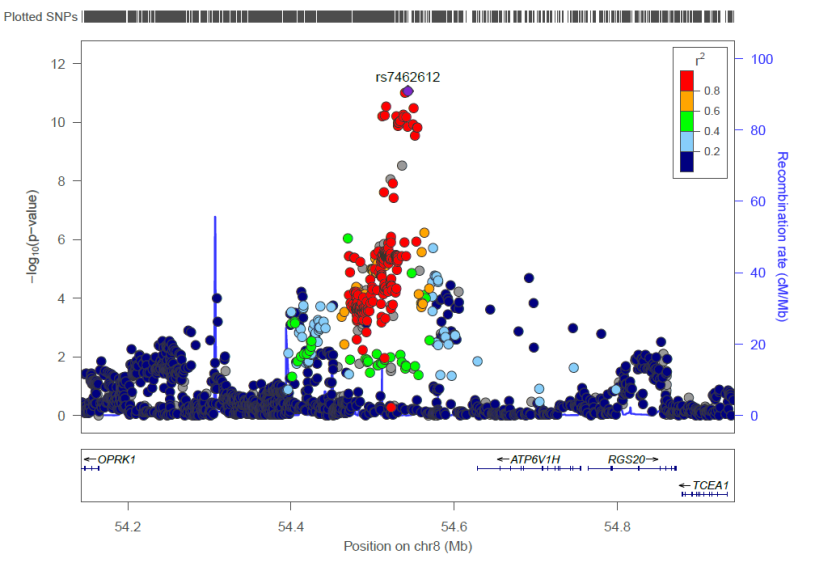

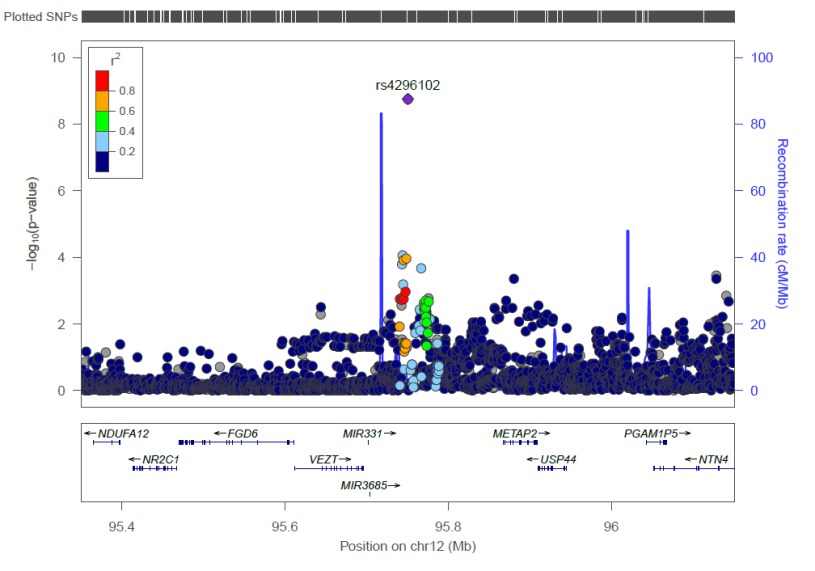

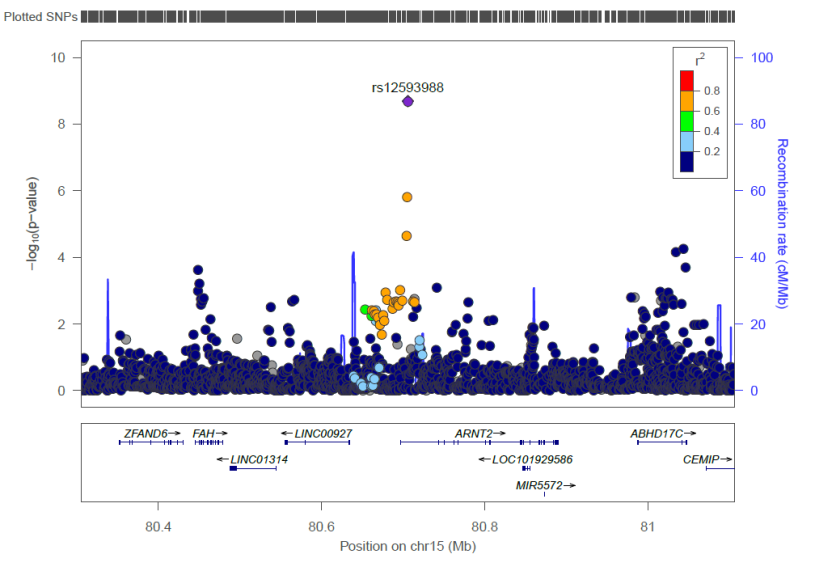

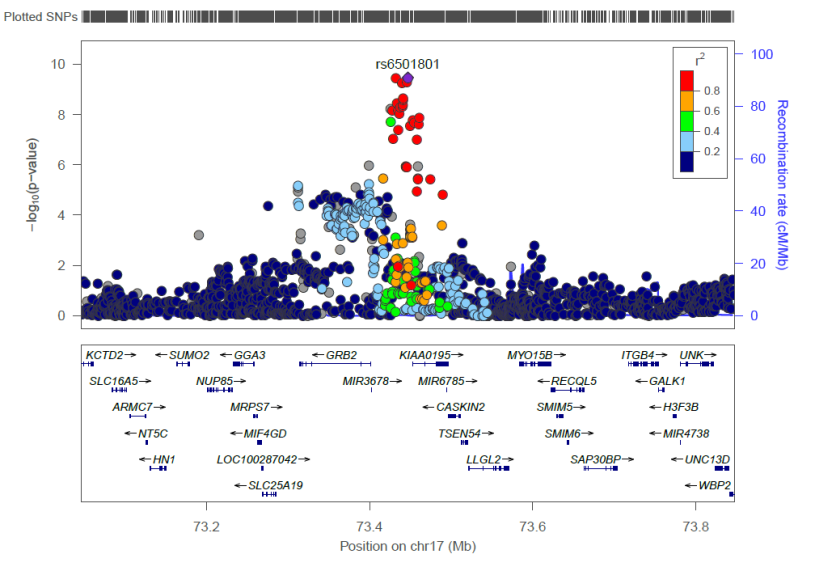

**Supplementary Figure 2: Regional plots of the replicated novel hits of the joint effect of short total sleep time on HDL cholesterol in the multi-ancestry meta-analyses**

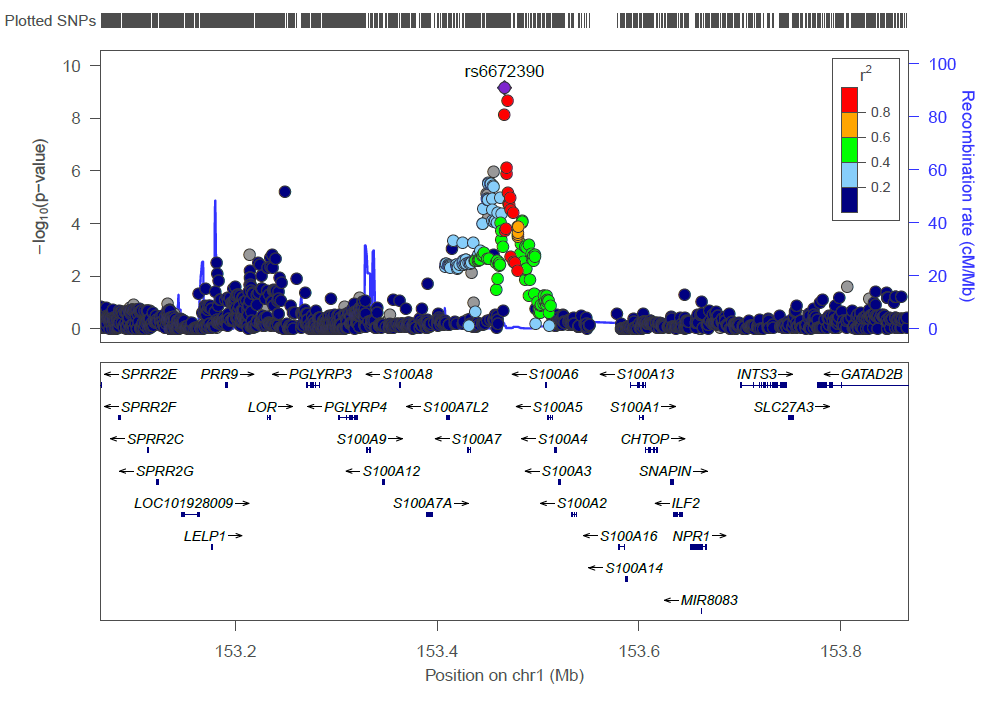

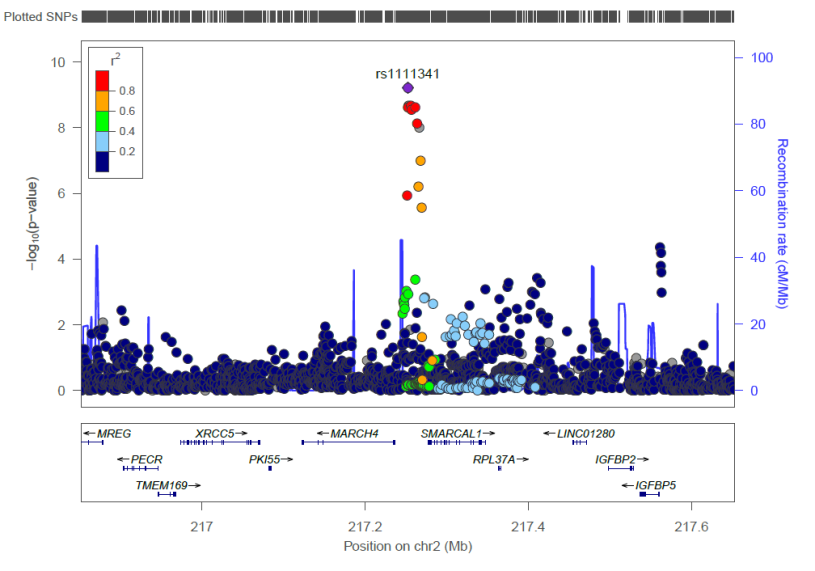

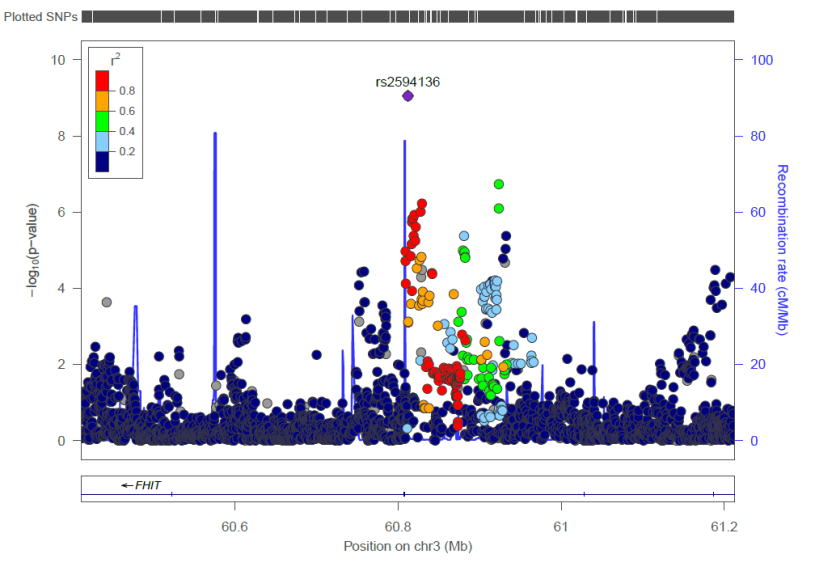

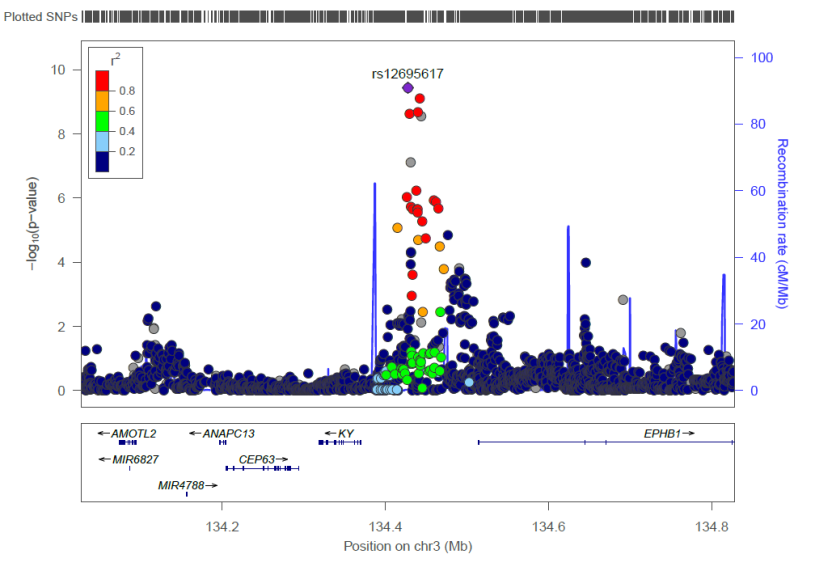

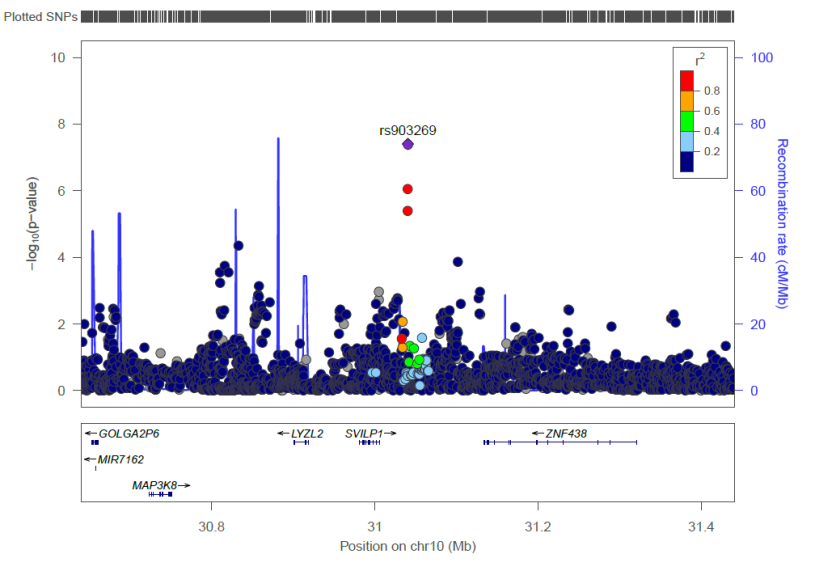

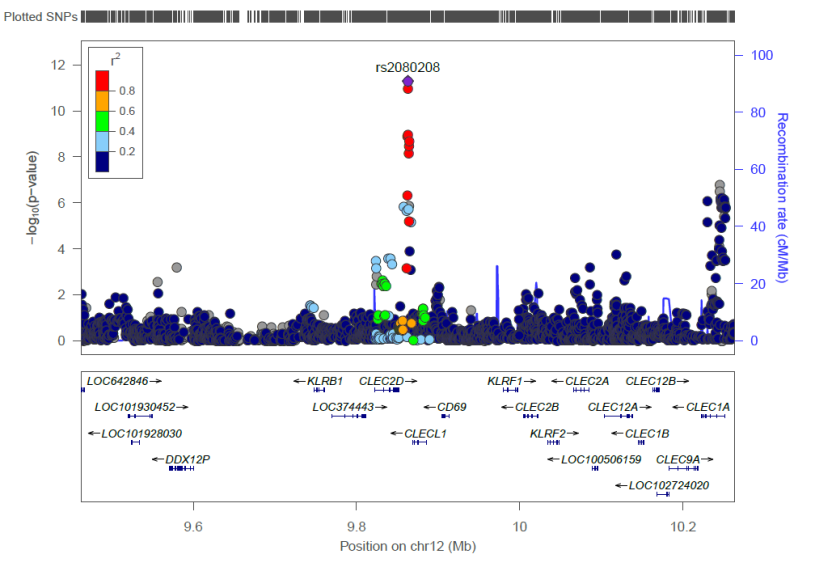

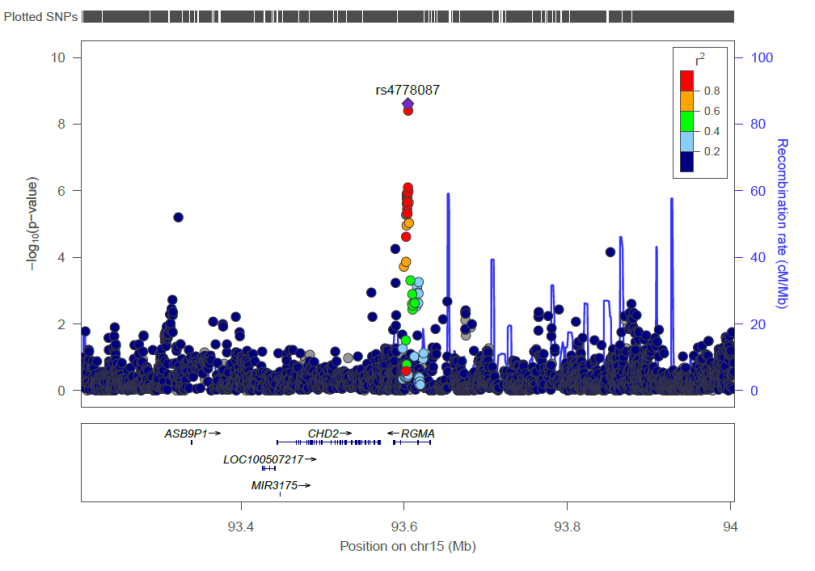

**Supplementary Figure 3: Regional plots of the replicated novel hits of the joint effect of long total sleep time on LDL cholesterol in the multi-ancestry meta-analyses**

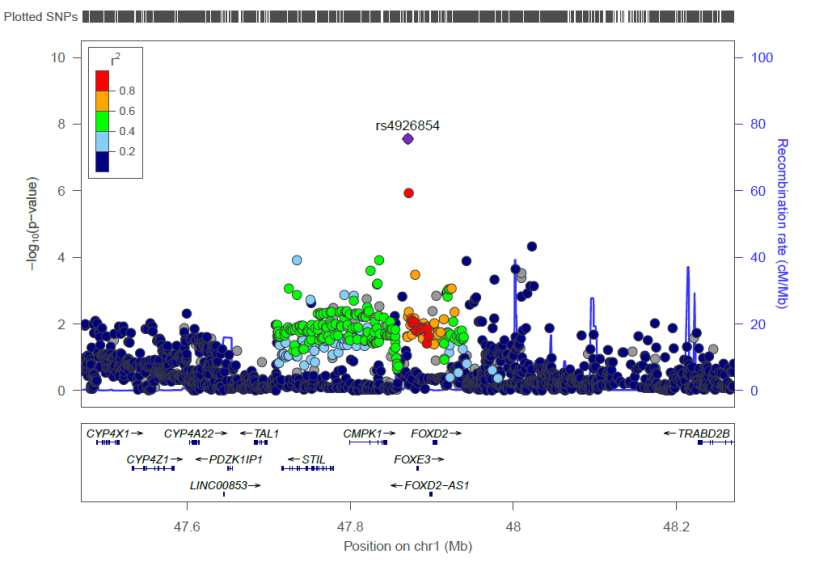

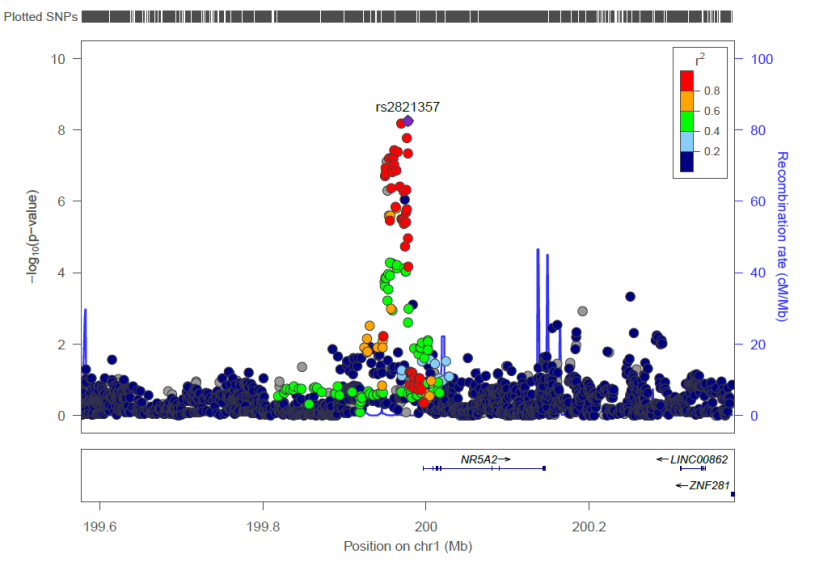

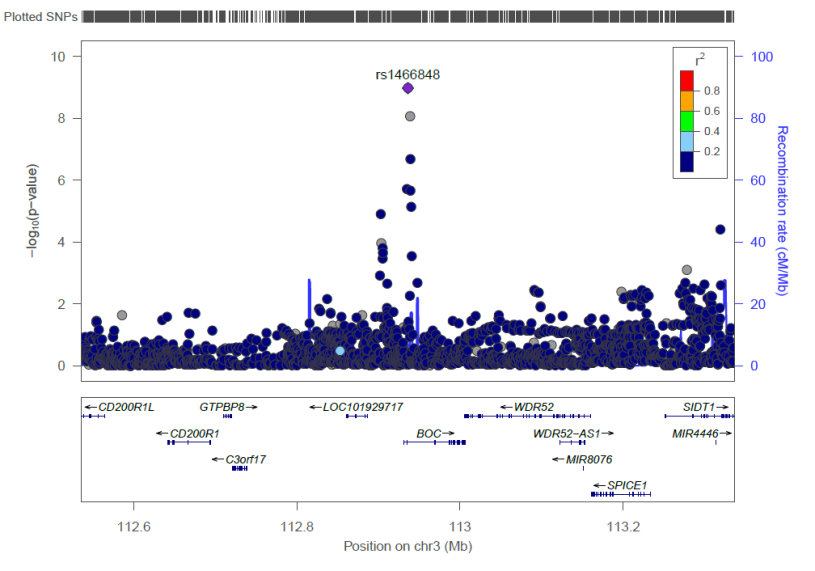

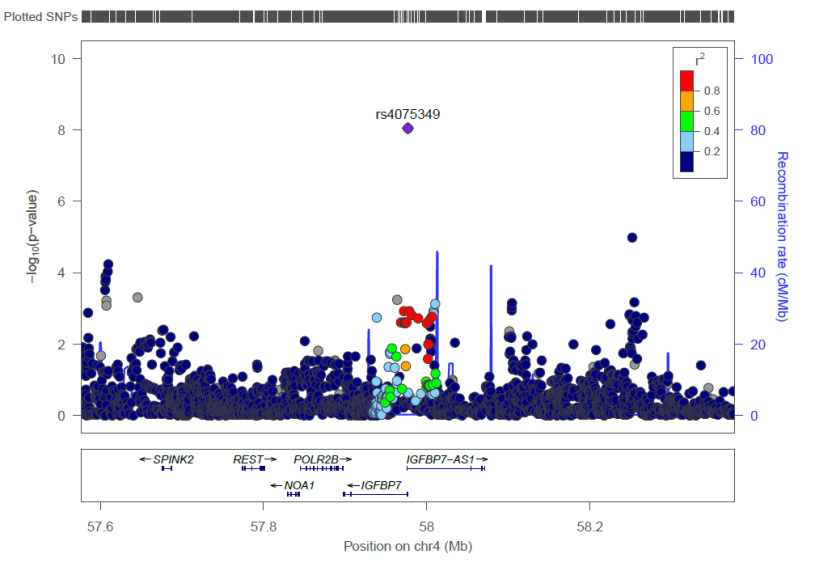

**Supplementary Figure 4: Regional plots of the replicated novel hits of the joint effect of short total sleep time on LDL cholesterol in the multi-ancestry meta-analyses**

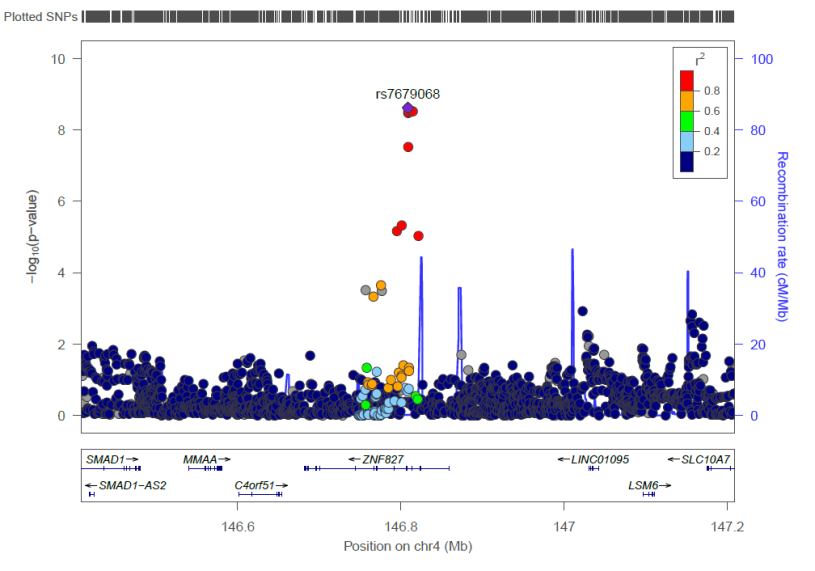

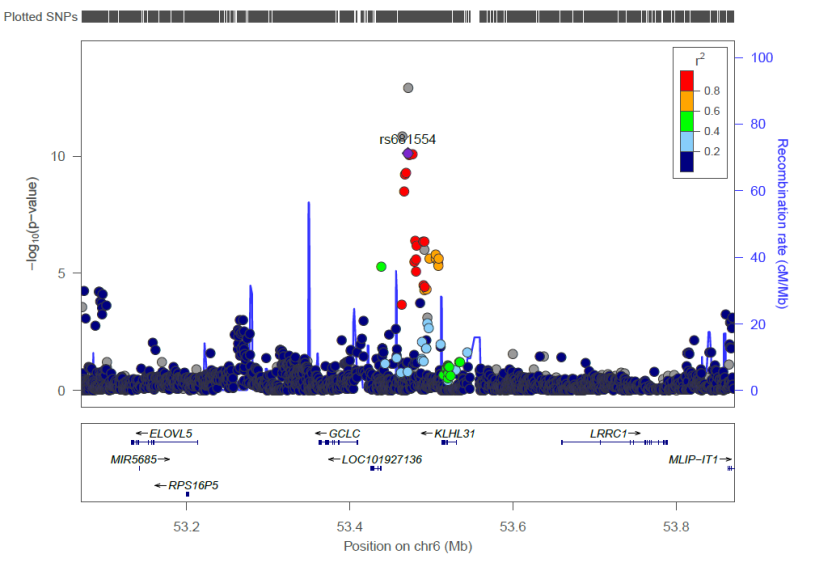

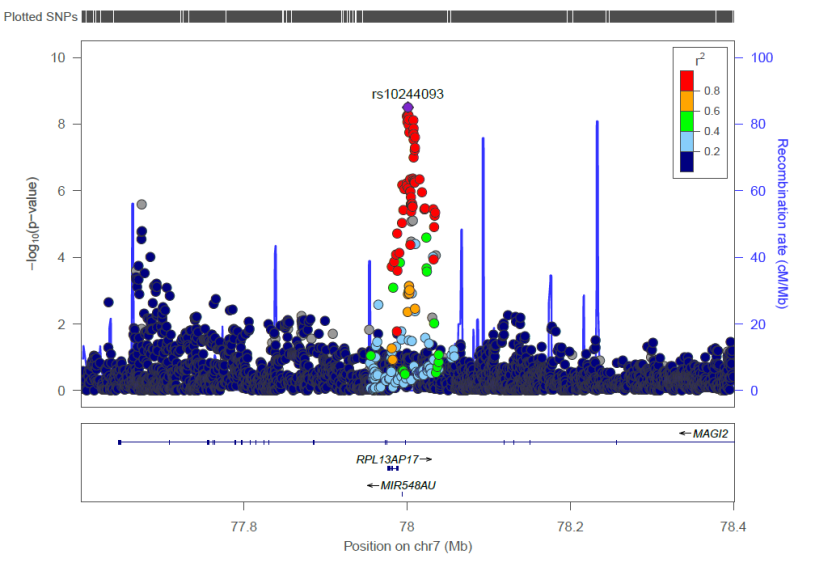

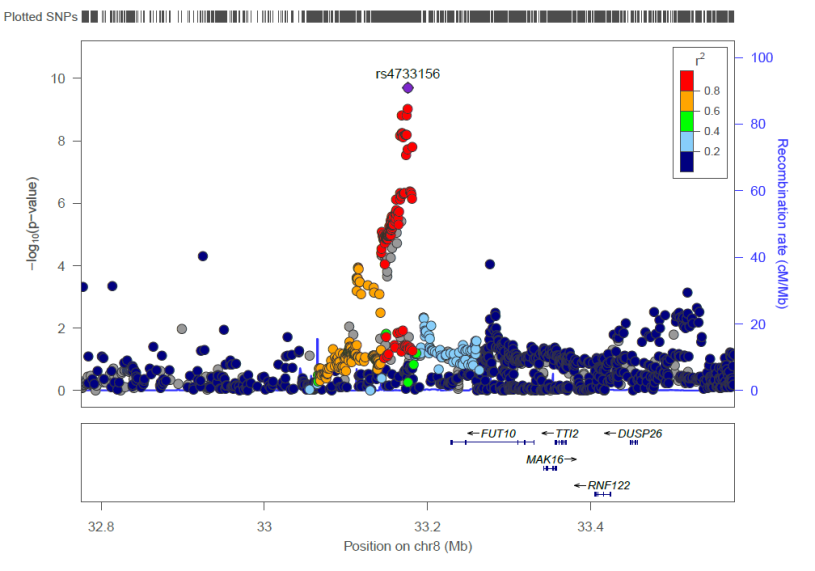

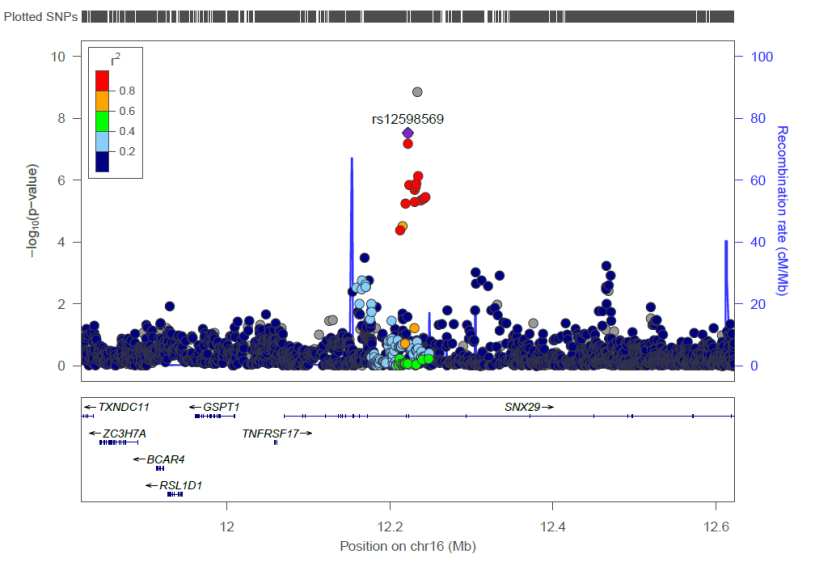

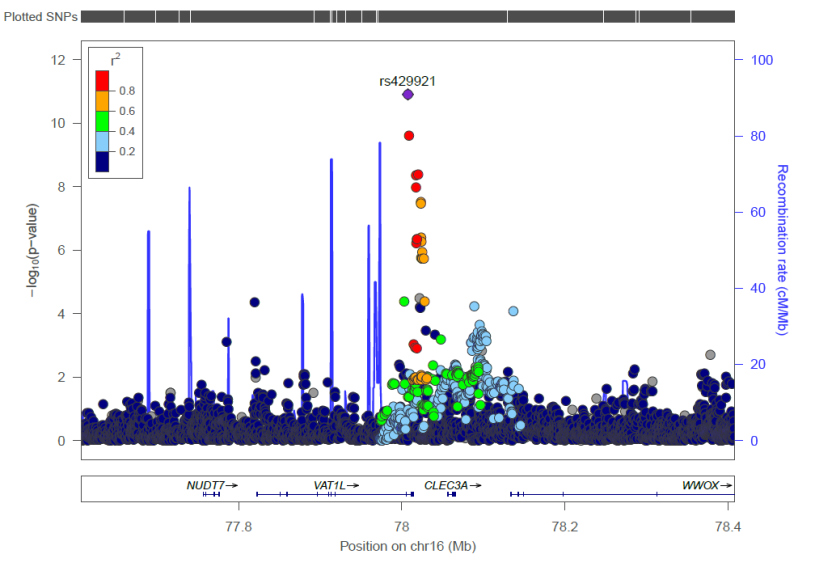

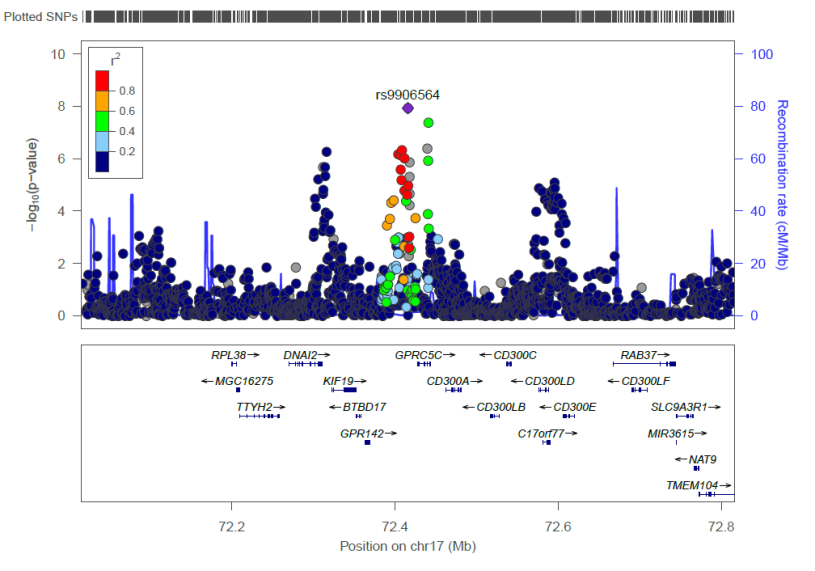

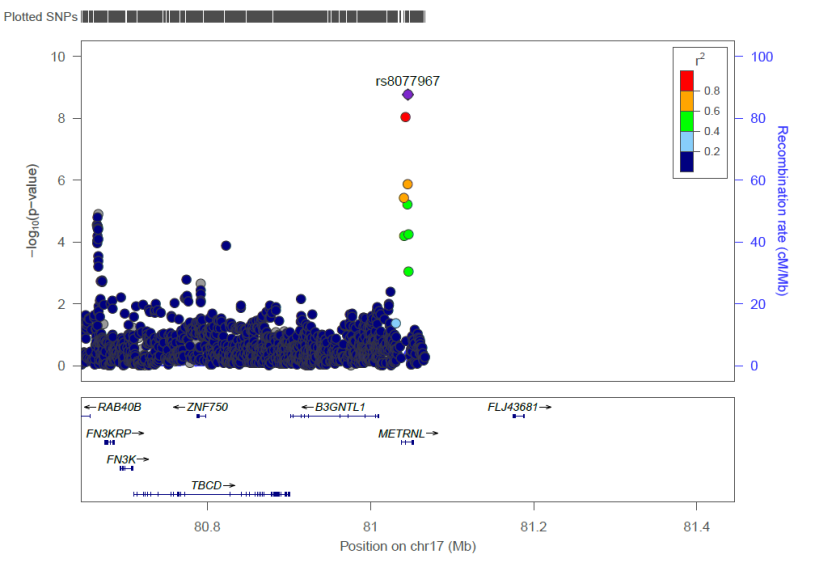

**Supplementary Figure 5: Regional plots of the replicated novel hits of the joint effect of long total sleep time on Triglycerides in the multi-ancestry meta-analyses**

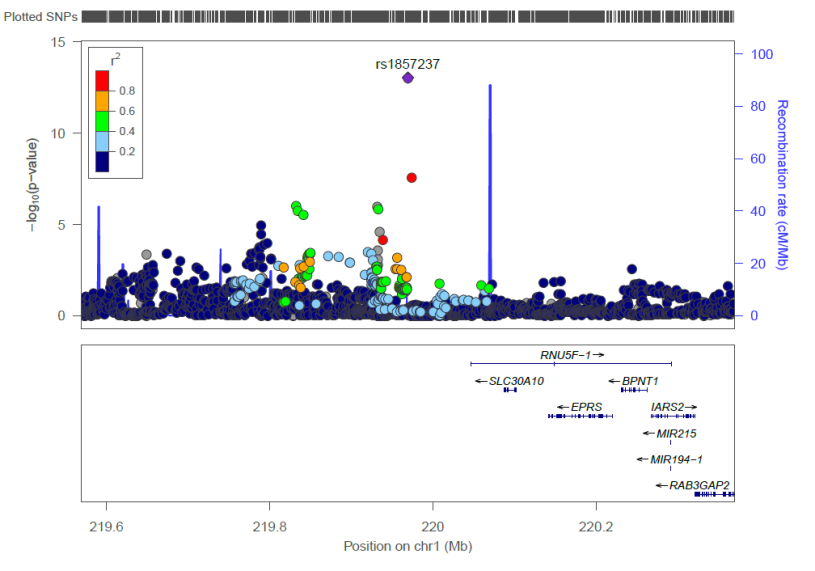

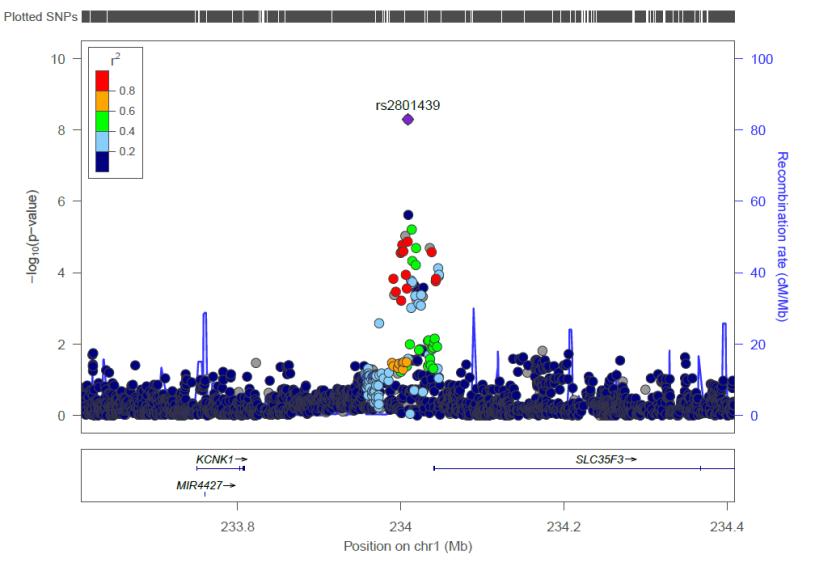

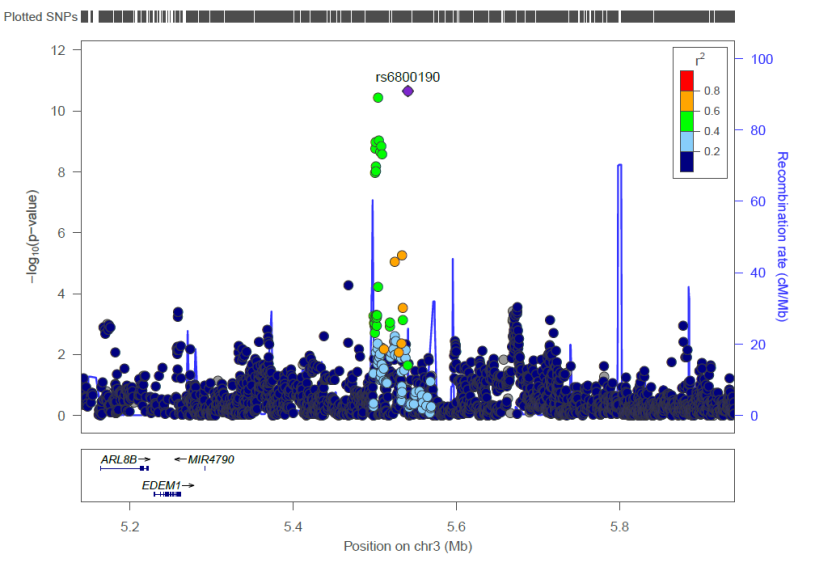

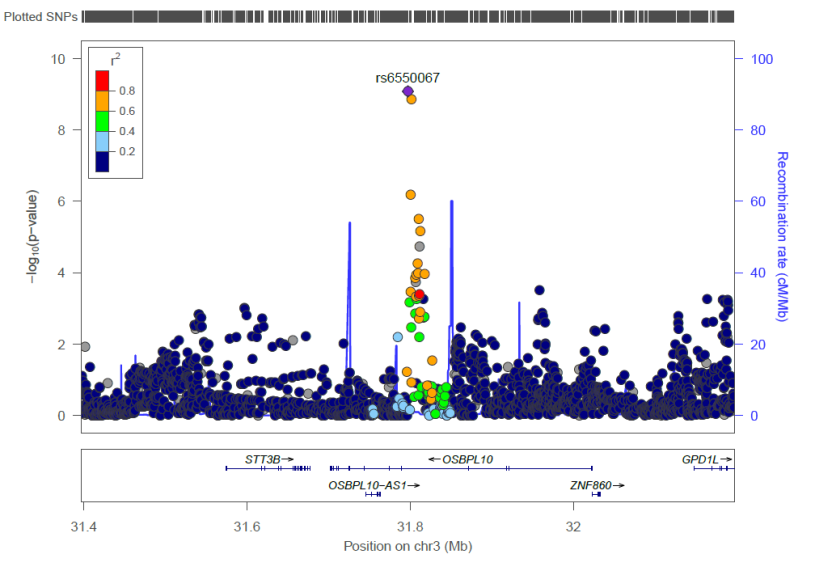

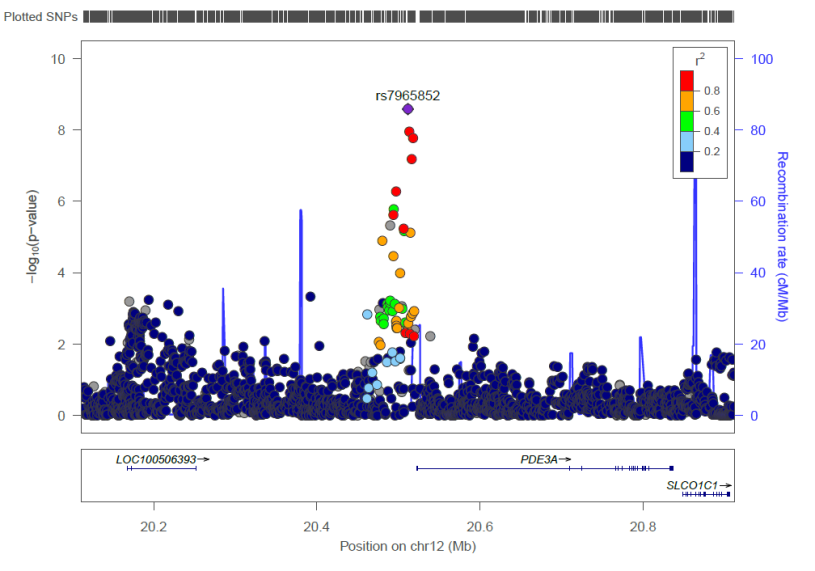

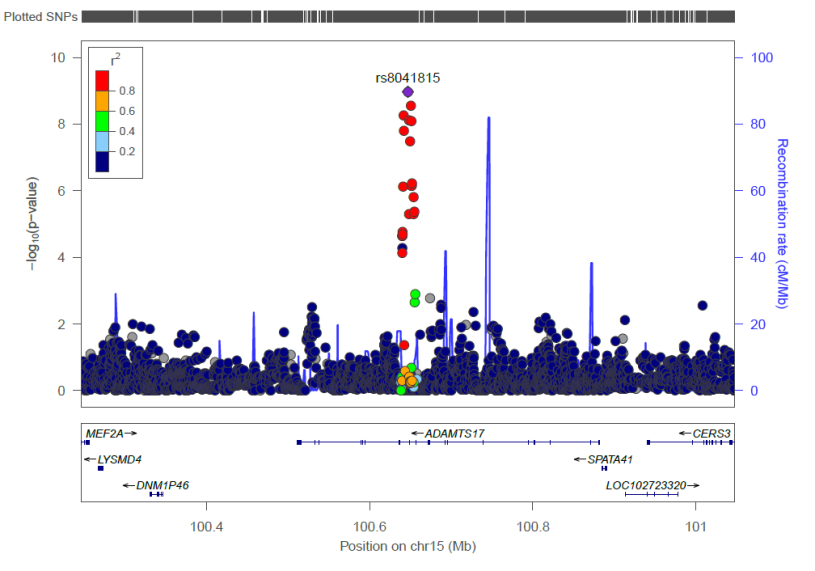

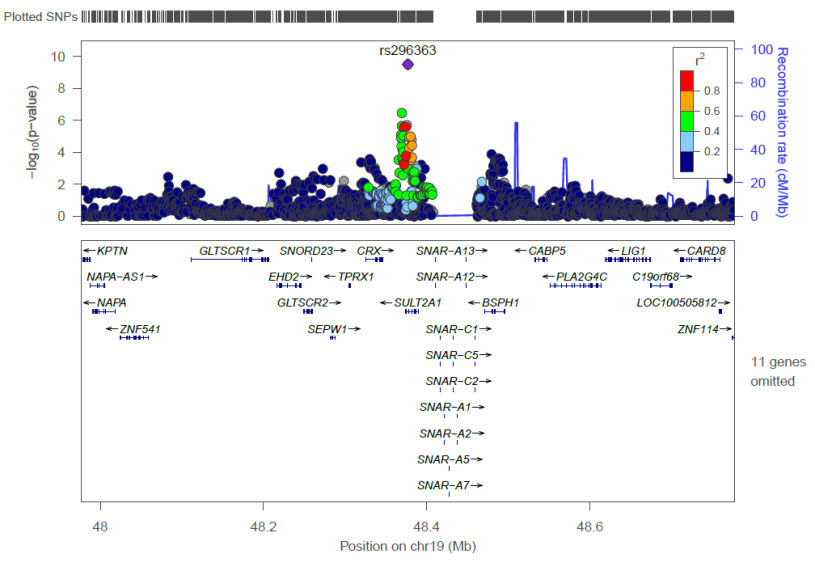

**Supplementary Figure 6: Regional plots of the replicated novel hits of the joint effect of short total sleep time on Triglycerides in the multi-ancestry meta-analyses**

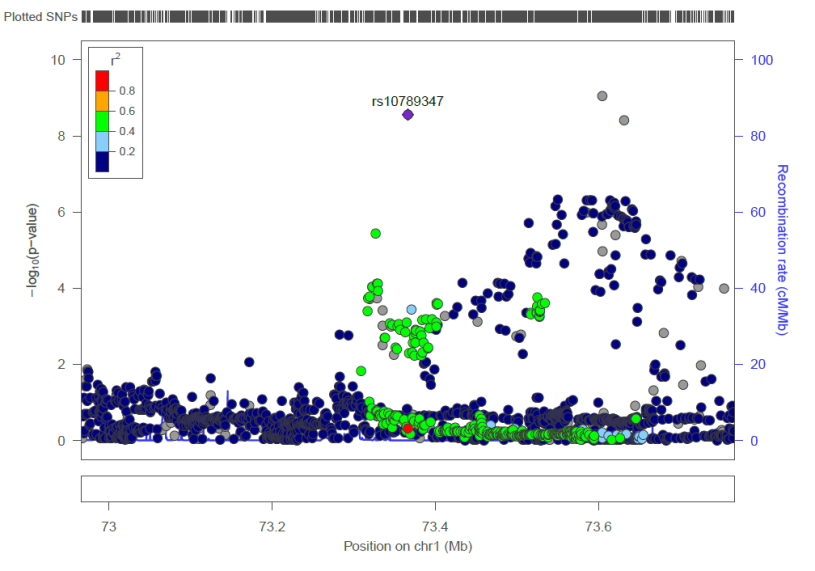

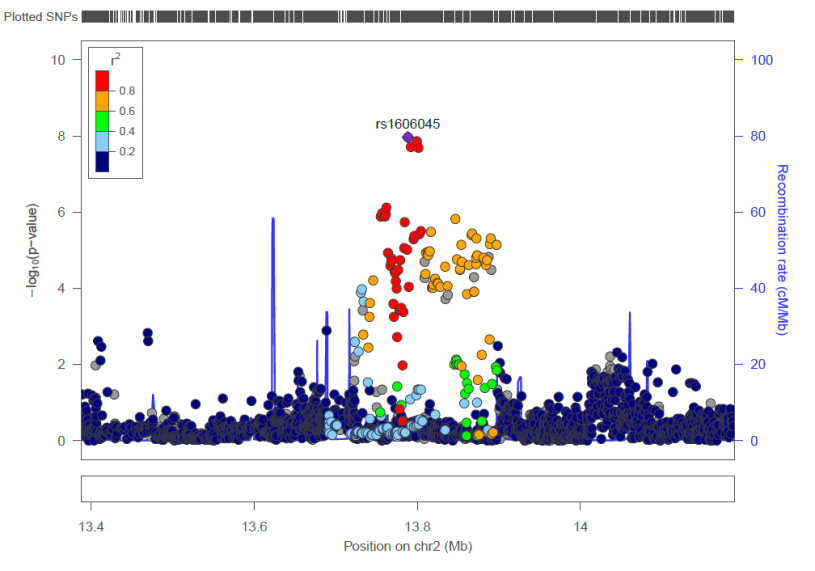

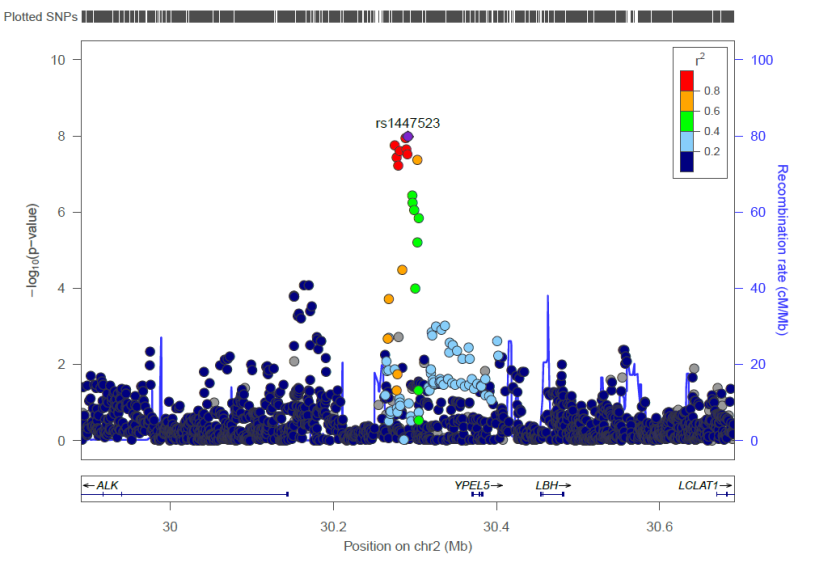

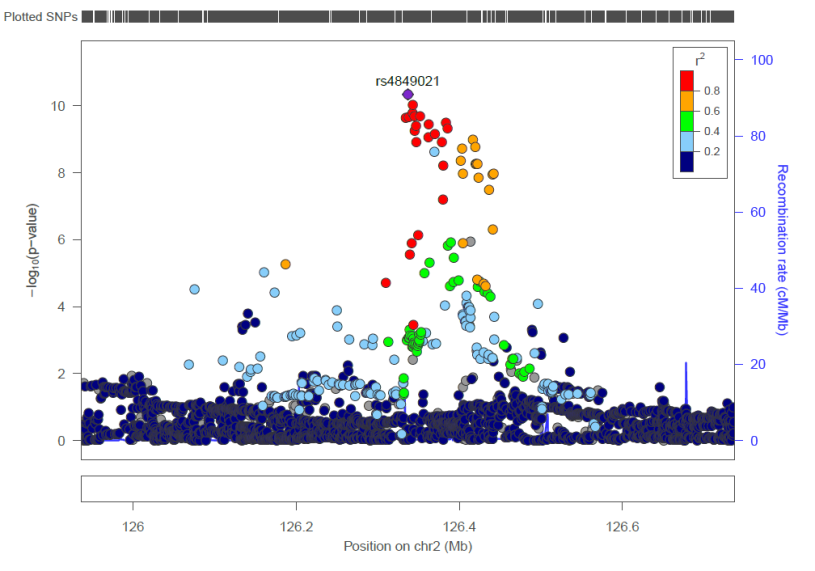

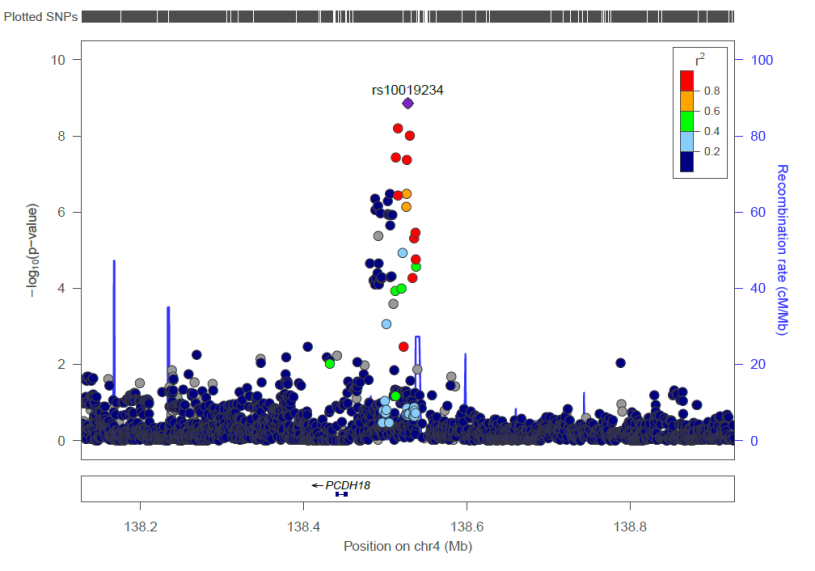

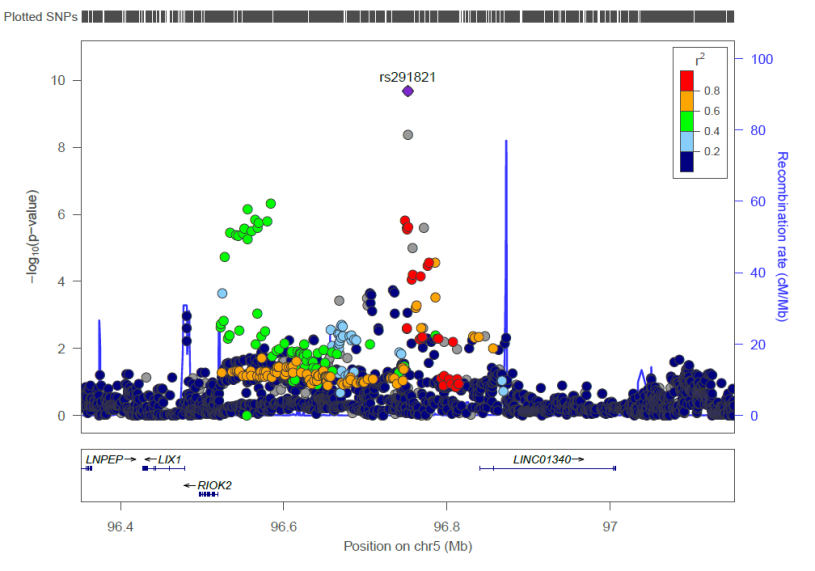

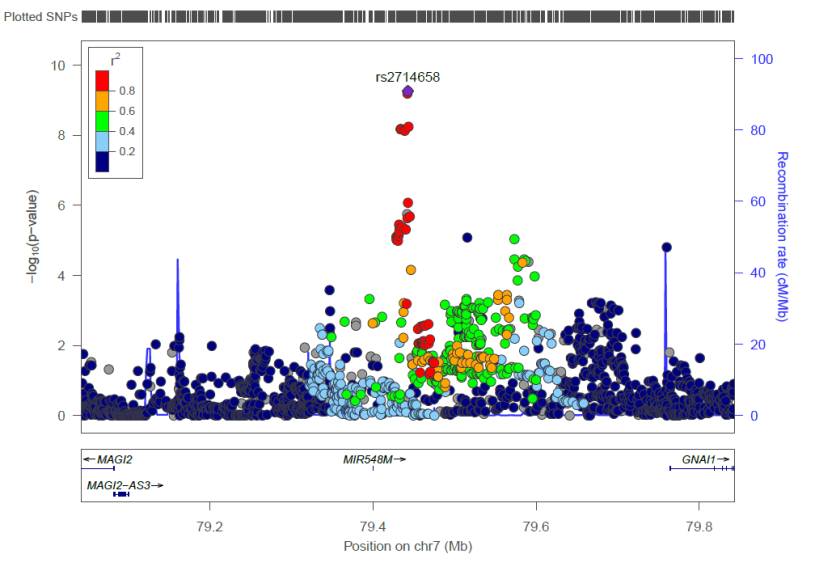

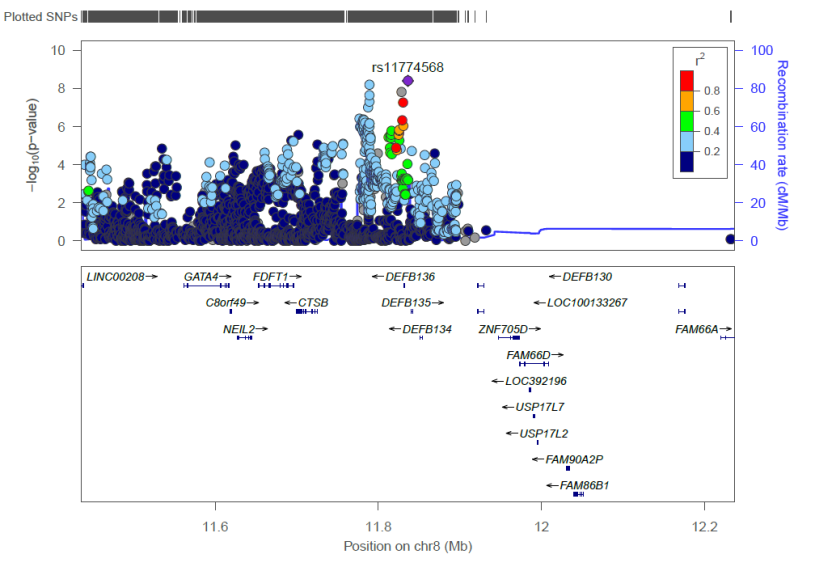

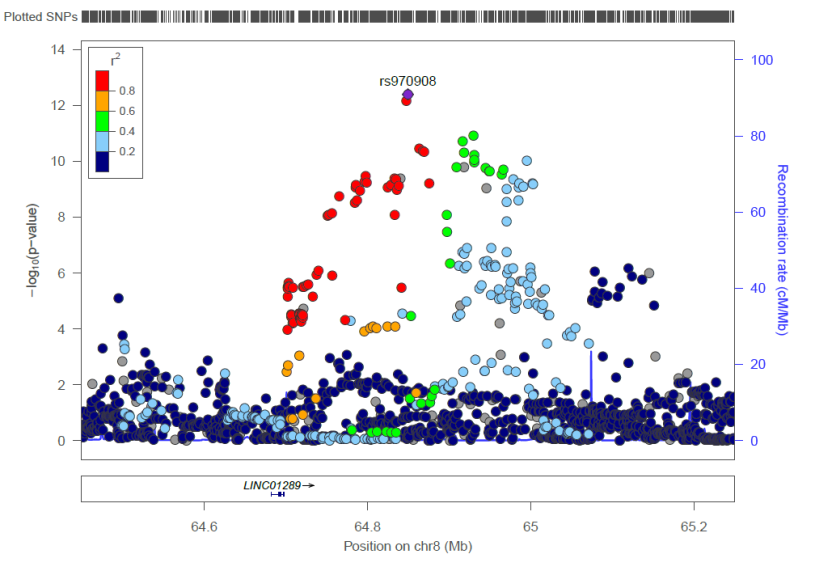

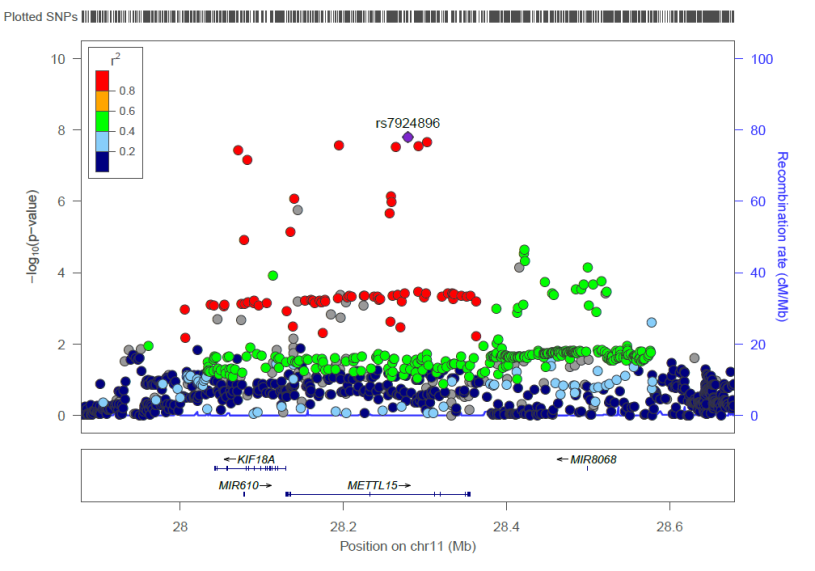

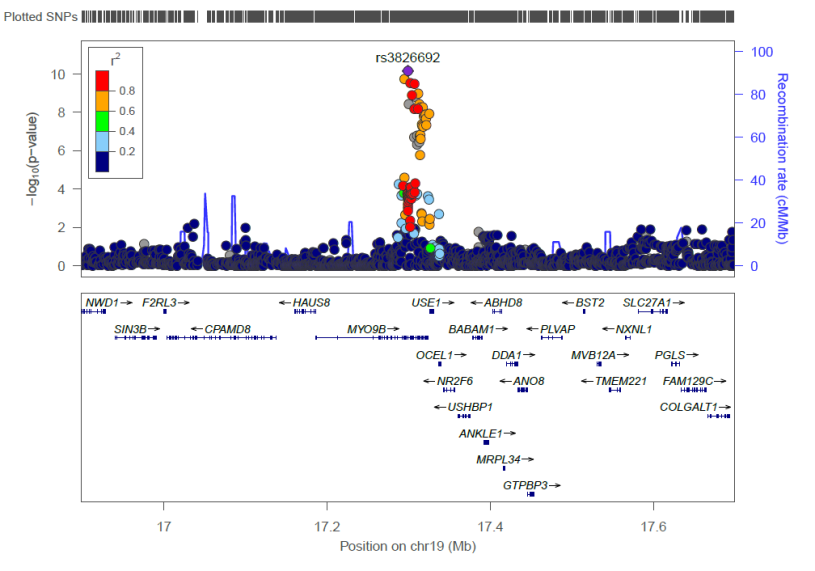

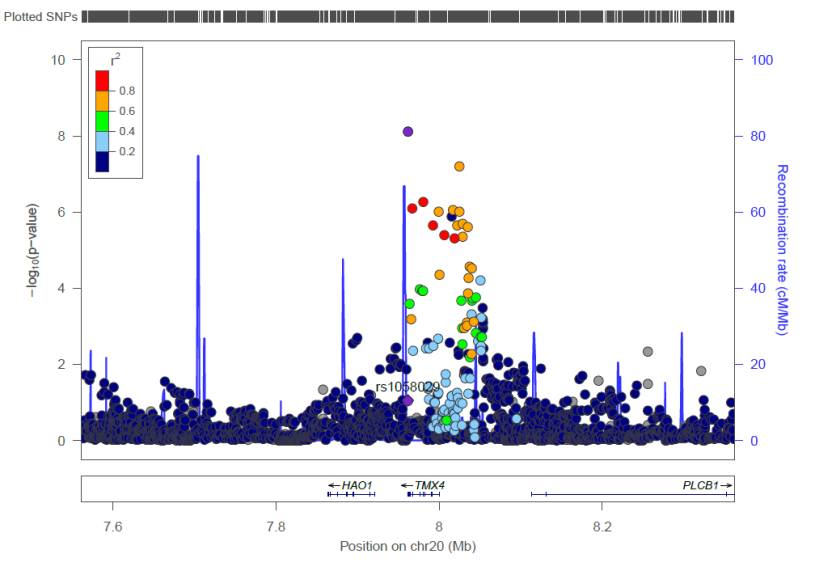

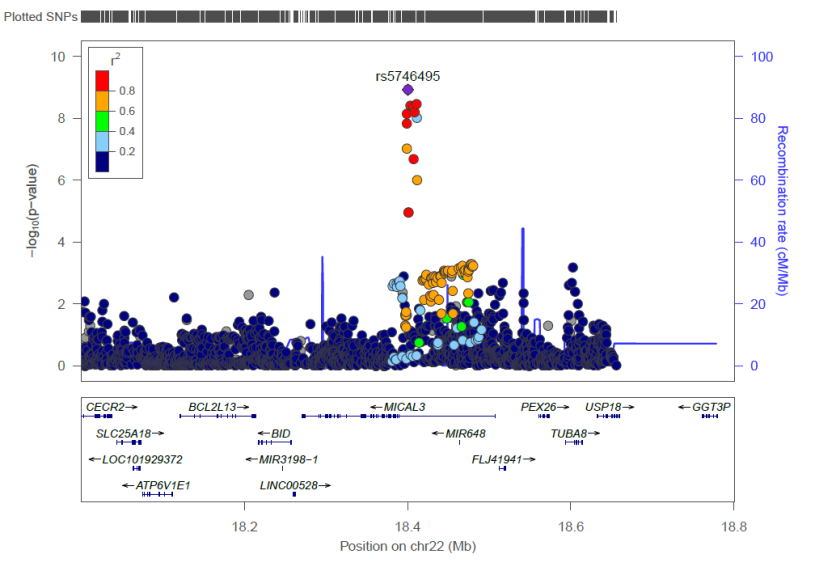

**Supplementary Figure 7: Project overview and SNP selection in the European-ancestry analyses**

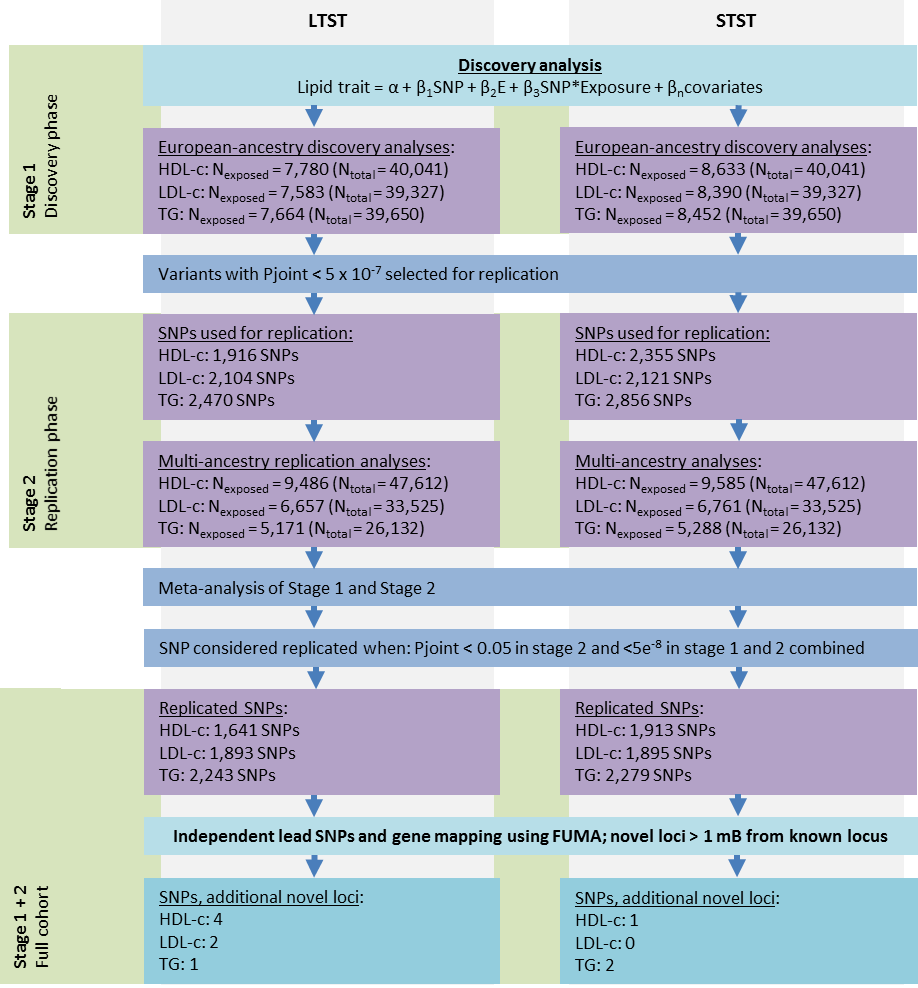

Project overview of the multi-ancestry analyses of how the new lipid loci were identified in the present project. Replicated variants had to have a Pjoint < 5x10^-7^ in stage 1, Pjoint < 0.05 in stage 2, and Pjoint <5x10^-8^ in the combined stage 1 + 2.

**Supplementary Figure 8: Circular –log(p-value) plots of the European-ancestry sleep-SNP interactions analyses for the three lipid traits**

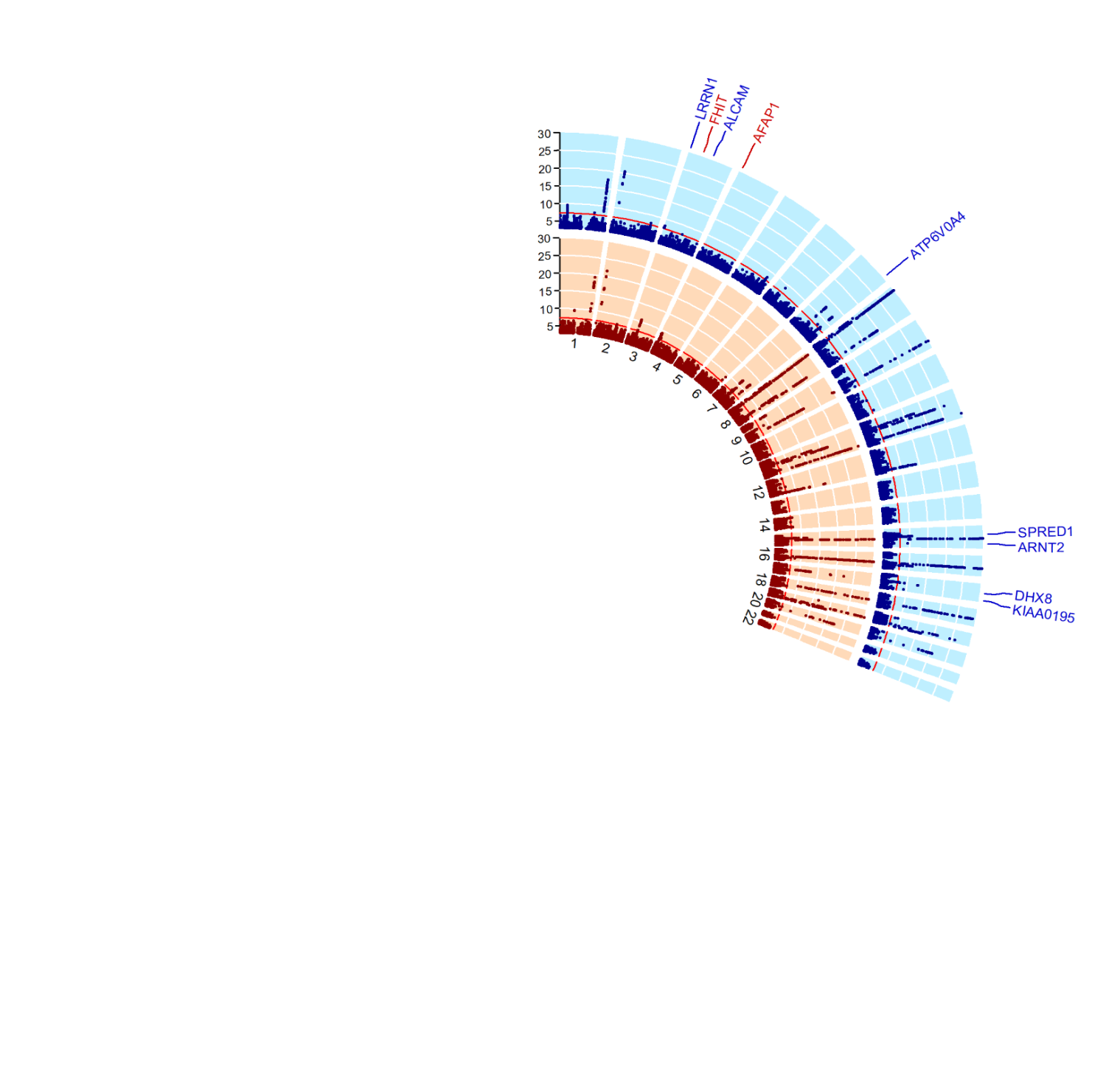

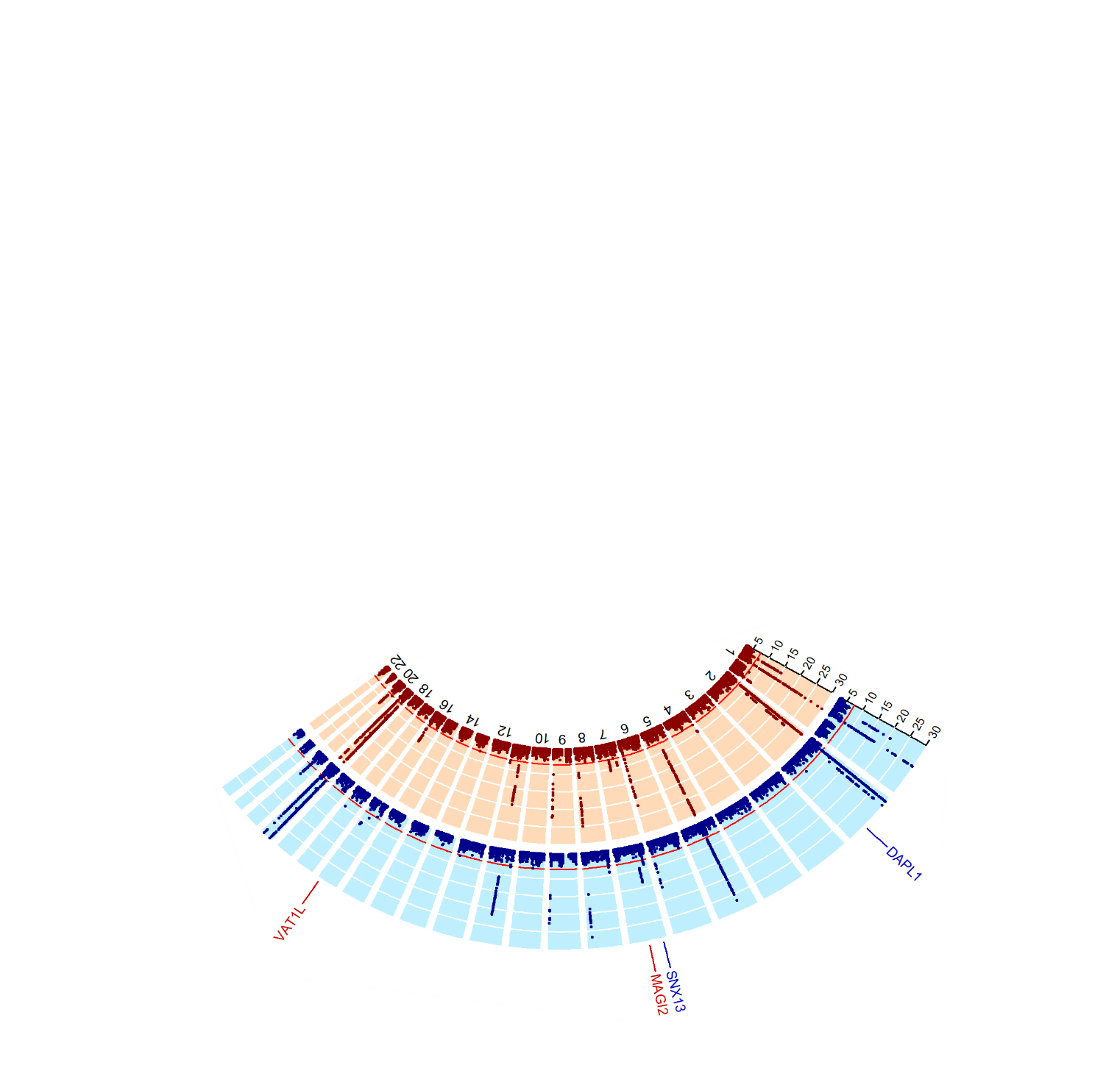

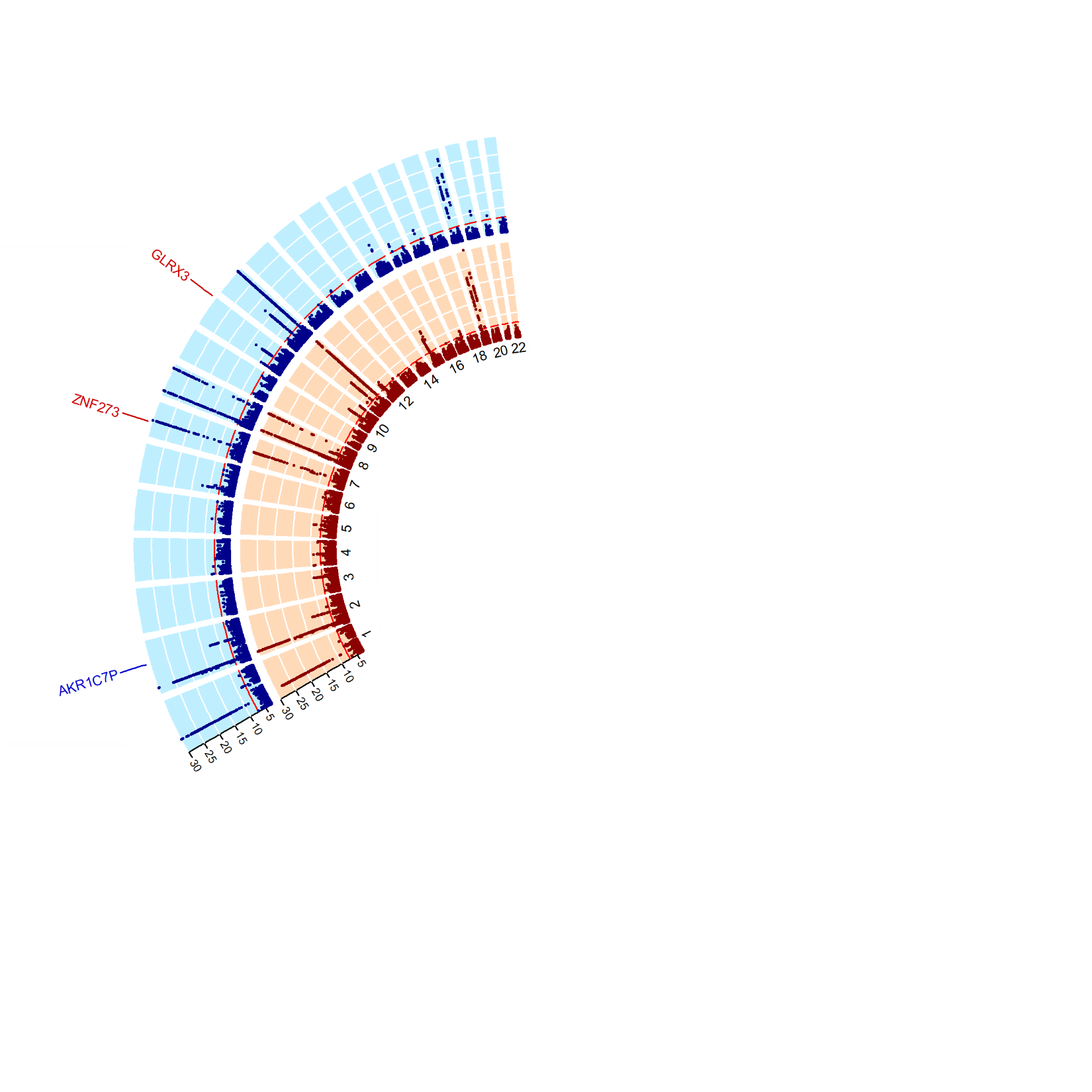

TG

HDL-c

LDL-c

Plot visualizes the –log(pjoints) for HDL-c, LDL-c and TG per chromosome. In red (inner circle) are the –log(p-value) plots for the analyses taking into account potential interaction with short total sleep time. In blue (outer circle) are the –log(p-value plots for the analyses taking into account potential interaction with long total sleep time. Loci defined as novel and being replicated are labeled. Replicated variants had to have a Pjoint < 5x10^-7^ in stage 1, Pjoint < 0.05 in stage 2, and Pjoint <5x10^-8^ in the combined stage 1 + 2. Labeled gene names in red were identified in the STST analysis; Labeled gene names in blue were identified in the LTST analysis. –log(p-values) > 30 were truncated for presentation purposes only. All –log(p_joints_) > 30 were truncated to 30 for visualization purposes only. Figure prepared using the R package circlize^38^.

**Supplementary Figure 9: Regional plots of the replicated additional novel hits of the joint effect of long total sleep time on HDL cholesterol in the European meta-analyses**

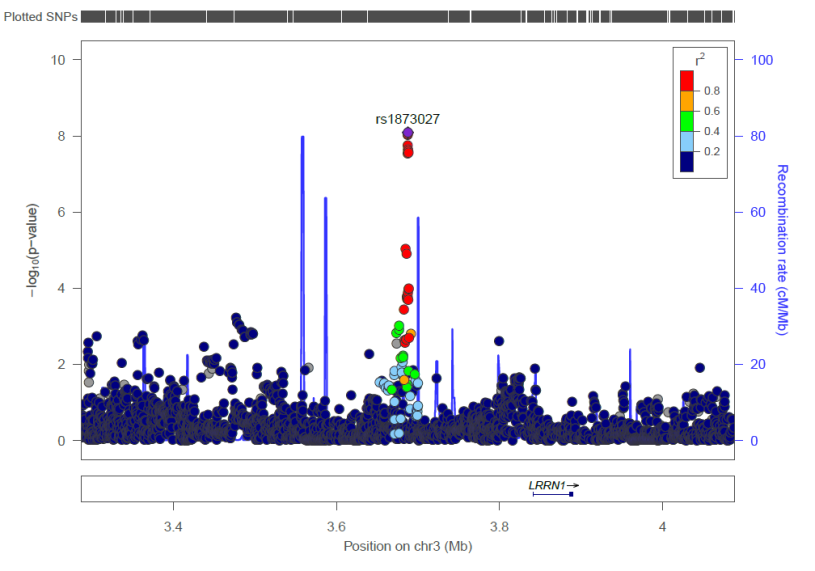

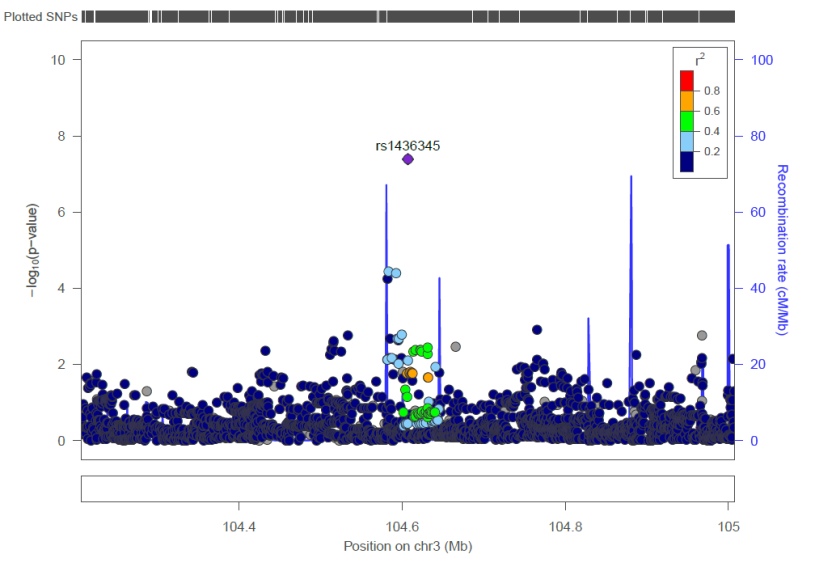

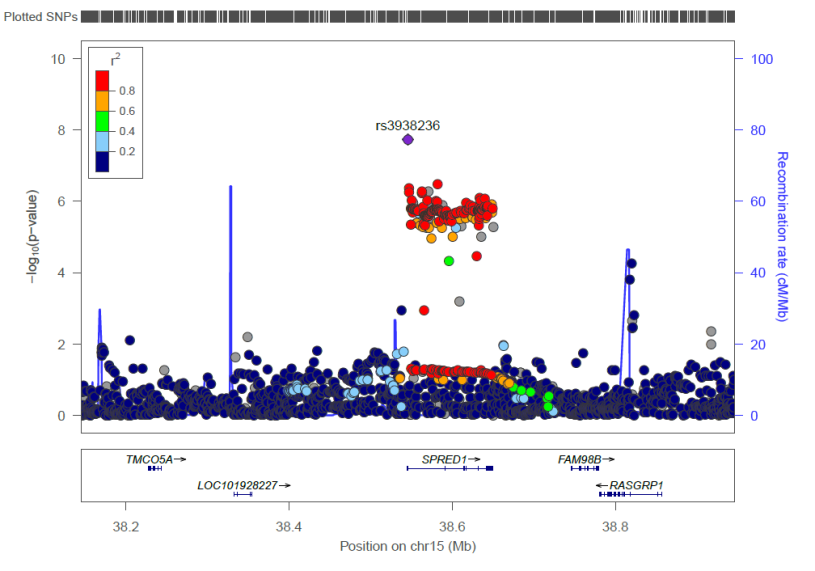

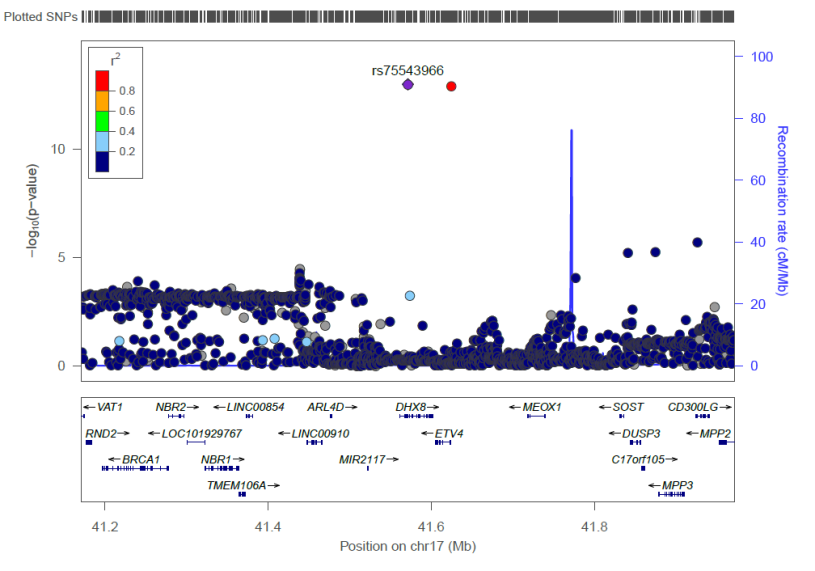

**Supplementary Figure 10: Regional plots of the replicated additional novel hits of the joint effect of short total sleep time on HDL cholesterol in the European meta-analyses**
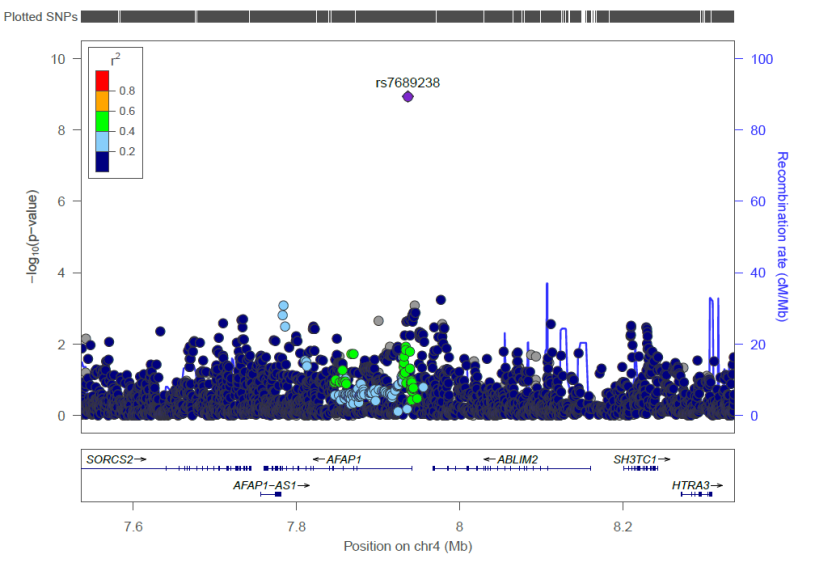

**Supplementary Figure 11: Regional plots of the replicated additional novel hits of the joint effect of long total sleep time on LDL cholesterol in the European meta-analyses**
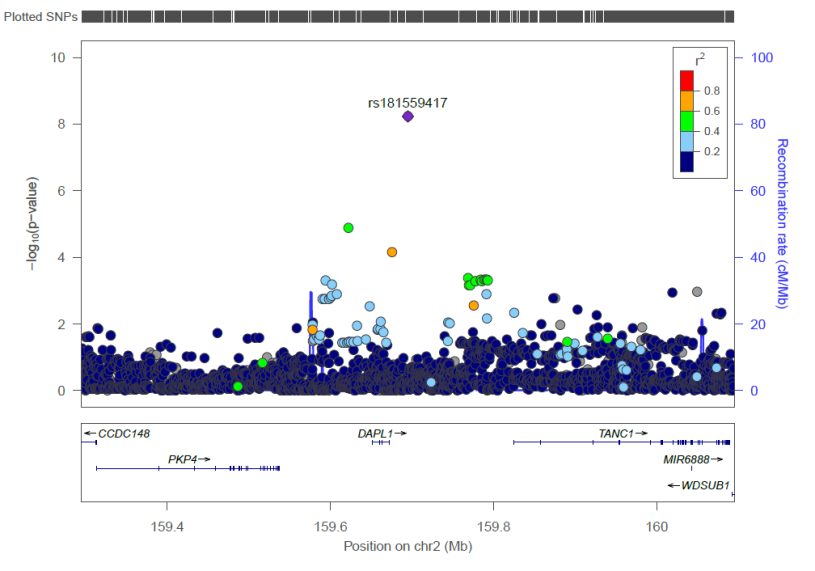

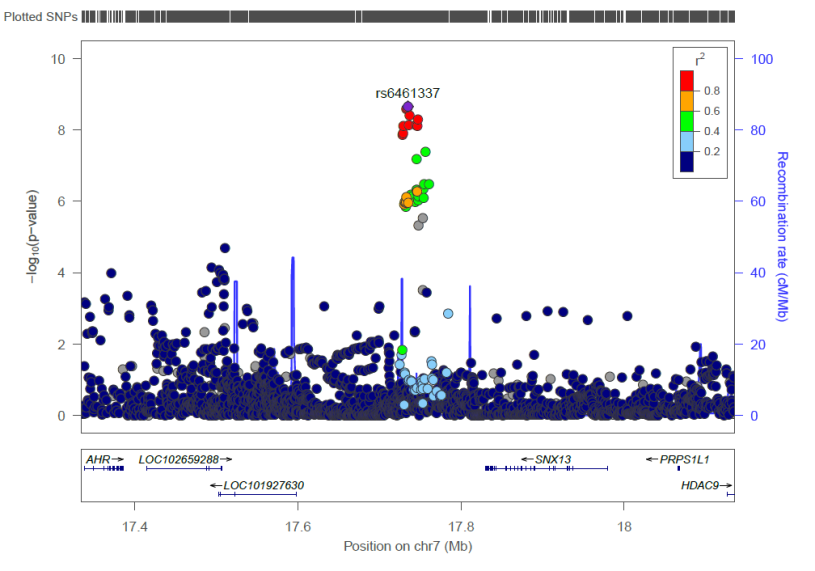

**Supplementary Figure 12: Regional plots of the replicated additional novel hits of the joint effect of long total sleep time on Triglycerides in the European meta-analyses**
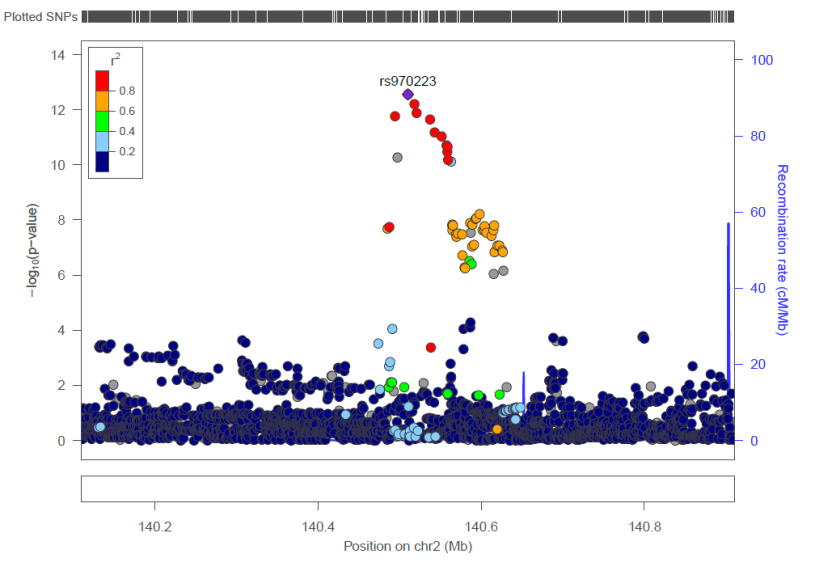

**Supplementary Figure 13: Regional plots of the replicated additional novel hits of the joint effect of short total sleep time on Triglycerides in the European meta-analyses**
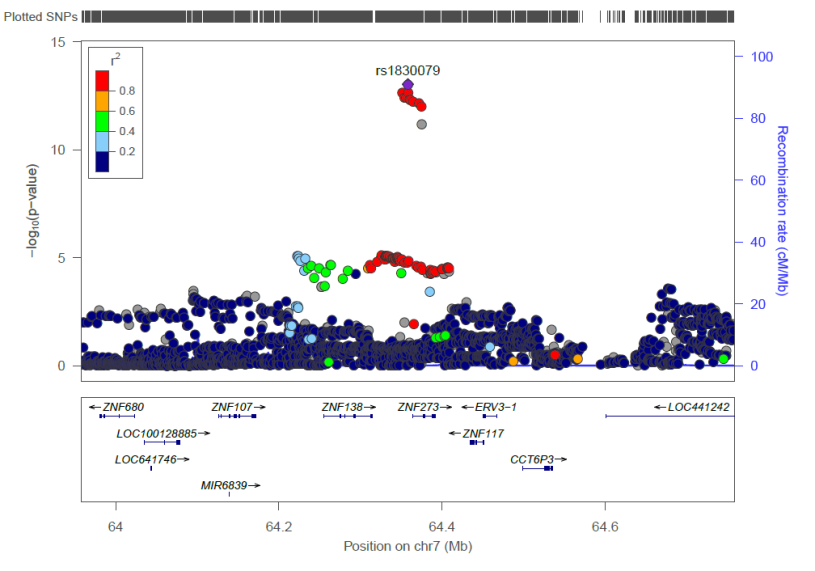

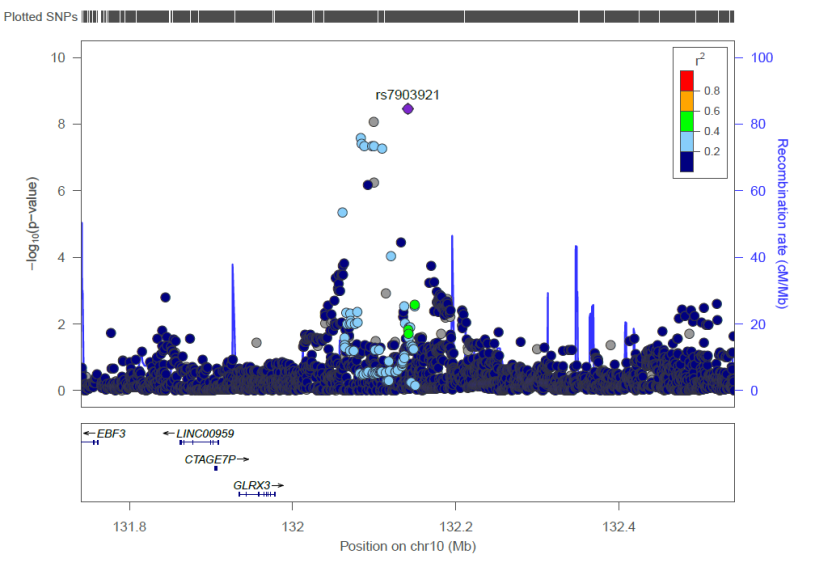
